# FLEXTAG: A Small and Self-renewable Protein Labeling System for Anti-fading Multi-color Super-resolution Imaging

**DOI:** 10.1101/2025.07.19.665678

**Authors:** Han Zhang, Yuan Yao, Xuye Wang, Yuanmin Zheng, Shaoqing Zhang, Yuan Tao, Haopeng Yang, Yinqi Wang, Mengde Liu, Marina Feric, Ruobo Zhou

## Abstract

Super-resolution fluorescence imaging enables visualization of subcellular structures and molecular interactions at the nanoscale, but its broader application has been hindered by long-standing limitations in current protein tagging systems, including rapid photobleaching, tag-induced artifacts, poor post-fixation labeling efficiency, and restricted multiplexing capability. Here, we present **FLEXTAG** (**<u>F</u>**luorescent **<u>L</u>**abeling for **<u>E</u>**xchangeable, **<u>X</u>**-resilient **<u>T</u>**agging in **<u>A</u>**dvanced **<u>G</u>**eneric Nanoscopy), a comprehensive protein labeling system comprising three orthogonal, ultrasmall (12–18 kDa), and self-renewable protein tags that collectively overcome these major limitations of existing tagging systems, enabling optimized multi-color super-resolution imaging. Through continuous exchange of organic fluorophores, FLEXTAG supports unprecedented durations of high-resolution imaging in both live and fixed cells with minimal photobleaching. It is compatible with major super-resolution modalities, such as SIM, STED, STORM, and PAINT, and is applicable to a wide range of subcellular targets. To further address fixation-induced labeling inefficiency and background fluorescence, we developed a novel protection-based fixation method and chemical blocking strategies that significantly preserve tag accessibility and enhance signal-to-noise ratio, improvements that are broadly applicable to other protein tagging systems. Altogether, FLEXTAG enables long-term tracking of dynamic behaviors and interactions of subcellular targets, as well as mapping of nanoscale protein organizations and cellular architecture, advancing both basic research and translational applications in cell biology.

## Introduction

Fluorescence microscopy has revolutionized biological research by enabling the studies of cellular structures and dynamics across scales from single molecules to live animals, due to its unique advantages compared to other microscopies, such as high image contrast, high molecular specificity, non-invasive multi-color labeling capability, and long-term live imaging capability. Over the past decade, super-resolution fluorescence imaging, also known as fluorescence nanoscopy, has been invented to overcome the light diffraction limit and improved the spatial resolution from ∼250 nm (this is ∼50-fold larger than the average size of a protein) to molecular scale (1–20 nm)^1,2^, allowing molecular-resolution fluorescence imaging. The most widely used nanoscopy techniques fall into two categories: 1) Patterned light illumination microscopy (PLIM), such as structured illumination microscopy (SIM) and stimulated emission depletion (STED) microscopy; 2) Single-molecule localization microscopy (SMLM), such as stochastic optical reconstruction microscopy (STORM)^3^, photoactivated localization microscopy (PALM) ^4^, and point accumulation for imaging in nanoscale topography (PAINT)^5^. These advancements in spatial resolution enable the visualization of previously unresolvable subcellular details, such as the organization of proteins within molecular complexes, nanoscale changes in cellular architecture, and dynamic molecular interactions, providing unprecedented mechanistic insights into biological processes.

The effectiveness of fluorescence nanoscopy relies on precise protein labeling strategies, which can be classified into three main approaches: 1) Immunofluorescence uses antibodies conjugated with organic dyes to recognize target proteins. While this method enables labeling of endogenous proteins, it is restricted to fixed-cell imaging. Additionally, only a small fraction of commercially available antibodies exhibit the high affinity and specificity required for nanoscopy, and their large size (∼150 kDa) limits access to densely packed cellular structures; 2) Fluorescent protein tags, such as EGFP, mEos3.2^6^, and mMaple3^7^, allow live-cell imaging but are often outperformed in brightness and photostability by organic dyes; 3) Self-labeling protein tags, such as Halo-^8^, CLIP-^9^, and SNAP-Tag^10^, provide site-specific labeling by reacting with small-molecule ligands conjugated to organic dyes. This approach offers improved photostability over fluorescent protein tags and is compatible with both live and fixed cells. However, even with organic dyes, photobleaching remains a major limitation, particularly in fluorescence nanoscopy, which requires prolonged high-intensity laser illumination. Additionally, in post-fixation labeling for fixed-cell imaging, commonly used fixatives such as paraformaldehyde (PFA) and glutaraldehyde (GA) can extensively crosslink proteins, reducing accessibility to dye-conjugated ligands and thereby lowering labeling efficiency of these self-labeling protein tags.

To mitigate photobleaching, self-renewable (i.e., exchangeable) protein tags have been recently developed, allowing continuous fluorescent dye replacement during imaging. Early efforts focused on fluorogenic GFP-like chromophores that reversibly bind to specific protein tags, such as pFAST^11^ and *blc* tags^12^. However, these chromophore-based renewable tags are incompatible with STED imaging due to their low brightness and photostability, and are also unsuitable for single-molecule localization microscopy (SMLM) such as STORM or PAINT: these chromophores lack the photoswitchability required for STORM, and under prolonged illumination in PAINT, they degrade into aldehyde derivatives that react with amines in the binding pocket^13^, preventing chromophore rebinding (Supplementary Fig. 1a-c). Additionally, these tags do not address the low post-fixation labeling efficiency caused by protein crosslinking during cell fixation. Another major challenge lies in expanding the spectral range of these tags for multi-color imaging, because chromophore binding is highly dependent on its precise chemical structure; even minor modifications to extend the color palette can disrupt tag recognition and reduce labeling efficiency. This limitation underscores the need to separate the fluorescent properties of the tag from its binding mechanism. More recently, HaloTag-based systems have been modified for reversible binding by engineering the HaloTag and its ligand to interact non-covalently^14^. This approach enables prolonged super-resolution imaging using both STED and PAINT. By linking nonfluorescent HaloTag ligands to various nanoscopy-compatible dyes, these systems (i.e., dye-conjugated HaloTag ligands) offer flexible color selection. However, the relatively large size of HaloTag (∼33 kDa) can interfere with the native localization and function of tagged proteins, as evidenced by mislocalization when tagging ATP5ME, COX8, YTHDC1, and the coronaviral envelope (E) protein^15^ (Supplementary Fig. 1d) and there are no similar self-renewable but orthogonal tags available that function across both PLIM- and SMLM-based techniques to enable three or more color super-resolution imaging. Furthermore, these existing tags do not overcome the challenge of low post-fixation labeling efficiency caused by protein crosslinking.

In this work, we present **<u>F</u>**luorescent **<u>L</u>**abeling for **<u>E</u>**xchangeable, **<u>X</u>**-resilient **<u>T</u>**agging in **<u>A</u>**dvanced **<u>G</u>**eneric Nanoscopy (**FLEXTAG**), a comprehensive framework that integrates three compact, self-renewable protein tags (FLEXTAG1, FLEXTAG2, and FLEXTAG3) with optimized labeling protocols to enable anti-fading multi-color super-resolution imaging. This FLEXTAG framework (Fig. 1) distinguishes itself from previous protein tagging and labeling methods in the following six key aspects: 1) The tag sizes are small (< 20 kDa) with minimal self-aggregation and tag-induced protein mislocalization issues, 2) The tags are self-renewable with minimal photobleaching, 3) It is compatible with both live and fixed cell imaging, 4) A protection-based fixation protocol has been developed to overcome low post-fixation labeling efficiency, 5) A chemical treatment and blocking protocol has been developed to minimize non-specific labeling, and 6) The tags are orthogonal and compatible with both PLIM-based and SMLM-based super-resolution imaging, providing a “one-for-all” solution, making it convenient to switch between different super-resolution imaging methods and enables three or more color super resolution nanoscopy. We envision this versatile super-resolution protein labeling system would allow for more flexible and detailed investigations on cellular processes, facilitating super-resolution imaging efforts to study molecular interactions, protein dynamics, and cellular functions.

**Fig. 1.**
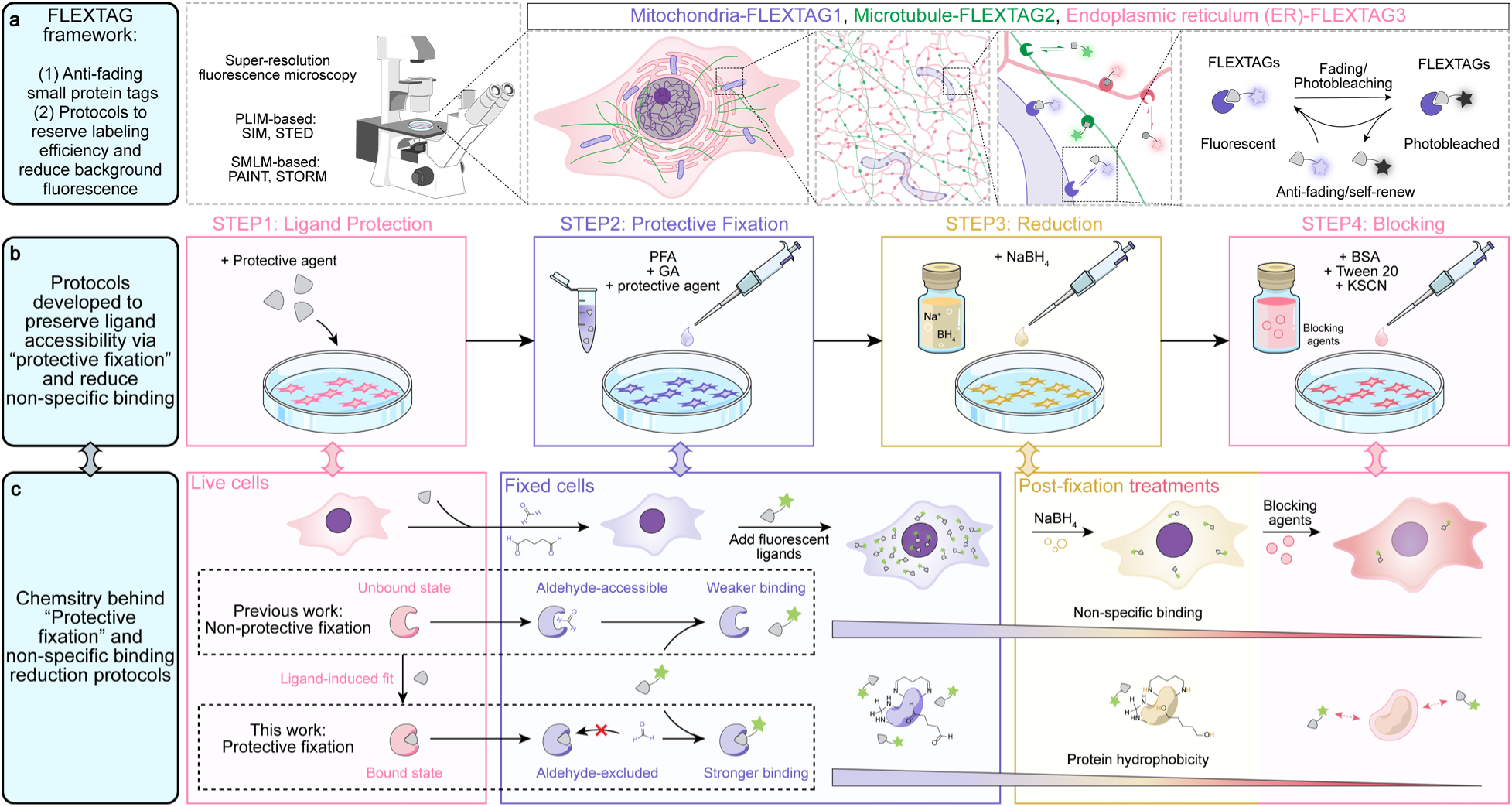
Overall scheme of the FLEXTAG framework. **a,** The developed FLEXTAG framework comprises three orthogonal, compact, and self-renewable (anti-fading) protein tags, along with optimized protocols designed to enhance labeling efficiency and reduce background fluorescence. Self-renewal allows photobleached fluorophores to be replaced by fresh fluorophores thereby restoring fluorescent signals. **b,** Schematic overview of the 4-step protocol designed to preserve ligand accessibility via “protective fixation” and reduce non-specific binding. Step 1: Preincubate cells expressing FLEXTAGs with a high-concentration of dye-free binding moieties of the FLEXTAG ligands, used as a protective agent. Step 2: Fix cells with an aldehyde-based fixative containing the same concentration as the protective agents. Step 3: Quench excess aldehydes with NaBH_4_. Step 4: Block fixed protein surfaces with a combination of BSA, Tween-20, and KSCN, identified as effective blocking agents. **c,** Chemical basis behind “Protective fixation” and nonspecific binding reduction strategies. Step 1: Preincubation with the dye-free binding moieties of the FLEXTAG ligand locks each tag into its ligand-bound conformation. Step 2: Occupation of the binding pockets shields reactive residues from crosslinking by aldehyde fixatives, thereby preserving ligand accessibility. Step 3: Residual aldehydes on protein surfaces are reduced to alcohols by NaBH_4_ decreasing surface hydrophobicity and reducing nonspecific binding of fluorescent ligands. Step 4: Blocking agents further modulate the microenvironment of fixed protein surfaces to suppress remaining nonspecific interactions.

## Results

### Development and characterization of FLEXTAG1

To develop new protein tags that address the limitations of current protein tagging systems, we sought to create a series of orthogonal, small, self-renewable protein tags (FLEXTAGs). These FLEXTAGs utilize synthetic fluorescent ligands, some of which are cell-permeable for live-cell imaging, while others are designed specifically for fixed-cell applications. Upon entering the live cell or accessing fixed cellular structures, the fluorescent ligands reversibly bind to and dissociate from non-fluorescent genetically encoded protein tags fused to target proteins, thereby minimizing photobleaching. To enable easy expansion for multi-color imaging, we decouple the fluorescent properties of the ligand from its binding to the tag by designing these fluorescent ligands with two components: a non-fluorescent part that specifically recognizes the protein tag and a nanoscopy-compatible fluorescent dye, connected through a polyethylene glycol (PEG) linker. The criteria used for the tag development are: 1) minimal aggregation and small size (< 20 kDa) to minimize tag-induced mislocalization and potential malfunction of the tagged protein, 2) sufficiently high labeling efficiency for compatibility with all major fluorescence nanoscopic methods, and 3) dynamic exchange between the fluorescent ligand and protein tag to support PAINT imaging and effectively mitigate photobleaching across all major fluorescence nanoscopy techniques.

Based on these criteria, we developed FLEXTAG1 from a mutant form of the second bromodomain of human BRD4, BRD4BD2^L55A^, a small protein degron tag (∼15 kDa, about half the size of the commonly used EGFP tag). Originally designed for tag-targeted proteolysis, this tag can be specifically recognized by the small molecule ligand ET-JQ1^16^. JQ1 is a small molecule inhibitor of bromodomains that bind to wild-type BRD4BD2 (BRD4BD2^WT^)^17^. To enhance binding specificity using the “bump-and-hole” strategy, an ethyl group was added to JQ1 to generate ET-JQ1 and the L55A mutation was introduced into BRD4BD2 to minimize off-target ligand binding to endogenous bromodomain proteins. Although this degron tag-ligand system has been successfully applied in tag-targeted proteolysis of human cells, it has not yet been further optimized for fluorescence imaging applications due to two challenges: 1) While ET-JQ1 shows minimal cross-reactivity with endogenous bromodomain proteins, as inferred from western blot data where ET-JQ1 induces significantly less degradation of endogenous bromodomains than of BRD4BD2^L55A^, this does not directly translate to imaging applications^16^. ET-JQ1 may still weakly and transiently bind to endogenous bromodomain proteins, generating unwanted fluorescence background; and 2) BRD4BD2 has a strong tendency to dimerize^18^, meaning BRD4BD2^L55A^ could promote aggregation of the tagged protein, potentially altering its subcellular localization and function and leading to experimental artifacts.

To evaluate these two potential issues, we synthesized tetramethylrhodamine (TMR)-conjugated ET-JQ1 and JQ1 (as a control). After labeling U2OS cells, with and without overexpressing a bromodomain protein BRD4, using ET-JQ1-TMR and JQ1-TMR (Fig. 2a), we observed that ET-JQ1-TMR showed no detectable binding to either overexpressed BRD4-GFP or endogenous bromodomain proteins (Fig. 2c), whereas JQ1-TMR exhibited strong binding to overexpressed BRD4-GFP and detectable binding to the nucleus-localized endogenous bromodomain proteins, indicating that the ET-JQ1, in contrast to JQ1, has negligible binding affinity to endogenous bromodomain proteins under imaging conditions (Fig. 2b-d). Next, we assessed the potential aggregation issue of BRD4BD2^L55A^. Indeed, after labeling cells overexpressing BRD4BD2^L55A^-tagged TOM20, a mitochondria-localized protein, with ET-JQ1-TMR, we observed severe aggregation of BRD4BD2^L55A^, drastically altering the expected mitochondrial morphology and distribution (Fig. 2f).

**Fig. 2.**
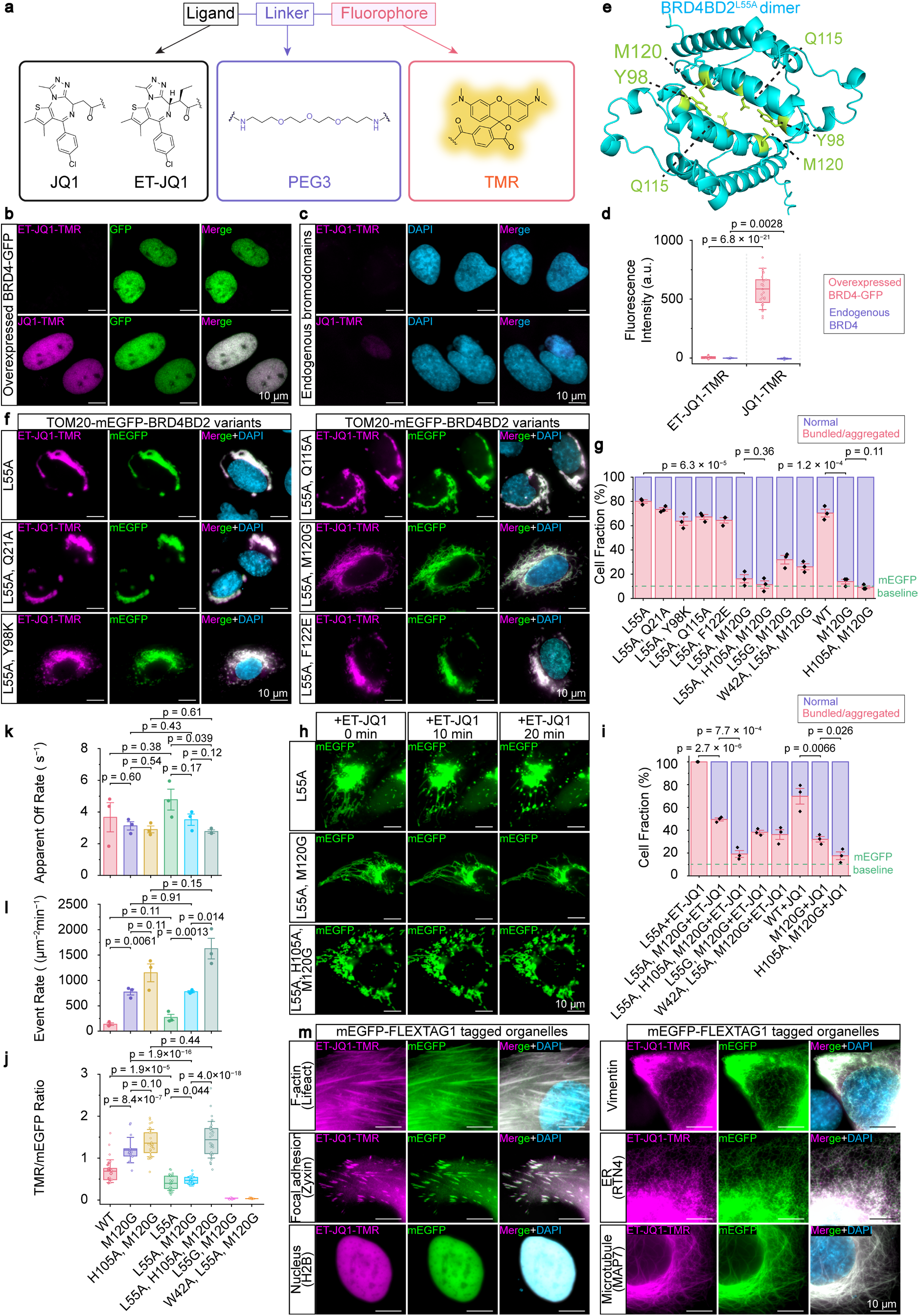
Development and characterization of FLEXTAG1. **a,** Chemical structures of ligand-linker-fluorophore conjugates targeting BRD4BD2^WT^ or BRD4BD2^L55A^. **b,** Representative epi-fluorescence images of GFP-tagged full length BRD4 (BRD4-GFP), labeled with 1 µM ET-JQ1-TMR or JQ1-TMR. Scale bars: 10 μm. **c,** Representative epi-fluorescence images of endogenous bromodomain proteins, labeled by 1 µM ET-JQ1-TMR or JQ1-TMR and 1 µg/mL DAPI. Scale bars: 10 μm. **d,** Quantification of ET-JQ1-TMR or JQ1-TMR fluorescence intensity in cells overexpressing BRD4-GFP or endogenous bromodomain proteins. Each data point represents the average TMR signal within a GFP- or DAPI-masked nuclear region from a single field of view (FOV). *n* = 13–25 FOVs per condition were examined. **e,** ColabFold-predicted dimer interface of BRD4BD2^L55A^. Key inter-protein interactions are highlighted in green, including Y98-M120 and Q115-Q115. **f,** Representative epi-fluorescence images of TOM20-mEGFP-BRD4BD2^L55A^ variants. All mutations retain binding capability to ET-JQ1-TMR. Scale bars: 10 μm. **g,** Quantification of the fraction of cells overexpressing TOM20-mEGFP-BRD4BD2 variants showing normal versus bundled/aggregated mitochondrial morphology. Cells overexpressing TOM20-mEGFP, which does not dimerize, served as the monomeric baseline control (dashed green line). *n* = 3 biological replicates; 60–100 cells were examined for each BRD4BD2 variant. **h,** Representative epi-fluorescence images of mitochondrial morphology in cells overexpressing TOM20-mEGFP-BRD4BD2 variants at 0, 10, and 20 minutes after the addition of 100 μM ET-JQ1. Scale bars: 10 μm. **i,** Same as **g**, except the mitochondrial morphology was assessed 1 hour after the addition of 100 μM ET-JQ1. *n* = 3 biological replicates, 60–100 cells were examined for each BRD4BD2 variant. **j,** Quantification of the TMR/mEGFP fluorescence ratio following labeling of cells expressing TOM20-mEGFP-BRD4BD2 variants with 1 µM ET-JQ1-TMR. Each data point represents the average TMR/mEGFP ratio of mEGFP-masked regions in a single FOV. *n* = 13–33 FOVs per condition were examined. **k,** Quantification of apparent off-rates for 1 nM ET-JQ1 or JQ1-TMR binding to TOM20-mEGFP-BRD4BD2 variants in fixed cells, based on single-molecule localization microscopy (SMLM) movies (10,000 frames, 10 Hz). *n* = 3 biological replicates; 3 SMLM movies were examined for each BRD4BD2 variant. **l,** Same as **k**, but quantifying binding event rates. **m,** Representative epi-fluorescence images of BRD4BD2^L55A, H105A, M120G^ (FLEXTAG1)-tagged cellular structures. Samples were labeled with 1 µM ET-JQ1-TMR or JQ1-TMR and 1µg/mL DAPI. Scale bars: 10 μm. Data are presented as mean ± s.e.m. Boxplots show the mean and boundaries (first and third quartiles); whiskers denote s.d. *p*-values calculated with two-sided unpaired Student’s *t*-test.

To minimize the aggregation of BRD4BD2^L55A^, we used ColabFold^19^, an artificial intelligence system that predicts 3D structures of proteins and protein interactions from their amino acid sequences, to predict its dimer interface and identify the key interfacial interactions (≤ 3.5 Å bond distances), primarily between the Q115 residues of each monomer and between Y98 of one monomer and M120 of the other (Fig. 2e, Supplementary Fig. 2a). This predicted dimer interface closely resembles the intermolecular contacts observed in the crystal lattice formed by co-crystallizing BRD4BD1 and BRD4BD2^18^, In addition, a previous study identified several mutations that reduce homodimer formation of BRD2BD1^20^, a bromodomain that shares a nearly identical structure with BRD4BD2 (Supplementary Fig. 2b). Based on the key interfacial residues identified from the predicted dimer interface and the previous mutations known to disrupt BRD2BD1 dimerization, we selected five mutations including Q21A, Y98K, Q115A, M120G, and F122E, to test their ability to disrupt BRD4BD2 dimerization while retaining ET-JQ1 binding. We introduced each of these mutations into the BRD4BD2^L55A^ background and quantified the percentage of cells exhibiting abnormal (i.e., either bundled or aggregated) mitochondrial morphology when overexpressing TOM20 tagged with these BRD4BD2 variants (Hereafter referred to as the mitochondria morphology assay, which is a quantitative method for assessing the extent of protein tag aggregation; Supplementary Fig. 3). Among the tested mutations, Q21A, Y98K, Q115A, and F122E failed to reduce BRD4BD2^L55A^ dimerization. In contrast, M120G was the only mutation that effectively minimized dimerization-induced mitochondrial bundling or aggregation, reducing the percentage of cells with abnormal mitochondrial morphology from 80.0 ± 1.4% to 16.2 ± 3.4%, while still maintaining ET-JQ1 binding. Furthermore, M120G not only significantly reduced BRD4BD2^L55A^ dimerization but also disrupted BRD4BD2^WT^ dimerization, confirming the essential role of M120 in BRD4BD2 dimerization (Fig. 2g; Supplementary Fig. 4).

As in our FLEXTAG system, we developed protection-based fixation method (described in a later section) to preserve tag accessibility after cell fixation and address low post-fixation labeling efficiency, where it requires adding a relatively high concentration (∼ 100 μM) of the ligand (ET-JQ1 for BRD4BD2^L55A, M120G^ or JQ1 for BRD4BD2 ^M120G^) during the cell fixation process to protect residues surrounding the binding pocket from fixation and hence prevent loss of ligand binding after fixation. Therefore, we performed additional tests to see if the tag remains prone to aggregation in the presence of its ligand ET-JQ1. Unfortunately, when incubating cells overexpressing TOM20 tagged with BRD4BD2^L55A^ or BRD4BD2^L55A, M120G^ with 100 μM ET-JQ1, both constructs exhibited strong mitochondrial bundling and aggregation (Fig. 2h; Supplementary Fig. 4). The percentage of cells overexpressing BRD4BD2^L55A, M120G^ with abnormal mitochondrial morphology increased from 16.2 ± 3.4% back to 100.0 ± 0.0%. Similarly, 100 μM JQ1-TMR increased the percentage of cells overexpressing BRD4BD2^M120G^ with abnormal mitochondrial morphology from 13.9 ± 1.9% to 32.1 ± 2.4%, although this ligand-induced aggregation was less pronounced than in BRD4BD2^L55A, M120G^, indicating that ligand binding mediates a new dimer interface near the ligand binding pocket for both protein tags (Fig. 2i; Supplementary Fig. 4). To investigate this newly induced dimerization, we used ColabFold to predict the dimer of BRD4BD2^L55A, M120G^. Although ColabFold does not support structure prediction of ligand-bound dimers, a new dimer interface, predicted in the absence of ligand, emerged near the ligand-binding pocket, with the key interfacial interactions between H105 residues of each monomer and between W42 of one monomer and A52 of the other (Supplementary Fig. 2c). Based on the predicted new dimer interface, and that L55A exhibited more severe ligand-induced aggregation, we designed three variants to disrupt this new dimer interface: BRD4BD2^L55G, M120G^, BRD4BD2^W42A, L55A, M120G^, and BRD4BD2^L55A, H105A, M120G^. Using the mitochondrial morphology assay (Supplementary Fig. 3), we found that only BRD4BD2^L55A, H105A, M120G^ disrupted the ligand-induced mitochondrial aggregation (Fig. 2g, 2i, Supplementary Fig. 4), while the other two mutants abolished ET-JQ1-TMR binding (Supplementary Fig. 2d).

Next, following our criteria for FLEXTAG aforementioned, we evaluated the labeling efficiencies and the dynamic exchange rates between the fluorescent ligand and protein tag across several BRD4BD2 variants. To this end, we fused each BRD4BD2 variant, along with a fluorescent protein tag (mEGFP, used as an expression level reference for normalization), to the mitochondria-localized protein TOM20 (TOM20-mEGFP-BRD4BD2 variants). After labeling U2OS cells overexpressing a variant of TOM20-mEGFP- BRD4BD2 with JQ1-TMR (for variants without L55A mutation) or ET-JQ1-TMR (for variants with L55A mutation), we determined the fluorescence intensity ratios of TMR to mEGFP, which quantitatively represent the labeling efficiencies of each BRD4BD2 variant. Among the variants tested, BRD4BD2^L55A, H105A, M120G^ exhibited the highest TMR/EGFP fluorescence ratio, indicating superior labeling efficiency (Fig. 2j; Supplementary Fig. 2d). Using PAINT imaging, we further quantified two dynamic exchange parameters in the presence of 1 nM ligand: 1) Event rate, defined as the average number of fluorescent ligand binding events per unit imaging area per unit time; 2) Apparent off-rate, defined as the average rate of disappearance of the single molecule fluorescent spot. While these rates do not precisely reflect the true ligand binding kinetics, since the apparent off-rate may be overestimated due to TMR photobleaching, and the event rate differs from the true on-rate as it is measured per imaging area rather than per tag, they still provide a practical assessment of PAINT compatibility. When these kinetic properties align with PAINT requirements, they often also help mitigate photobleaching in other nanoscopic techniques such as STORM, SIM, and STED. We found while the apparent off-rate of BRD4BD2^L55A, H105A, M120G^ was slightly lower than other BRD4BD2 variants (Fig. 2k), its event rate was significantly higher (Fig. 2l), making it the most optimized for PAINT imaging.

Finally, we validated the performance of BRD4BD2^L55A, H105A, M120G^ for tagging diverse proteins, localized to actin filaments, focal adhesions, the nucleus, vimentin, ER, and microtubules, respectively. The observed fluorescence patterns closely matched the expected localization and morphology of the targeted cellular structures (Fig. 2m). We therefore designated BRD4BD2^L55A, H105A, M120G^ as our developed FLEXTAG1.

### Development and characterization of FLEXTAG2

We developed FLEXTAG2 from *E. coli* dihydrofolate reductase (eDHFR), a previously reported small (∼18 kDa) protein imaging tag that can be specifically recognized by its specific small molecule inhibitor trimethoprim (TMP)^21^ (Fig. 3a). Although eDHFR and dye-conjugated TMP have been successfully used for protein imaging in live cells, this system remains largely incompatible with cell fixation, and its fluorescence signal is relatively weak due to low labeling efficiency^22,23^ (defined as the percentage of protein tags in the fluorescent state at any given time). This inefficiency stems from the relatively low affinity of the non-covalent interaction between eDHFR and dye-conjugated TMP, leaving a large fraction of eDHFR tags in a dark, TMP-unbound state at any given time, limiting its applications in fluorescence nanoscopy where high fluorescence signal is often required. To address this, eDHFR mutations (e.g., L28C) and TMP variants have been developed to enable covalent bonding between eDHFR and TMP^22,23^, However, this covalent interaction eliminates eDHFR’s reversible binding and dissociation properties, which are crucial for mitigating photobleaching. Therefore, to achieve higher labeling efficiency while retaining its self-renewable nature of the tag, it is crucial to engineer eDHFR to enhance the binding rate constant (*k*_on_) of TMP-dye binding to eDHFR.

**Fig. 3.**
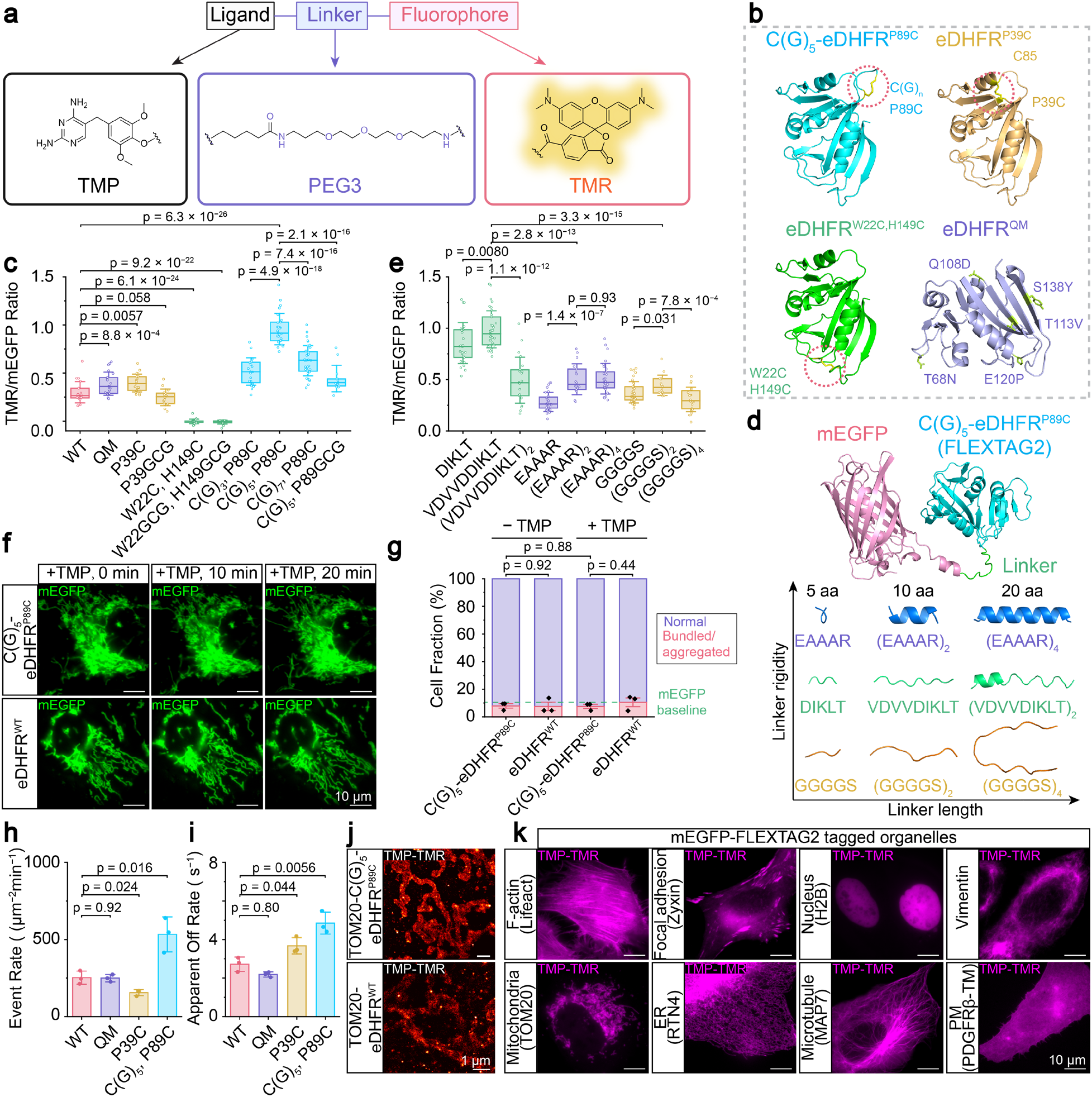
Development and characterization of FLEXTAG2. **a,** Chemical structures of ligand-linker-fluorophore conjugates targeting eDHFR. **b,** ColabFold-predicted structures of eDHFR variants. Designed intramolecular disulfide bonds (yellow) in C(G)_5_-eDHFR^P89C^, eDHFR^P39C^, and eDHFR^W22C, H149C^ are highlighted with red circles. Thermally stabilizing mutations in eDHFR^QM^ are shown in green. **c,** Quantification of the TMR/mEGFP fluorescence ratio in cells overexpressing TOM20-mEGFP-eDHFR variants labeled with 1 µM TMP-TMR. Each data point represents the average TMR/mEGFP ratio of mEGFP-masked regions in a single FOV. *n* = 22–46 FOVs per condition were examined. **d,** ColabFold-predicted structures of mEGFP-(VDVVDDIKLT)-C(G)_5_-eDHFR^P89C^ (FLEXTAG2) and designs of linkers inserted between mEGFP and eDHFR variants. Linkers of varying lengths (5, 10, or 20 amino acids) and rigidity (rigid, moderately flexible, or flexible, based on the predicted confidence of stable α-helix formation) were screened. **e,** Same as **c** but quantifying cells overexpressing TOM20-mEGFP-(designed linkers)-C(G)_5_-eDHFR^P89C^. *n* = 15–38 FOVs per condition were examined. **f,** Representative epi-fluorescence images showing mitochondrial morphology in cells overexpressing TOM20-mEGFP-C(G)_5_-eDHFR^P89C^ or TOM20-mEGFP-eDHFR^WT^ at 0, 10, and 20 minutes after the addition of 100 μM TMP. Scale bars: 10 μm. **g,** Quantification of the fraction of cells overexpressing TOM20-mEGFP-C(G)_5_-eDHFR^P89C^ or TOM20-mEGFP-eDHFR^WT^ showing normal versus bundled/aggregated mitochondrial morphology, in the absence or 1 hour after the addition of 100 μM TMP. The baseline fraction of cells overexpressing TOM20-mEGFP that exhibit bundled/aggregated mitochondria is indicated by the green dashed line. *n* = 3 biological replicates; 60–100 cells were examined for each eDHFR variant. **h,** Quantification of binding event rates for 1 nM TMP-TMR binding to TOM20-mEGFP-eDHFR variants in fixed cells, based on SMLM movies (10,000 frames, 10 Hz). Each data point represents the average number of localizations per μm² per minute for one cell. Data are presented as mean ± s.e.m. *n* = 3 biological replicates; 3 SMLM movies were examined for each eDHFR variant. p-values calculated with two-sided unpaired Student’s t-test. **i,** Same as **h** but for quantifying apparent off-rates. **j,** Representative PAINT images of TOM20-mEGFP-C(G)_5_-eDHFR^P89C^ and TOM20-mEGFP-eDHFR^WT^, imaged with 1 nM TMP-TMR in fixed cells. Scale bars: 1 μm. **k,** Representative epi-fluorescence images of C(G)_5_-eDHFR^P89C^ (FLEXTAG2)-tagged cellular structures, labeled with 1 µM TMP-TMR. Scale bars: 10 μm. Data are presented as mean ± s.e.m. Boxplots show the mean and boundaries (first and third quartiles); whiskers denote s.d. *p*-values calculated with two-sided unpaired Student’s *t*-test.

Previous studies have reported that eDHFR has a relatively low thermal denaturation temperature *T*_m_ of 43°C^24^, and exhibits low mechanical stability, with an average unfolding force of ∼27 pN, much lower than that of most globular proteins^25^. These characteristics led us to hypothesize that its inherent instability partially contributes to insufficient labeling. This instability may result in a fraction of eDHFR molecules being in a conformational state in which TMP ligands cannot bind, thereby reducing fluorescence intensity. To test this hypothesis, we identified and designed ten eDHFR variants with potentially improved stability, categorized into four groups: 1) eDHFR^T68N, Q108D, T113V, E120P, S138Y^ (eDHFR with quintuple mutations, termed eDHFR^QM^), which has been reported to exhibit a higher *T*_m_ than wild-type eDHFR (eDHFR^WT^)^26^; 2) eDHFR^P39C^ and eDHFR^P39GCG^. eDHFR^P39C^, a mutant previously reported to have a higher midpoint denaturant concentration^27^, features a Proline-39 to Cysteine (P39C) substitution, enabling disulfide bond formation between P39C and the cysteine-85 to stabilize the folded eDHFR structure. In eDHFR^P39GCG^, P39 is substituted by GCG instead of C to allow additional flexibility for disulfide formation. 3) eDHFR^W22C, H149C^ and eDHFR^W22GCG, H149GCG^, where two residues in proximity, at positions 22 and 149, are substituted by either a C or GCG to enable disulfide bond formation between them; 4) C(G)_n_-eDHFR^P89C^ and C(G)_5_-eDHFR^P89GCG^ where n = 3, 5, 7. P89 is substituted by either C or GCG, and an additional amino acid sequence C(G)_n_ is introduced at the N-terminus of eDHFR to promote disulfide bond formation between P89C and C(G)_n_. Using ColabFold, we confirmed that all designed eDHFR variants, except for eDHFR^P39GCG^, can form the intended disulfide bonds (Fig. 3b; Supplementary Fig. 5a-d).

To compare the labeling efficiencies among the ten eDHFR variants, we utilized the similar double tagged TOM20 constructs (TOM20-mEGFP-eDHFR variants) as used for BRD4BD2 variants. After labeling U2OS cells overexpressing TOM20-mEGFP-eDHFR with TMR-conjugated TMP (TMP-TMR) (Fig. 3a), we determined the fluorescence intensity ratios of TMR to mEGFP, which quantitatively represents the labeling efficiencies of each eDHFR variant. The TMR/mEGFP fluorescence ratios for eDHFR^W22C, H149C^ and eDHFR^W22GCG, H149GCG^ dropped to nearly zero, indicating that mutations at positions 22 and 149 resulted in a near complete loss of TMP-TMR binding affinity. In contrast, all other eDHFR variants retained their TMP-TMR binding. Among these, eDHFR^P39C^ and eDHFR^QM^ showed a 30-40% increase in labeling efficiency, as indicated by the TMR/mEGFP ratio, compared to eDHFR^WT^, while eDHFR^P39GCG^ exhibited labeling efficiency comparable to eDHFR^WT^, consistent with ColabFold predictions indicating its inability to form a disulfide bond. Notably, C(G)_n_-eDHFR^P89C^ variants exhibited greater labeling efficiency, with the highest enhancement (∼3.3-fold) observed at the poly-glycine linker length of n = 5. Introducing additional glycine residues on both sides of cysteine (GCG) to increase flexibility decreased the labeling efficiency compared to a single cysteine mutation. Therefore, we selected C(G)5-eDHFR^P89C^ for further optimization (Fig. 3c, Supplementary Fig. 5e).

Having identified that the length of poly-glycine linker between the N-terminal cysteine and the N-terminus of eDHFR^P89C^ affects labeling efficiency, we further aimed to optimize the linker length and sequence on the other side of the N-terminal cysteine (i.e., the linker between C(G)_5_-eDHFR^P89C^ and the target protein). We designed the linkers of varying lengths (5, 10, and 20 amino acids) with different flexibilities, predicted using ColabFold (Fig. 3d). By quantifying TMR/mEGFP ratios, we found that the 10-amino-acid linkers yielded the highest labeling efficiency of C(G)_5_-eDHFR^P89C^ compared to 5- or 20-amino-acid linkers. Among the tested sequences, the moderately flexible linkers resulted in the highest labeling efficiency compared to the flexible (i.e., GGGGS-based) or the rigid linkers (i.e., EAAAR-based) linkers. In contrast, the labeling efficiency of eDHFR^WT^ was unaffected by linker length and flexibility, suggesting that the observed linker-dependent effect in C(G)_5_-eDHFR^P89C^ is likely due to its influence on disulfide bond formation (Fig. 3e; Supplementary Fig. 6).

Next, we assessed the potential aggregation tendency of C(G)_5_-eDHFR^P89C^. Although eDHFR^WT^ is a monomer^28^, we needed to confirm that C(G)_5_-eDHFR^P89C^ mutation does not increase aggregation. Using the mitochondrial morphology assay, we found the percentage of cells overexpressing TOM20-mEGFP-eDHFR^P89C^ exhibiting aggregation was comparable to those overexpressing TOM20-mEGFP-eDHFR^WT^. Additionally, the presence of 100 μM TMP, the concentration used in our protection-based fixation method, did not noticeably increase aggregation (Fig. 3f,g; Supplementary Fig. 7). To test whether the ligand binding kinetics of C(G)₅-eDHFR^P89C^ supports PAINT imaging, which also allows effectively mitigate photobleaching across all major fluorescence nanoscopic methods, we quantified event rate and apparent off-rate, as done for BRD4BD2 variants, at 1 nM TMP-TMR. C(G)₅-eDHFR^P89C^ exhibited the highest event rate (Fig. 3h) and apparent off-rate (Fig. 3i) in PAINT imaging, indicating that TMP binds to and dissociates from C(G)₅-eDHFR^P89C^ more rapidly than other eDHFR variants. This faster ligand exchange makes C(G)₅-eDHFR^P89C^ particularly well-suited for higher speed PAINT imaging. Indeed, PAINT images C(G)₅-eDHFR^P89C^ outperformed those obtained using eDHFR^WT^ (Fig. 3j).

Finally, we evaluated the performance of C(G)₅-eDHFR^P89C^ for tagging proteins localized to diverse cell organelles, as done for FLEXTAG1. The obtained fluorescence images showed expected subcellular localization and morphology of the targeted cellular structures (Fig. 3k). We therefore designated C(G)₅-eDHFR^P89C^ as FLEXTAG2.

### Development and characterization of FLEXTAG3

We developed FLEXTAG3 from a mutant form of human FKBP, FKBP^F36V^, a previously reported small (∼12 kDa) protein tag used for tag-targeted proteolysis, chemical induction of protein interaction, and fluorescence imaging^29–31^. Based on the “bump-and-hole” strategy, the F36V mutation and its selective ligand SLF’ (also known as AP1867) were developed to prevent undesired ligand binding to endogenous FKBPs. The high binding affinity of SLF’ to FKBP^F36V^ (*K_D_* = 0.094 nM^30^) results in extremely slow exchange dynamics, making it incompatible with PAINT imaging and ineffective for mitigating photobleaching across major fluorescence nanoscopic methods. To enhance ligand exchange kinetics, a recent study identified the FKBP^F36L^ mutant^32^. However, we found that overexpression of FKBP^F36L^-tagged TOM20 in cells led to severe mitochondrial aggregation, with 74.3 ± 2.4% of cells exhibiting abnormal mitochondrial morphology (Fig. 4e,f; Supplementary Fig. 10), Structure prediction using ColabFold revealed that the FKBP^F36L^ dimer interface is indeed mediated primarily through L36, with L36 of one monomer interacting with K34 and I90 of the other, along with additional interfacial contacts (Supplementary Fig. 8a).

**Fig. 4.**
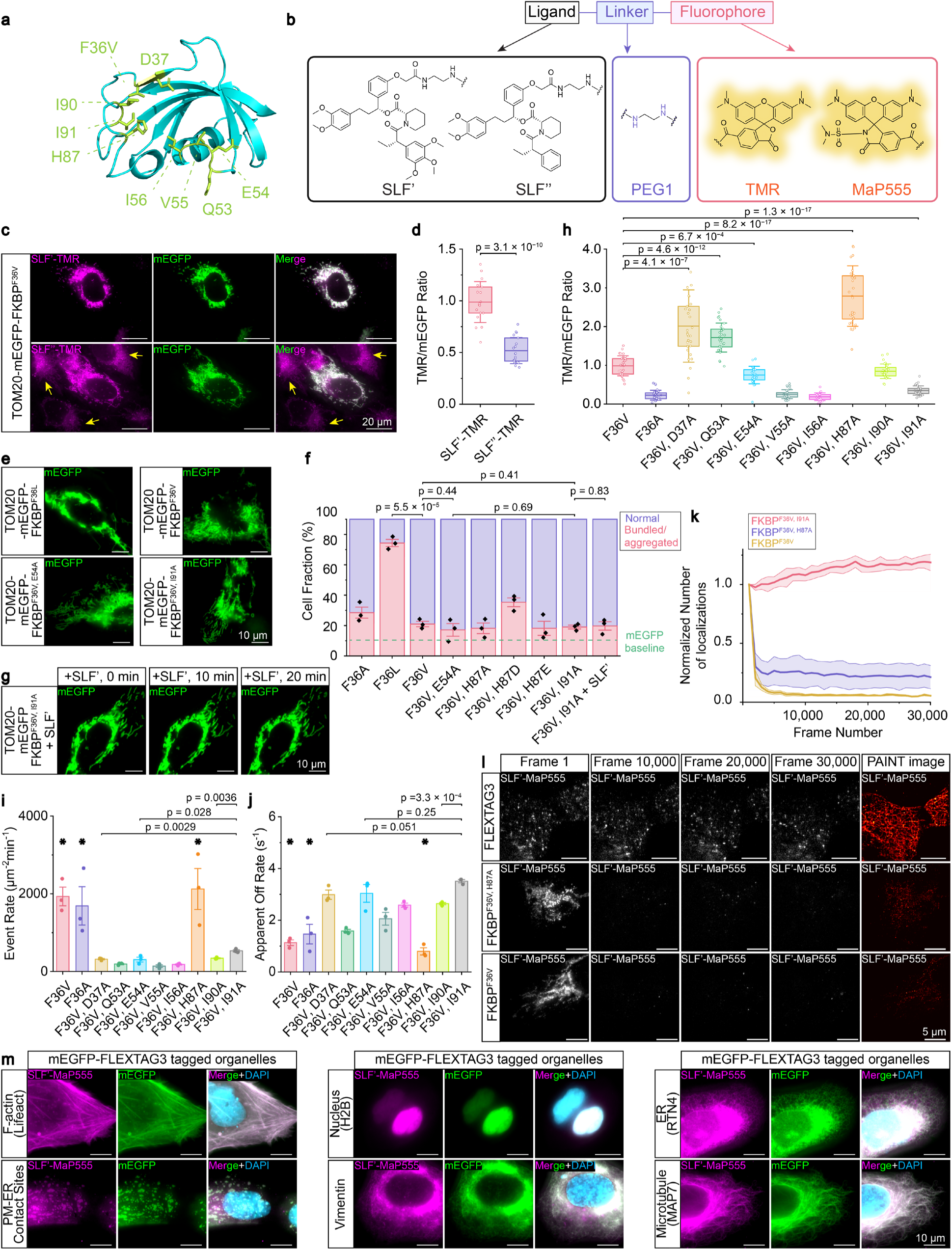
Development and characterization of FLEXTAG3. **a,** ColabFold-predicted structure of FKBP^F36V^. Amino acids near the ligand binding pocket potentially affecting exchange kinetics are highlighted in green, including D37, Q53, E54, V55, I56, H87, I90, and I91. **b,** Chemical structures of ligand-linker-fluorophore conjugates targeting FKBP variants. **c,** Representative epi-fluorescence images of TOM20-mEGFP-FKBP^F36V^labeled with 1 µM SLF’-TMR or SLF’’-TMR. SLF’-TMR shows high selectivity for FKBP^F36V^, whereas SLF’’-TMR exhibits fluorescence signals in cells lacking FKBP^F36V^ expression. Yellow arrowheads indicate non-specific binding in untransfected cells. Scale bars: 20 μm. **d,** Quantification of TMR/mEGFP fluorescence ratios in cells overexpressing FKBP^F36V^ labeled with 1 µM SLF’-TMR or SLF’’-TMR. Each data point represents the average TMR/mEGFP ratio of mEGFP-masked regions in a single FOV. *n* = 18 for SLF’-TMR; *n* = 19 for SLF’’-TMR. **e,** Representative images of mitochondrial morphology in cells overexpressing TOM20-mEGFP-FKBP variants. Scale bars: 10 μm. **f,** Quantification of the fraction of cells overexpressing TOM20-mEGFP-FKBP variants showing normal versus bundled/aggregated mitochondria. The baseline fraction of cells overexpressing TOM20-mEGFP that exhibit bundled/aggregated mitochondria is indicated by the green dashed line. *n* = 3 biological replicates; 40–100 cells were examined for each FKBP variant. **g,** Representative epi-fluorescence images showing mitochondrial morphology in cells overexpressing TOM20-mEGFP-FKBP^F36V, I91A^ at 0, 10, and 20 minutes after the addition of 100 μM SLF’. Scale bars: 10 μm. **h,** Quantification of the TMR/mEGFP fluorescence ratio in cells overexpressing TOM20-mEGFP-FKBP variants labeled with 1 µM SLF’-TMR. Each data point represents the average TMR/mEGFP ratio of mEGFP-masked region in a single FOV. 25 ≤ *n* ≤ 30. **i,** Quantification of binding event rates for 100 pM SLF’-MaP555 binding to TOM20-mEGFP-FKBP variants in fixed cells, based on SMLM movies (10,000 frames, 10 Hz). Mutations with prolonged binding durations, where photobleaching often precedes dissociation, potentially leading to inaccurate kinetic measurements, are marked with asterisks. Each data point represents the average number of localizations per μm² per minute for one cell. *n* = 3 biological replicates; 3 SMLM movies were examined for each FKBP variant. Data are presented as mean ± s.e.m.. **j,** Same as **i**, except for quantifying apparent off rates. **k,** Quantification of the survival rate of the single-molecule localizations during 30,000-frame PAINT imaging of 1 nM SLF’-MaP555 binding to FKBP^F36V, I91A^, FKBP^F36V, H87A^, and FKBP^F36V^ in fixed cells. Localization numbers per frame were normalized to the first frame. *n* = 3 biological replicates, 3 SMLM movies were examined for each FKBP variant. Data are presented as mean ± s.e.m. **l,** Representative images at frames 1, 10,000, 20,000, 30,000, and reconstructed PAINT images of FKBP^F36V, I91A^ (FLEXTAG3), FKBP^F36V, H87A^, and FKBP^F36V^. Scale bars: 5 μm. **m,** Representative epi-fluorescence images of FKBP^F36V, I91A^ (FLEXTAG3)-tagged cellular structures. Samples were labeled with 1 µM SLF’-MaP555 and 1µg/mL DAPI. Scale bars: 10 μm. Boxplots show the mean and boundaries (first and third quartiles); whiskers denote s.d. *p*-values calculated with two-sided unpaired Student’s *t*-test.

We next sought to identify mutations that could further disrupt the dimerization tendency of FKBP^F36V^. ColabFold predictions showed a distinct dimer interface for FKBP^F36V^ compared to FKBP^F36L^, with interfacial interactions (bond distances ≤ 3.5 Å) observed between: 1) the side chain of T85 and the backbone of Q53, E54, and V55, 2) the side chains of F46 and P88, and 3) the side chains of Y82 and H87 (Supplementary Fig. 8b). Unfortunately, this dimer interface lies near the ligand binding pocket. Despite multiple prediction rounds, we found that attempts to disrupt dimerization often resulted in structural disruptions to the binding pocket. Specifically, introducing alanine substitutions or rationally designed mutations at all interfacial residues led to poor folding around the binding pocket, while failing to abolish dimer formation—new interfacial interactions formed, resulting in a similar overall dimer structure (Supplementary Fig. 8c,e). To mitigate this, we narrowed our focus to only the strongest interfacial interactions (bond distances ≤ 3.0 Å), identifying E54–T85 and Y82–H87 as key interfacial contacts (Supplementary Fig. 8d). We tested four mutations on the FKBP^F36V^ background: E54A, H87A, H87D, and H87E. However, mitochondrial morphology assays showed that none significantly further reduce aggregation, with 17.2 ± 4.2%, 18.2 ± 3.5%, 35.3 ± 2.9%, and 18.2 ± 4.6% of cells displaying abnormal (i.e. aggregated or bundled) mitochondria, respectively (Fig. 4f). The double mutant E54A/H87E also failed to further disrupt dimer formation, based on ColabFold predictions (Supplementary Fig. 8e). Additionally, deletion of the Q53–V55 segment, intended to eliminate backbone-mediated interactions, resulted in severe protein misfolding (Supplementary Fig. 8e). Therefore, despite extensive experimental and computational efforts, further reducing FKBP dimerization tendency remains a challenge. We therefore chose to proceed with FKBP^F36V^, as it exhibits a tolerable aggregation profile, with ∼20% of cells displaying mitochondrial bundling or aggregation.

To increase ligand binding kinetics to FKBP^F36V^, we sought to evaluate alternative ligands and identify new mutations with the FKBP^F36V^ scaffold. SLF”, a derivative of SLF’ with the three methoxy groups removed, was reported to exhibit weaker affinity toward FKBP^F36V^ than SLF’. To compare the labeling efficiencies of SLF’ and SLF”, we labeled U2OS cells overexpressing double tagged TOM20 constructs (TOM20-mEGFP- FKBP^F36V^) with TMR-conjugated SLF’ (SLF’-TMR) and SLF” (SLF”-TMR) (Fig. 4b). Although SLF”-TMR exhibited reduced labeling efficiency to FKBP^F36V^, considerable fluorescent signal in the cytosol of untransfected cells suggested off-target binding of SLF”-TMR to endogenous FKBPs, making SLF’ the preferred ligand for imaging applications (Fig. 4c,d).

To further enhance SLF’ binding kinetics while avoiding aggregation issues associated with FKBP^F36L^, we engineered eight new single-point mutations near the SLF’ binding pocket in FKBP^F36V^ background: D37A, Q53A, E54A, V55A, I56A, H87A, I90A, and I91A. We assessed these variants for labeling efficiency using the TMR/mEGFP fluorescence ratios, and synthesized SLF’-MaP555 to evaluate the event rates and apparent off-rates using PAINT imaging (Fig. 4a; Supplementary Fig. 9a-h).

Quantification of the TMR/mEGFP fluorescence ratio revealed that FKBP^F36V, D37A^, FKBP^F36V, Q53A^, and FKBP^F36V, H87A^ exhibited higher brightness than FKBP^F36V^, indicating increased affinity (i.e., lower *K_D_*) (Fig. 4h, Supplementary Fig. 9i). However, PAINT imaging revealed that their higher affinity was largely due to prolonged binding (i.e., lower *k_off_*), leading to the disappearance of localizations primarily due to photobleaching rather than dissociation. As a result, the apparent on- and off-rate calculations for these variants were inaccurate (as indicated by the asterisks in Fig. 4i, 4j), implying that their exchange dynamics are insufficient to support PAINT imaging. For instance, in PAINT imaging, both FKBP^F36V^ and FKBP^F36V, H87A^ initially exhibited high single-molecule localization density benefit from their low *K_D_*. However, the MaP555 fluorophores bound to the protein tag photobleached rapidly, and were not efficiently replenished by fresh SLF’-MaP555. Consequently, localization density dropped sharply and maintained at a low level throughout the 45-minutes (30,000-frame) imaging duration (Fig. 4k,l). Meanwhile, FKBP^F36A^, FKBP^F36V, V55A^, FKBP^F36V, I56A^, FKBP^F36V, E54A^, FKBP^F36V, I90A^, and FKBP^F36V, I91A^ exhibited lower brightness than FKBP^F36V^, suggesting reduced affinity (i.e., higher *K_D_*) (Fig. 4h; Supplementary Fig. 9i). We found that FKBP^F36V, I91A^ exhibited reasonable on- and off-rates for PAINT (slightly outperformed FKBP^F36V, E54A^) (Fig. 4i, 4j). As a result, the binding and dissociation of SLF’-MaP555 with FKBP^F36V, I91A^ could continuously and stably generate localizations, making it well-suited for PAINT imaging (Fig. 4k, 4l). Although optimizing exchange dynamics required some trade-off in affinity and fluorescence intensity during conventional imaging, the brightness of FKBP^F36V, I91A^ remained sufficient to meet the demands of conventional fluorescence microscopy (Supplementary Fig. 9i).

Finally, using the mitochondrial morphology assay, we found 19.0 ± 1.3% of TOM20-mEGFP-FKBP^F36V, I91A^ expressing cells exhibited abnormal mitochondrial morphology (Fig. 4e, 4f), similar to that observed for FKBP^F36V^, and incubating these cells with 100 M SLF’ did not further increase mitochondrial bundling or aggregation (Fig. 4f,g; Supplementary Fig. 10).

Finally, we validated the capability of FKBP^F36V, I91A^ for tagging multiple organelles, as done for FLEXTAG1 and FLEXTAG2. The obtained fluorescence images showed expected subcellular localization and morphology of the targeted cellular structures (Fig. 4m). Based on these results, we designated FKBP^F36V, I91A^ as FLEXTAG3.

To quantitatively assess ligand binding, we purified FLEXTAG proteins from *E. coli* and measured affinities by isothermal titration calorimetry (ITC), yielding *in-vitro K_D_* values of 383 ± 116 nM, 91 ± 19 nM, and 107 ± 14 nM for FLEXTAG1-3, respectively (Supplementary Fig. 11). We further performed a cell-based fluorescence enhancement assay using H2B-mEGFP-FLEXTAG fusions in U2OS cells labeled with a ligand concentration series, which generated apparent cellular *K_D_* values of 159 nM, 68 nM, and 140 nM for FLEXTAG1–3, respectively (Supplementary Fig. 12).

### Development of post-fixation treatments to reduce non-specific binding for FLEXTAG

Although self-renewable protein tags offer many advantages over other fluorescence labeling methods and perform well for live-cell imaging as demonstrated by our three FLEXTAG tag/ligand systems, they face two major technical challenges in fixed-cell imaging: 1) Fixation with PFA or GA induces nonspecific binding of fluorescent ligands to biomolecule surfaces via hydrophobic and electrostatic interactions; and 2) Fixation partially hinders ligand access to the tags, thereby reducing labeling efficiency by preventing a portion of the protein tags from binding to their ligands.

Inspired by immunofluorescence strategies, we systematically optimized fixation and post-fixation treatments. Using TMP–TMR as a model ligand, we compared four fixatives (4% PFA, 4% PFA+0.1% GA, 3% glyoxal^33^, 4% methacrolein^34^) with or without reduction by NH_4_Cl or NaBH_4_. Without reduction, nonspecific binding followed the order glyoxal < PFA < PFA+GA ≈ methacrolein, and both reducing agents lowered background, with NaBH_4_ being most effective (Fig. 5a–c). Morphology preservation was highest with PFA+GA, followed by PFA and glyoxal, and lowest with methacrolein (Supplementary Fig. 13).

**Fig. 5.**
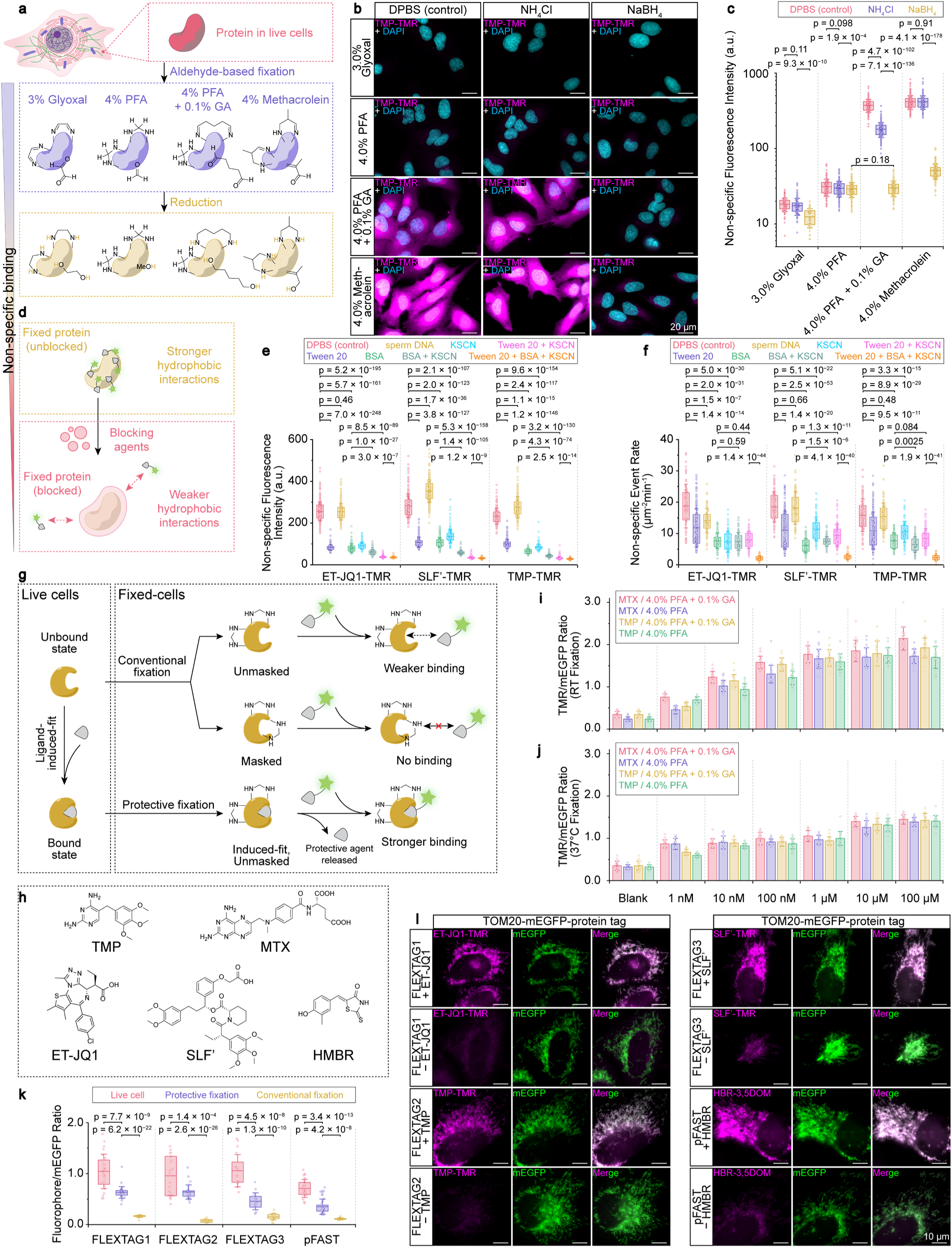
Strategies to reduce non-specific binding and enable protective fixation. **a,** Schematic of chemical reactions occurring on protein surfaces during fixation and the subsequent reduction. Increased surface hydrophobicity of fixed proteins, resulting from crosslink formation and residual aldehydes, are reduced by reducing agents, such as NH_4_Cl or NaBH_4_. **b,** Representative epi-fluorescence images showing non-specific binding in fixed cells. Cells were fixed with indicated fixatives, followed by treatment with DPBS (control), 50 mM NH_4_Cl in DPBS, or 50 mM NaBH_4_ in DPBS. Imaging was performed using 20 nM TMP-TMR in DPBS without washing. Scale bars: 20 μm. **c,** Quantification of non-specific TMP-TMR fluorescence intensity. Each data point represents the average fluorescence intensity of a single cell. *n* = 40–205 cells per condition were examined. **d,** Schematic illustrating the blocking effects of various agents on fixed protein surface, which alter the local microenvironment on the fixed protein surfaces and thereby reduces non-specific binding of ligand-linker-fluorophore conjugates. **e,** Quantification of non-specific fluorescence intensity of ET-JQ1-TMR, SLF’-TMR, or TMP-TMR in cells fixed with 4.0% PFA + 0.1% GA, followed by reduction with 50 mM NaBH_4_ in DPBS. Cells were imaged in 1) DPBS (control); 2) 0.1% Tween 20 (detergent-based blocking agent); 3) 100 μg/mL sperm DNA (nucleotide-based blocking agent); 4) 3.0% BSA (protein-based blocking agent); 5) 300 mM potassium thiocyanate (chaotropic ion); 6) 3.0% BSA + 300 mM KSCN; 7) 0.1% Tween-20 + 300 mM KSCN; 8) 3.0% BSA + 0.1% Tween-20 + 300 mM KSCN. Each data point represents the average fluorescence intensity of a single cell. *n* = 103–388 cells per condition were examined. **f,** Quantification of non-specific binding event rate for 1 nM ET-JQ1-TMR/SLF’-TMR/TMP-TMR in fixed cells, based on SMLM movies (10,000 frames, 10 Hz). *n* = 85–105 randomly selected cellular regions per condition were examined. **g,** Schematic of “protective fixation”. Pre-incubation with protective ligands prior to fixation shields binding pockets from aldehyde-induced modification and locks the protein tag in its ligand-induced-fit conformation. **h,** Chemical structures of protective agents, including ET-JQ1 for FLEXTAG1, TMP and MTX for FLEXTAG2, SLF’ for FLEXTAG3, and HMBR for pFAST. **i,** Quantification of the TMR/mEGFP fluorescence ratio in fixed cells overexpressing TOM20-mEGFP-FLEXTAG2 labeled with 20 nM TMP-TMR. Protective agents TMP/MTX (0–100 μM) were added to the culture medium 1 or 12 hours prior to fixation to allow cellular uptake, followed by fixation with 4.0% PFA or 4.0% PFA + 0.1% GA at room temperature, and imaged with 20 nM TMP-TMR to calculate the TMR/mEGFP ratio. Each data point represents the average TMR/mEGFP ratio of mEGFP-masked regions in a single FOV. *n* = 19–20 FOVs per condition were examined. **j,** Quantification of TMR/mEGFP fluorescence ratio in fixed cells. Cells were fixed at 37 °C. *n* = 12–30 FOVs per condition were examined. **k,** Comparison of the fluorophore (ET-JQ1-TMR/TMP-TMR/SLF’-TMR/HBR-3,5DOM)/mEGFP ratio in live cells, cells fixed by protective fixation, and cells fixed by conventional fixation without protection. Each data point represents the average fluorophore/mEGFP ratio of mEGFP-masked regions in a single FOV. *n* = 16–41 FOVs per condition were examined. **l,** Representative epi-fluorescence images comparing protective fixation and conventional fixation for FLEXTAG1, FLEXTAG2, FLEXTAG3, and pFAST. Scale bars: 10 μm. Boxplots show the mean and boundaries (first and third quartiles); whiskers denote s.d. *p*-values calculated with two-sided unpaired Student’s *t*-test.

We next evaluated blocking agents, including BSA, Tween-20, salmon sperm DNA, KSCN (a chaotropic salt previously reported to disrupt hydrophobic interactions in PAINT imaging^35^). BSA, Tween-20, and KSCN effectively reduced nonspecific binding, and their combination gave the strongest effect in both conventional and PAINT imaging (Fig. 5d–f, Supplementary Fig. 14a). KSCN did not impair FLEXTAG ligand binding, as fluorescence on TOM20 constructs remained unchanged (Supplementary Fig. 14b–c).

Finally, dye properties influenced nonspecific interactions. For six green-channel dyes (Cy2, BODIPY1, BODIPY2, rhodamine 110, fluorescein, and AF488), nonspecific binding correlated with hydrophobicity (LogD) (Supplementary Fig. 15a–c), consistent with prior observations^36^, whereas rhodamine fluorogenic dyes^37,38^ showed reduced nonspecific and free-ligand background, with more fluorogenic dyes producing lower background (Supplementary Fig. 15a, d–f).

Together, these results highlight fixation, reduction, blocking, and fluorophore choice as key factors for robust fixed-cell FLEXTAG imaging.

### Development of protective fixation to preserve post-fixation labeling efficiency of FLEXTAGs

Chemical fixation reduces labeling efficiency of self-renewable protein tags by: 1) aldehyde-based protein crosslinking and Schiff base formation can occur near or within the ligand-binding pocket, potentially limiting or obstructing ligand access; and 2) crosslinking can restrict the protein tag’s ability to undergo ligand-binding-induced conformational adaptation (i.e., ligand-induced fit), which stabilizes ligand binding. To address this, we developed a “protective fixation” strategy in which live cells are preincubated with dye-free (unconjugated) FLEXTAG ligands, followed by fixation in the presence of these ligands. This approach physically excludes aldehydes from the binding pocket and stabilizes the ligand-bound conformation; dye-conjugated ligands are then applied post-fixation for imaging (Fig. 5g).

Using U2OS cells expressing TOM20-mEGFP-FLEXTAG2, fixation with 4.0% PFA + 0.1% GA at room temperature (RT) in increasing concentrations of TMP (0–100 μM) showed TMP-dependent preservation of labeling efficiency that plateaued at 1 μM (Fig. 5h). Substitution with methotrexate (MTX), which binds eDHFR with ∼60-fold higher affinity than TMP^39,40^ (*K_i_* = 21 pM vs. 1.3 nM), did not further improve preservation, likely due to limited cellular uptake (Supplementary Fig. 16). Performing protective fixation at 37°C resulted in poorer labeling efficiency preservation compared to RT, likely due to increased ligand exchange dynamics at higher temperatures that prolonged the duration FLEXTAG2 remained unliganded during fixation (Fig. 5j, Supplementary Fig. 16). Fixation using PFA alone yielded similar trends but resulted in slightly lower labeling preservation compared to PFA + GA (Fig. 5i, 5j; Supplementary Fig. 16).

Protective fixation improved labeling efficiency across all three FLEXTAGs and the GFP-like chromophore-based renewable tag pFAST^11^ when their corresponding protective agents were used (ET-JQ1, TMP, SLF’, or HMBR) (Fig. 5h). Relative to live-cell labeling, post-fixation preservation reached 65.2 ± 7.5% for FLEXTAG1, 66.8 ± 2.7% for FLEXTAG2, 60.5 ± 1.9% for FLEXTAG3, and 49.7 ± 3.3% for pFAST, representing 4.5-, 8.8-, 3.9-, and 3.1-fold improvements over fixation without a protective agent, respectively (Fig. 5k, 5l). These results demonstrate the broad applicability of protective fixation for self-renewable tags.

Finally, cytotoxicity testing showed that most protective agents were well tolerated: after 24 h at 37 °C and 100 μM, only JQ1 caused acute toxicity (within 1–5 h), whereas ET-JQ1, TMP, and SLF’ exhibited only mild effects (Supplementary Fig. 17). Consistent with this, MTT assays showed IC_50_ > 100 μM for ET-JQ1, TMP, and SLF’ (72 h incubation), while HMBR, JQ1, and MTX displayed substantially lower IC_50_ values (34.1 ± 5.5 μM, 33.6 ± 12.8 μM, and 60.3 ± 3.3 μM, respectively) (Supplementary Fig. 18), supporting the suitability of FLEXTAG ligands for protective fixation.

### FLEXTAG enables PLIM-based nanoscopic imaging with anti-fade performance

After establishing the FLEXTAG framework including three small self-renewable protein tags, methods for reducing non-specific binding, and a labeling efficiency preservation strategy (i.e, protective fixation), we evaluated its performance in enabling anti-fade multi-color fluorescence nanoscopic imaging. Fluorescence nanoscopy methods fall into two main categories: 1) PLIM uses structured or patterned illumination to improve spatial resolution; and 2) SMLM relies on the precise localization of single fluorophores that stochastically blink on and off, enabling the reconstruction of super-resolved images with nanometer precision.

We first assessed FLEXTAG in SIM, a fast PLIM-based nanoscopy offering ∼100 nm spatial resolution. Unlike the pFAST ligand, which is a fluorescent chromophore, FLEXTAGs’ ligand-binding components are non-fluorescent, allowing greater flexibility in spectral selection for multi-color imaging. To demonstrate this versatility, we synthesized FLEXTAG ligands conjugated to five dyes spanning distinct spectral ranges: four cell-permeable dyes (JF525, TMR, MaP555, and JF635) suitable for both live and fixed-cell imaging^37,38,41^ and one cell-impermeable dye (AF594) for fixed-cell imaging only (Fig. 6a). These enabled three-color live-cell co-staining of cells co-expressing TOM20-FLEXTAG1, MAP7-FLEXTAG2, and FLEXTAG3-Sec61β, targeting mitochondria, microtubules, and the ER, respectively. FLEXTAG-SIM imaging enabled us to directly visualize the spatial interactions among these organelles (Fig. 6b; Supplementary Movie 1). We further evaluated FLEXTAGs’ performance in fixed cells using our protective fixation and nonspecific binding reduction protocols. SIM images from fixed cells closely resembled those from live cells, validating the applicability of FLEXTAG to fixed-cell imaging (Fig. 6d).

**Fig. 6.**
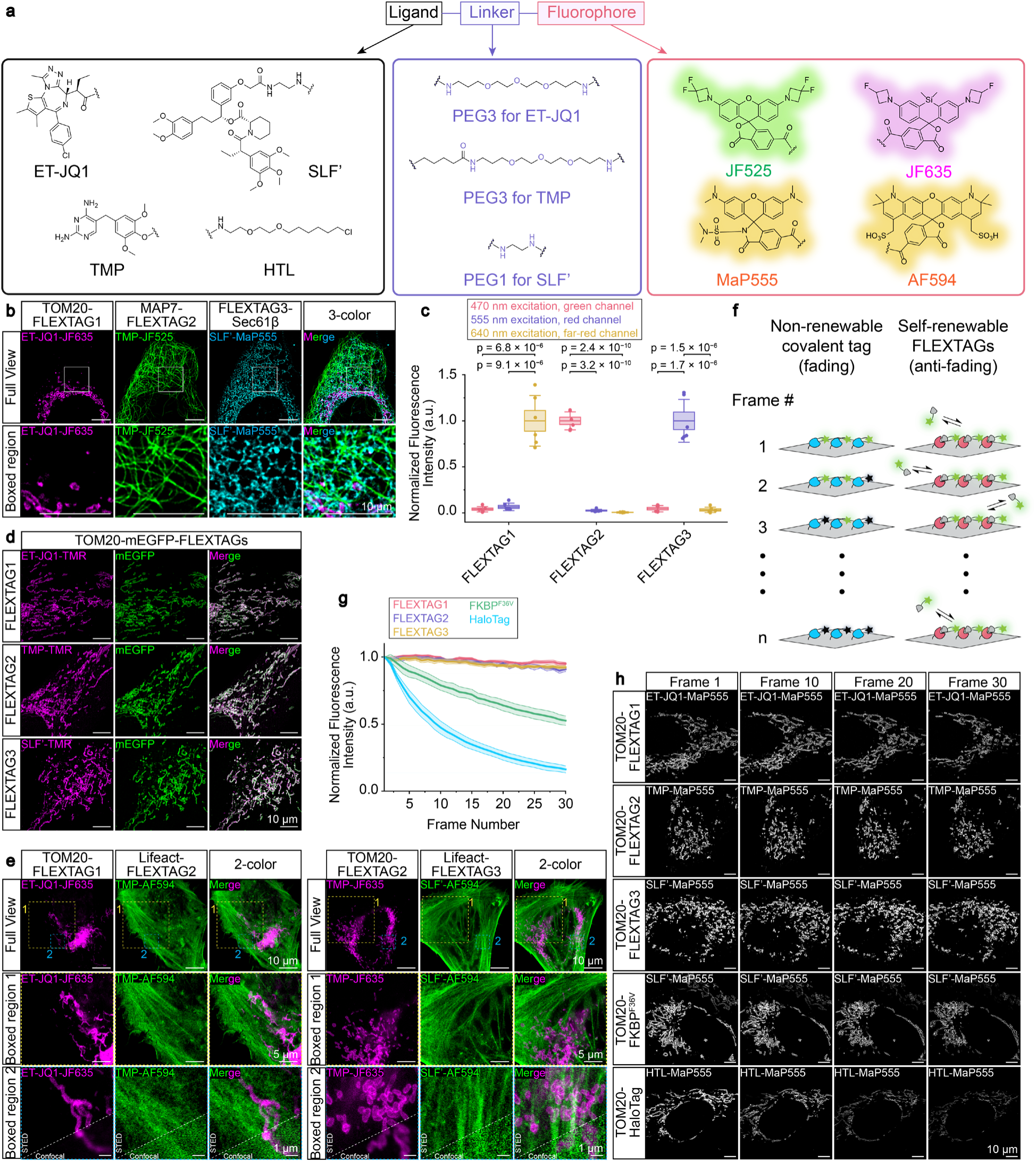
FLEXTAG enables PLIM-based nanoscopic imaging with anti-fade performance. **a,** Chemical structures of ligand-linker-fluorophore conjugates targeting FLEXTAG1, FLEXTAG2, FLEXTAG3, and HaloTag for PLIM-based super-resolution imaging. **b,** Top: Representative three-color live-cell SIM images of cells stably expressing TOM20-FLEXTAG1, MAP7-FLEXTAG2, and Sec61β-FLEXTAG3. Bottom: Magnified view of the boxed region highlighting the close spatial interactions among mitochondria, microtubules, and the ER. Samples were labeled with 100 nM ET-JQ1-JF635, 100 nM TMP-JF525, and 100 nM SLF’-MaP555. Scale bars: 10 μm. **c,** Verification of orthogonality of FLEXTAGs. For each FLEXTAG, six regions were manually selected from corresponding organelles, and average fluorescence intensities of the green (470 nm excitation), red (555 nm excitation), and far-red (640 nm excitation) channels in the selected regions were quantified. Boxplots show the mean and boundaries (first and third quartiles); whiskers denote s.d. *p*-values calculated with two-sided unpaired Student’s *t*-test. **d,** Representative fixed-cell SIM images of cells overexpressing TOM20-mEGFP-FLEXTAG1/FLEXTAG2/FLEXTAG3, labeled with 100 nM ET-JQ1-TMR, TMP-TMR, or SLF’- TMR. Scale bars: 10 μm. **e,** Representative two-color fixed-cell STED images with magnified views showing the potential spatial interactions between actin stress fibers and mitochondria. Left: Cells overexpressing TOM20-mEGFP-FLEXTAG1 and Lifeact-mEGFP-FLEXTAG2, labeled with 200 nM ET-JQ1-JF635 and 100 nM TMP-AF594; Right: Cells overexpressing TOM20-mEGFP-FLEXTAG2 and Lifeact-mEGFP-FLEXTAG3, labeled with 200 nM TMP-JF635 and 100 nM SLF’-AF594. Scale bars: 10 μm for full view, 5 μm for boxed region 1 (yellow), and 1 μm for boxed region 2 (blue). **f,** Schematic illustrating the anti-fading mechanism of self-renewable FLEXTAGs during prolonged imaging. Fluorescence intensity of non-renewable covalent protein tags progressively decreases over time due to the irreversible photobleaching of fluorophores (fading). In contrast, self-renewable protein tags continuously replace photobleached fluorophores with intact ones, thereby maintaining stable fluorescence intensity (anti-fading). **g,** Quantification of fluorescence intensity of FLEXTAG1/FLEXTAG2/FLEXTAG3/FKBP^F36V^, and HaloTag over 30-frame live-cell SIM imaging, labeled with 100 nM ET-JQ1-MaP555/TMP-MaP555/SLF’-MaP555/SLF’-MaP555/HTL-MaP555. *n* = 3 cells were imaged independently for each protein tag. Each frame was acquired at 1-minute intervals, and fluorescence intensities were normalized to the first frame. Data are presented as mean ± s.e.m. **h,** Representative time-lapse SIM images of a single live cell overexpressing TOM20-mEGFP-FLEXTAG1, FLEXTAG2, FLEXTAG3, FKBP^F36V^ or HaloTag, labeled with 100 nM MaP555 conjugated ligand. Images are shown at frames 1, 10, 20, and 30. Scale bars: 10 μm.

To assess the orthogonality of our three FLEXTAGs, we quantified the three-color SIM images and observed minimal crosstalk between color channels (Fig. 6c), demonstrating the high orthogonality of our three protein tags for multiplexed imaging.

Next, we assessed FLEXTAG in STED, a PLIM-based nanoscopy offering higher spatial resolution than SIM by incorporating a depletion laser beam. FLEXTAG enabled two-color STED imaging in fixed cells co-expressing TOM20-FLEXTAG1 and Lifeact-FLEXTAG2 (or TOM20-FLEXTAG2 and Lifeact-FLEXTAG3), showing the expected mitochondrial and actin cytoskeleton localizations, respectively, along with potential interactions between mitochondria and actin stress fibers (Fig. 6e). Fluorescence intensity profiles across the mitochondrial outer membrane revealed a marked resolution improvement over conventional confocal microscopy (Supplementary Fig. 19). Notably, our two-color FLEXTAG-STED imaging was performed on a STEDYCON system equipped with a single depletion laser, which limits emission depletion to two color channels. We anticipate that three-color STED imaging would be feasible with a STED microscope featuring multiple depletion lasers covering a broader wavelength range.

Photobleaching remains a major limitation for PLIM-based nanoscopy: SIM requires acquisition of tens of images to reconstruct a single SIM image; STED’s high-power depletion lasers exacerbate fluorophore bleaching. FLEXTAG addresses this by dynamically replacing photobleached ligands with fresh fluorescent ones, maintaining relatively stable signals over time (Fig. 6f). To quantify this anti-fade performance, we acquired time-lapse SIM images of TOM20 tagged with FLEXTAG1, FLEXTAG2, FLEXTAG3, HaloTag (a non-renewable protein tag), or FKBP^F36V^ (the protein tag described above with slow renewable rate), each labeled with MaP555-conjugated ligands. Over 30 consecutive frames, fluorescence intensity dropped to ∼20% for HaloTag and ∼50% for FKBP^F36V^. In contrast, the fluorescence intensity of all three FLEXTAGs remained nearly constant (Fig. 6g,h). Repeating the experiment with JF635-conjugated ligands yielded similar results (Supplementary Fig. 20). These findings highlight FLEXTAG’s rapid self-renewal kinetics and its ability to overcome photobleaching during PLIM-based nanoscopic imaging.

### FLEXTAG enables SMLM-based nanoscopic imaging with anti-fade performance

In SMLM, a super-resolved image is reconstructed from a time series of frames, each capturing a sparse subset of fluorescently labeled protein targets that can be localized at the single-molecule level. While SMLM typically achieves higher spatial resolution than PLIM, it requires slower image acquisition. STORM/PALM achieve this sparsity by using photoswitchable fluorophores that stochastically blink on and off; A common PAINT strategy uses dye-conjugated DNA oligonucleotides that transiently hybridize with complementary strands linked to antibody-labeled targets.

We first assessed FLEXTAG in PAINT. FLEXTAG-PAINT offers intrinsic advantages over DNA-PAINT: it does not rely on antibody availability, and its smaller labeling size provides a more accurate representation of protein localization (Fig. 7e). Using FLEXTAG ligands conjugated to green (AF488), red (TMR/MaP555), or far red (JF635/HMSiR/AF647) dyes, we obtained three-dimensional (3D) PAINT images for all three FLEXTAG tag/ligand systems, demonstrating that each tag/ligand pair supports self-renewable dynamics compatible with PAINT (Fig. 7a-d). FLEXTAG tag/ligand pairs can also be combined or integrated with established PAINT probes, such as phalloidin-PAINT^35^ for actin cytoskeleton imaging, enabling multi-color PAINT imaging. Three-color PAINT imaging enabled direct visualization of the spatial interactions among actin stress fibers, mitochondria, and the ER (Fig. 7f).

**Fig. 7.**
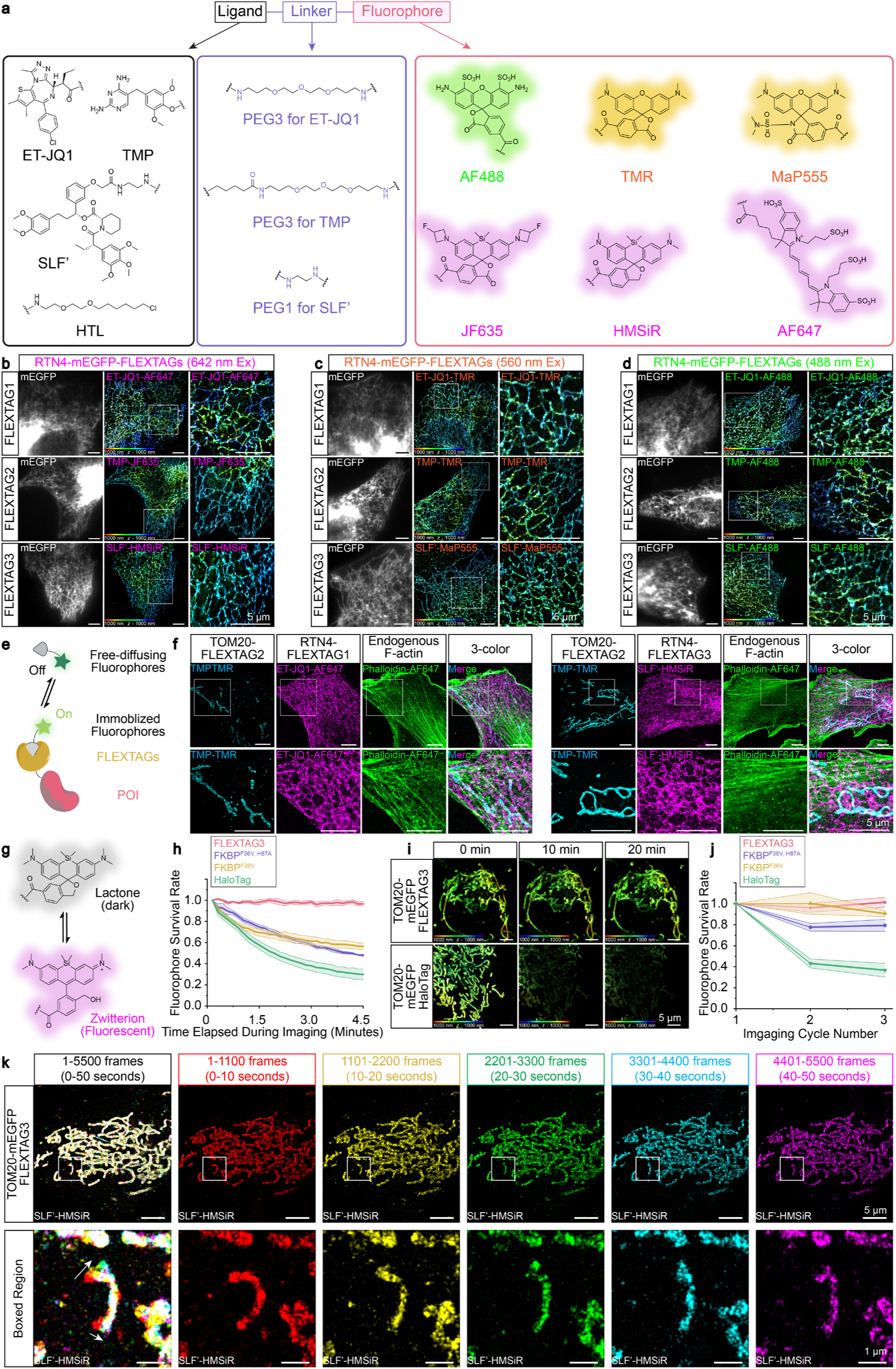
FLEXTAG enables SMLM-based nanoscopic imaging with anti-fade performance. **a,** Chemical structures of ligand-linker-fluorophore conjugates targeting FLEXTAG1, FLEXTAG2, or FLEXTAG3 for SMLM-based super-resolution imaging. **b,** Far-red channel (642 nm excitation) single-color 3D-PAINT images of cells overexpressing RTN4-mEGFP-FLEXTAG1/FLEXTAG2/FLEXTAG3. FLEXTAGs were imaged with 1 nM ET-JQ1-AF647, 1 nM TMP-JF635, and 10 nM SLF’-HMSiR, respectively. Scale bars: 5 μm. **c,** Red channel (560 nm excitation) single-color 3D-PAINT images of cells overexpressing RTN4-mEGFP-FLEXTAG1/FLEXTAG2/FLEXTAG3. FLEXTAGs were imaged with 1 nM ET-JQ1-TMR, 1 nM TMP-TMR, and 1 nM SLF’-MaP555, respectively. Scale bars: 5 μm. **d,** Green channel (488 nm excitation) single-color 3D-PAINT images of cells overexpressing RTN4-mEGFP-FLEXTAG1/FLEXTAG2/FLEXTAG3. FLEXTAGs were labeled with 1 nM ET-JQ1-AF488, 1 nM TMP-AF488, and 1 nM SLF’-AF488, respectively. Scale bars: 5 μm. **e,** Schematic of FLEXTAG-PAINT. Fluorophore-conjugated FLEXTAG ligands transiently bind to self-renewable FLEXTAGs, producing stochastic localization events. Repeated binding and dissociation allow for the accumulation of single-molecule localizations over time, enabling reconstruction of a super-resolution (PAINT) image. **f,** Representative three-color PAINT images with magnified views showing the spatial interactions between actin stress fibers, mitochondria, and the ER. Left: Cells overexpressing TOM20-FLEXTAG2 and RTN4-FLEXTAG1, imaged with 1 nM TMP-TMR, 1 nM ET-JQ1-AF647, and 1 nM phalloidin-AF647 (For endogenous F-actin imaging). Right: Cells overexpressing TOM20-FLEXTAG2 and RTN4-FLEXTAG3, labeled with 1 nM TMP-TMR, 30 nM SLF’-HMSiR, and 1 nM phalloidin-AF647. Scale bars: 5 μm. **g,** Schematic of the HMSiR self-blinking mechanism for SMLM. HMSiR spontaneously switches between a dark lactone form and a fluorescent zwitterion form. **h,** Quantification of fluorophore survival rate in a single live-cell STORM movie (30,000 frames, 110 Hz, with total data acquisition time of 4.5 min). *n* = 3 biological replicates, 3 FOVs were examined for each protein tag. Data are presented as mean ± s.e.m. **i,** Representative live-cell STORM images of cells overexpressing TOM20-mEGFP-FLEXTAG3 or TOM20-mEGFP-FKBP variants labeled with 30 nM SLF’-HMSiR, and cells overexpressing TOM20-mEGFP-HaloTag labeled with 30 nM HTL-JF630b (HMSiR has a ∼9-fold higher duty cycle than JF630b so HMSiR is more prone to photobleaching^46^)from three imaging cycles. Each cycle contains 4.5 min data acquisition time (30,000 frames, 110 Hz) and 5.5 min dark time. Scale bars: 5 μm. **j,** Quantification of fluorophore survival rates over three STORM imaging cycles. *n* = 3 biological replicates; 3 FOVs were examined for each protein tag. Data are presented as mean ± s.e.m. **k,** Live-cell STORM images of TOM20-mEGFP-FLEXTAG3 in the same FOV over 0–50 seconds (5,500 frames at 110 Hz). The boxed region with a magnified view highlights the gradual transition of mitochondria from a curved to a straight morphology over successive 10-second intervals (1,100 frames each). Scale bars: 5 μm (full view), 1 μm (boxed region).

We next assessed FLEXTAG in STORM. Unlike PAINT, STORM relies on covalent labeling with photoswitchable dyes or fluorescent proteins, which are prone to photobleaching, limiting long-term imaging of dynamics processes^42^. FLEXTAG-STORM, which uses dynamically exchanging photoswitchable ligands, can overcome this limitation. We used FLEXTAG3 and SLF’ as a model system and synthesized a chimeric ligand by conjugating SLF’ to HMSiR (a cell-permeable, spontaneously photoswitchable Si-rhodamine dye developed for live-cell STORM^43,44^) (Fig. 7g). We first confirmed that SLF’-HMSiR successfully labeled fixed cells expressing TOM20-FLEXTAG3 (Supplementary Fig. 21a). Live-cell FLEXTAG-STORM imaging, acquired at a temporal resolution of 10 seconds, captured dynamic mitochondrial morphology (Fig. 7k) and enabled 3D reconstruction (Supplementary Fig. 21b; Supplementary Movie 2). To evaluate anti-fade performance, we compared FLEXTAG3 with slower-renewing precursor tags, FKBP^F36V^ and FKBP^F36V, H87A^, and the non- renewable HaloTag. We monitored the number of single-molecule localizations per image frame at 110 Hz, generating apparent photobleaching curves. FLEXTAG3 retained 96.6 ± 2.3% of single-molecule localizations per frame after ∼ 4.5 minutes (30,000 frames), compared to 56.5 ± 3.1%, 48.0 ± 1.1%, and 29.9 ± 5.2% for FKBP^F36V^, FKBP^F36V, H87A^, and HaloTag, respectively (Fig. 7h). Furthermore, FLEXTAG3 showed minimal photobleaching in multi-cycle time-lapse live-cell STORM imaging, whereas the other tags exhibited progressive signal loss over time (Fig. 7i,j; Supplementary Fig. 21c). These results demonstrate FLEXTAG-STORM’s superior anti-fade performance.

## Discussion

We have developed FLEXTAG, a self-renewable protein tagging and labeling framework that enables robust multi-color fluorescence nanoscopy. Through extensive evaluation across multiple nanoscopy modalities with distinct working principles, including SIM, STED, PAINT, and STORM, we demonstrated that FLEXTAG supports 3D multi-color super-resolution imaging in both fixed and live-cell conditions. Because nanoscopy imposes stringent performance requirements, FLEXTAG should be readily applicable to conventional fluorescence microscopy as well.

While self-renewable tags like dHaloTag^14^ (a self-renewable mutant form of HaloTag) and GFP-like chromophore-based renewable protein tags (e.g., pFAST^11^) have been developed, FLEXTAG1, FLEXTAG2, and FLEXTAG3 are distinct in combining self-renewability with a small tag size (∼half that of dHaloTag), minimal aggregation, superior apparent photostability (chromophore-based renewable protein tags suffer from poor photostability and brightness; see Supplementary Fig. 1a-c), and easier expansion to the use of bright organic dyes for multiplexed imaging (chromophore-based renewable protein tags are not easily expandable to other colors because their ligands’ fluorescent component is inseparable from the binding motif). Each FLEXTAG tag/ligand system incorporates a dynamic ligand exchange mechanism that enables continuous replenishment of dye-conjugated ligands, maintaining stable fluorescence intensity over extended imaging durations and outperforming HaloTag and the slower-renewing FKBP^F36V^ tag in photobleaching tests. This feature should benefit all nanoscopy techniques examined, where low-brightness fluorophores and photobleaching often compromise image resolution and duration, as well as long-term conventional fluorescence imaging, where extended imaging enables prolonged tracking of protein or organelle dynamics.,

The small size of FLEXTAGs, all under 20 kDa, is crucial for preserving correct protein localization and function. In contrast, the well-known HaloTag, is relatively large (∼33 kDa) and mislocalizes proteins, such as COX8, ATP5ME, YTHDC1, and the coronaviral envelope protein. Our three FLEXTAGs preserved proper localization for these proteins, supporting broader applicability for minimally perturbing cellular targets (Supplementary Fig. 1d). Moreover, FLEXTAG’s small size improves spatial accuracy in imaging by minimizing the dye–target displacement, especially compared to bulky antibody-based methods (∼150 kDa for IgGs, ∼15 nm span^45^).

FLEXTAGs also exhibited minimal aggregation, regardless of ligand presence. Across proteins localized to diverse subcellular compartments, FLEXTAG consistently preserved expected morphology and distribution. This trait is advantageous for both imaging and non-imaging applications. The precursor proteins of FLEXTAG have previously enabled ligand-inducible manipulations, such as ligand-induced proteolysis of the tagged protein or ligand-induced interactions between tagged proteins. In such contexts, tag-induced aggregation can alter protein localization and function even before the ligand is added to trigger proteolysis or interaction, potentially leading to artifacts or incorrect conclusions. FLEXTAG’s minimal aggregation property should improve data reliability in these applications.

As part of our FLEXTAG framework, we developed protective fixation and methods for reducing nonspecific binding of fluorescent ligands, enabling high-quality fixed-cell imaging for self-renewable protein tags. Previous self-renewable protein tags, such as dHaloTag and pFAST, were primarily used for live-cell imaging, likely because fixation often led to greatly reduced labeling efficiencies, resulting in much lower fluorescence intensity. FLEXTAG preserves labeling efficiency and enables higher signal-to-noise ratio imaging. In the future, exploring additional types of blocking agents may help further suppress nonspecific binding. Furthermore, FLEXTAG’s compatibility with both live- and fixed-cell imaging makes it promising for applications in correlated light and electron microscopy (CLEM)^42^. In CLEM, live-cell fluorescence imaging visualizes protein dynamics, while electron microscopy (EM), following fixation, reveals ultrastructure of organelles or protein complexes. This combined approach may provide new insights into the relationship between protein complex dynamics and their ultrastructural organization with EM resolutions.

SMLM methods such as STORM and PAINT offer higher spatial resolution than PLIM techniques like SIM and STED, but live-cell SMLM has long been constrained by photobleaching. While dHaloTag has enabled live-cell STED and shows promise for live-cell SMLM^14^, this application has not been demonstrated. Using FLEXTAG-STORM, we achieved prolonged live-cell SMLM imaging with minimal photobleaching while leveraging bright photoswitchable dyes for enhanced spatial resolution (which scales with dye photon output).

Finally, FLEXTAG’s “one-for-all” design enables researchers to label proteins once and image them using any of the major nanoscopic techniques, including SIM, STED, PAINT, and STORM, based on experimental requirements. Each technique has distinct strengths and limitations: SMLM offers the highest resolution but requires longer acquisition times; STED provides fast imaging but demands specialized optics and specific fluorophores; and SIM offers the broadest fluorophore compatibility, albeit with lower resolution. Three FLEXTAGs support all three imaging regimes with high orthogonality with minimal crosstalk when co-expressed and co-stained in cells and can flexibly pair with fluorophores across the green, red, and far-red channels, making FLEXTAG a convenient and versatile toolbox and framework for anti-fade multi-color nanoscopic imaging (Supplementary Tables 1–3). Among the three FLEXTAGs, FLEXTAG2 exhibited the most favorable kinetics, with higher binding affinity and faster apparent on/off rates than FLEXTAG1 and FLEXTAG3, making it the preferred choice for PAINT. FLEXTAG3 displayed mild aggregation compared to FLEXTAG1 and FLEXTAG2. Otherwise, all three FLEXTAGs showed comparable imaging resolution.

In summary, FLEXTAG provides a compact, minimally aggregating, and self-renewable tagging framework compatible with diverse nanoscopy techniques and imaging conditions. Its unique combination of small tag size, superior photostability, bright organic dye compatibility, and orthogonal multiplexing capabilities enables prolonged, high-resolution imaging in both live and fixed cells. By overcoming key limitations of previous renewable tags, FLEXTAG opens new opportunities for visualizing dynamic protein behaviors and deciphering molecular mechanisms with nanoscopic precision, offering broad utility for advancing cell biology and biomedical research.

## Methods

### Plasmids construction

All plasmids encoding TOMM20, ATP5ME, RTN4, H2B, Zyxin, MAP7, vimentin, and lifeact (proteins to be tagged on the C terminus) were modified based on the pAG156 Mito-FAST plasmid (Addgene #130723) by replacing the COX8 sequence with other organelle-targeting sequences, and the sequence of FAST was replaced by other protein tags as needed. All plasmids encoding YTHDC1, Sec61β, and coronaviral envelope (E) protein (proteins to be tagged on the N terminus) were modified based on pAG876 pDisplay-pFAST (Addgene #172868) by replacing the PDGFRβ-transmembrane domain by other organelle-targeting sequences, and the sequence of pFAST was replaced by other protein tags as needed. All plasmids encoding YTHDC1, Sec61β, and the coronaviral envelope (E) protein (proteins to be tagged on the N terminus) were modified based on pAG876 pDisplay-pFAST (Addgene #172868) by replacing the PDGFRβ-transmembrane domain by other organelle-targeting sequences, and the sequence of pFAST was replaced by other protein tags as needed.

FLEXTAG1 was cloned from GFP-BRD4 plasmid (Addgene, #65378), with primers 5’-AAGGACGTGCCCGACTCTC-3’ and 5’-CTCGTCCGGCATCTTGGCC-3’.

Single point mutations were introduced by quick-change.

W42A mutation was introduced using primers 5’-CTACGCCGCACCCTTCTACAAGCCTGTGGAC-3’ and 5’-AAGGGTGCGGCGTAGGCGGCGTG-3’.

L55A mutation was introduced using primers 5’-CTGGGCGCACACGACTACTGTGAC-3’ and 5’-GTCGTGTGCGCCCAGTGCCTCCAC-3’.

Q21A mutation was introduced using primers 5’-CGGAGGCACTCAAGTGCTGCAGCGG-3’ and 5’-ACTTGAGTGCCTCCGAGACCTTGCTGCTCTTC-3’.

Y98K mutation was introduced using primers 5’-CAACTGCAAGAAGTACAACCCTCCTGACCA TGAG-3’ and 5’-GTACTTCTTGCAGTTGGAGAACATCAATCGGACGTCAGCAC-3’.

H105A mutation was introduced using primers 5’-CTCCTGACGCCGAGGTGGTGGCCATG-3’ and 5’-CCTCGGCGTCAGGAGGGTTGTACTTATAGCAGTTG-3’.

Q115A mutation was introduced using primers 5’-CAAGCTCGCCGATGTGTTCGAAATGCGCTTTG-3’ and 5’-ACATCGGCGAGCTTGCGGGCCATG-3’.

M120G mutation was introduced using primers 5’-GTGTTCGAAGGTCGCTTTGCCAAGATGCC-3’ and 5’-AAGCGACCTTCGAACACATCCTGGAGCTTG-3’.

F122E mutation was introduced using primers 5’-AATGCGCGAAGCCAAGATGCCGGACG-3’ and 5’-CTTGGCTTCGCGCATTTCGAACACATCCTGGAG-3’.

FLEXTAG2 was cloned from mCherry-eDHFR (Addgene #107266), with primers 5’-ATCAGTCTGATTGCGGCGTTAGC-3’ and 5’-CCGCCGCTCCAGAATCTCAAAG-3’. P89C mutation was introduced by quick-change using primers 5’-GGTGACGTATGTGAAATCATGGTGATT GGCGGCG-3’ and 5-

GATTTCACATACGTCACCACACGCCGCGATGG-3’. P39C mutation was introduced by primers 5’- CTTAAATAAATGTGTGATTATGGGCCGCCATACCTGG-3’ and 5’-CACACATTTATTTAAGGTGTTGCGTTTAAACCAGGCGAGATC-3’. eDHFR^P39GCG^, eDHFR^W22C, H149C^, eDHFR^W22GCG, W149GCG^, were obtained from custom gene fragment synthesis (Twist Biosceince).

For eDHFR linker screening experiments, linker sequences were encoded in primers and directly inserted between mEGFP and eDHFR^WT^/FELXTAG2. E.g. VDVVDDIKLT linker for FLEXTAG2 were introduced using primers 5’-GTAGATGTCGTAGATGATATCAAGCTTACCTGTGGTGGTG GTGGTGGTATG-3’ and 5’-ATCATCTACGACATCTACCTTGTACAGCTCGTCCATGCC-3’.

FLEXTAG3 was cloned from PM-FRB-mRFP-T2A-FKBP-5-ptase (Addgene #40896), with primers 5’-GGAGTGCAGGTGGAAACCATCTC-3’ and 5’-TTCCAGTTTTAGAAGCTCCACATCGAAG-3’.

F36V mutation was introduced using primers 5’-GGAAAGAAAGTGGATTCCTCCCGGGACAGAAAC-3’ and 5’-ATCCACTTTCTTTCCATCTTCAAGCATCCCGGTGTAG-3’.

F36L mutation was introduced using primers 5’-GGAAAGAAACTCGATTCCTCCCGGGACAGAAAC-3’ and 5’-ATCGAGTTTCTTTCCATCTTCAAGCATCCCGGTGTAG-3’.

F36A mutation was introduced using primers 5’-GAAAGAAAGCCGATTCCTCCCGGGACAGAAA CAAGC-3’ and 5’-GAATCGGCTTTCTTTCCATCTTCAAGCATCCCGGTG-3’.

D37A mutation was introduced using primers 5’-GAAAGTCGCTTCCTCCCGGGACAGAAACAAGC-3’ and 5’-GAGGAAGCGACTTTCTTTCCATCTTCAAGCATCCCG-3’.

Q53A mutation was introduced using primers 5’-CAAGGCGGAGGTGATCCGAGGC-3’ and 5’-GATCACCTCCGCCTTGCCTAGCATAAACTTAAAGG-3’.

E54A mutation was introduced using primers 5’-CAGGCGGTGATCCGAGGCTGGGAAG-3’ and 5’-CCTCGGATCACCGCCTGCTTGCCTAGCATAAAC-3’.

V55A mutation was introduced using primers 5’-GGAGGCGATCCGAGGCTGGGAAGAAGGG-3’ and 5’-CCTCGGATCGCCTCCTGCTTGCCTAGCATAAAC-3’.

I56A mutation was introduced using primers 5’-GGAGGTGGCCCGAGGCTGGGAAGAAGGG-3’ and 5’-CCTCGGGCCACCTCCTGCTTGCCTAGCATAAAC-3’.

H87A mutation was introduced using primers 5’-CCACTGGGGCCCCAGGCATCATCCCACC-3’ and 5’-CCTGGGGCCCCAGTGGCACCATAGGCATAATC-3’.

I90A mutation was introduced using primers 5’-CAGGCGCCATCCCACCACATGCCACTCTCG-3’ and 5’-GGTGGAATGGCGCCTGGGTGCCCAGTG-3’.

I91A 5’-GGCATCGCCCCACCACATGCCACTGTCTTC-3’ and 5’-GTGGTGGGGCGATGCCTGGG TGCCCAG-3’.

HaloTag7 was cloned from pCDNA5/FRT/TO_TOMM20_HaloTag7 plasmid (Addgene #169330), with primers 5’-GAAATCGGTACTGGCTTTCCATTCGAC-3’ and 5’-GCCGGAAATCTCGAGCGTG-3’.

TOMM20 was cloned from mTagBFP2-TOMM20-N-10 plasmid (Addgene #55328), with primers 5’-ATGGTGGGTCGGAACAGCG-3’ and 5’-TTCCACATCATCTTCAGCCAAGCTC-3’.

RTN4 was cloned from RTN4b-GFP plasmid (Addgene #186952), with primers 5’-ATGGAAGACCTGGACCAGTCTC-3’ and 5’-TTCAGCTTTGCGCTTCAATCCAG-3’.

H2B was cloned from pAG657 H2B-pFAST plasmid (Addgene #172862), with primers 5’-ATGCCCGAACCTGCGAAG-3’ and 5’-CTTGGAGCTGGTGTACTTGGTCAC-3’.

Zyxin was cloned from mEos4a-Zyxin-6 plasmid (Addgene #57499), with primers 5’-CACCGCCACCATGGCGGCCCCCCGCCCG-3’ and 5’-GACCGGTGGATCCGTCTGGGCTCTAGCA G-3’.

MAP7 was cloned from Ensconsin-18-mEos2 plasmid (Addgene #99230), with primers 5’-CGGACTCAGATCCGAATTCGCCACCGCCACCATGGCGGAGCTAGGAGCTG-3’ and 5’-TATAACTTCTGCAGTCTGCTGTGTCTG-3’.

Vimentin was cloned from Vimentin-APEX2 in pECFP plasmid (Addgene #66170), with primers 5’-ATGTCCACCAGGTCCGTGTC-3’ and 5’-TTCAAGGTCATCGTGATGCTGAGAAGTTTC-3’.

Lifeact sequence was encoded in primers 5’-GATCAAGAAGTTCGAGAGCATCAGCAAGGAGGA GATCCATTCGTTGAGATCTGCCACC-3’ and 5’-CTCGAACTTCTTGATCAGGTCGGCCACGCC CATGGTGGCGGTGGCGAATTC-3’ and directly inserted into pIRES vectors.

YTHDC1 was cloned from pcDNA3-FLAG-HA-hYTHDC1 plasmid (Addgene #85167), with primers 5’-ATGGCGGCTGACAGTCG-3’ and 5’-TCTTCTATATCGACCTCTCTCCCCTC-3’.

ATP5ME, Sec61β, and the coronaviral envelope (E) protein^15^ were obtained from gene fragment synthesis (Twist Bioscience).

GFP-MAPPER-FLEXTAG was modified from GFP-MAPPER plasmid (Addgene #117721), with FLEXTAGs inserted between (GA)_6_L and (EAAAR)_4_.

### U2OS cell culture

Human bone osteosarcoma epithelial cells (U2OS cells; ATCC, HTB-96) were plated onto 18 mm coverslips (Electron Microscopy Sciences, 72222-01) in a 12-well plate and maintained in complete growth Dulbecco’s modified Eagle’s medium (DMEM; Corning, 10-013-CV), supplemented with 10% (vol/vol) fetal bovine serum (FBS, Gibco, A52567-01), 0.1% (vol/vol) penicillin–streptomycin (Gibco, 10378-016). For cell passage, 0.25% Trypsin-EDTA (Corning, 25-0530CI) was used to detach adherent cells from the culture surface.

### Cell transfection

Transfection was performed using TransIT-LT1 (Mirus, MIR 2300) according to the manufacturer’s protocol. Briefly, when cells reached 60-80% confluency, 1.0 μg of plasmid encoding the gene of interest was diluted in 100 µL of Opti-MEM (Gibco, 31985-070). The diluted DNA was then mixed with 3 µL of TransIT-LT1 reagent, gently pipetted to mix, and incubated for 15 minutes at room temperature (RT) to allow complex formation. The transfection mixture was subsequently added dropwise to the culture medium. After 12 hours, the medium was replaced with fresh DMEM supplemented with 10% fetal bovine serum (FBS) to promote cell recovery and expression of the transgene. Live-cell imaging or cell fixation experiments were performed 48 hours post-transfection.

### Stable cell line establishment

Stable cell lines were generated using the PiggyBac transposon system. The gene of interest was cloned into a transposon plasmid based on the Xlone vector backbone (Addgene, Plasmid 96930). At 60-80% confluency in 6-well plates, cells were cotransfected with 1.6 µg of the transposon plasmid and 0.4 µg of the transposase helper plasmid (PBase) plasmid using the TransIT-LT1 reagent. 48 hours after transfection, cells were subjected to selection with Blasticidin (BSD) at a final concentration of 10 µg/mL. Selection was maintained for 7–14 days, with medium replacement every 2–3 days.

### Protective fixation and non-specific binding reduction treatments

Cultured U2OS cells were incubated with 100 μM protective ligand (ET-JQ1, TMP, SLF’, or HMBR) at 37 °C in a 5.0% CO₂ incubator for 1 hour (12 hours for MTX). Cells were subsequently fixed with a fixation buffer containing 4.0% (wt/vol) paraformaldehyde (PFA, Electron Microscopy Sciences, 15714), 0.1% glutaraldehyde (GA, Electron Microscopy Sciences, 10619), 4.0% (wt/vol) sucrose, and 100 μM protective ligand in Dulbecco’s phosphate buffered saline (DPBS, Leinco Technologies, D388). Fixation was carried out for 15 min at room temperature (RT), followed by three washes with DPBS. Cells were then permeabilized with 0.2% (vol/vol) Triton X-100 in DPBS for 30 min at RT. After permeabilization, cells were washed three times with DPBS and treated with 50 mM sodium borohydride (NaBH_4_) or ammonium chloride (NH_4_Cl), for 1 hour to quench excess aldehydes. Finally, cells were then washed three times with DPBS and blocked overnight at 4 ℃ in blocking buffer containing 3.0% (wt/vol) bovine serum albumin (BSA), 0.1% Tween-20, and 300 mM Potassium thiocyanate (KSCN) in DPBS. To optimize the protective fixation protocol described above, variations in incubation temperature, protective ligand concentration, and fixation buffer composition (using 4.0% (w/v) PFA with 4.0% sucrose in DPBS supplemented with ligand) were tested during cell fixation.

Without reducing agent treatment (NH_4_Cl or NaBH_4_), the levels of non-specific binding followed the order: Glyoxal < PFA < PFA + GA ≈ methacrolein. Both reducing agents significantly reduced non-specific binding, with NaBH_4_ yielding a much stronger reduction than NH_4_Cl. After NaBH_4_ treatment, the order of non-specific binding levels changed to: Glyoxal < PFA ≈ PFA + GA < methacrolein (Fig. 5b, 5c). To assess structural preservation, we used IF to examine nuclear, mitochondrial, ER, and microtubule morphology following fixation. The extent of organelle morphology preservation followed the order: methacrolein < Glyoxal < PFA < PFA + GA (Supplementary Fig. 13). Based on these results, we recommend PFA + GA as the preferred fixative for FLEXTAG applications when minimizing fixation artifact is a priority, while glyoxal or PFA alone may further reduce nonspecific binding, albeit at the cost of reduced structural preservation.

### Conventional cell fixation for immunofluorescence

Cultured U2OS cells were fixed using one of the following conditions: ⅰ) 4.0% (wt/vol) methacrolein; ⅱ) 4.0% (wt/vol) paraformaldehyde (PFA) + 0.1% glutaraldehyde (GA); ⅲ) 4.0% (wt/vol) paraformaldehyde (PFA); or ⅳ) 3.0% glyoxal (pH 4.5), in DPBS for 30 min at RT, washed three times with DPBS, and permeabilized with 0.2% (vol/vol) Triton X-100 in DPBS for 10 min. Cells were then blocked in blocking buffer containing 3.0% (wt/vol) bovine serum albumin (BSA) in DPBS for 1 hour at RT and subsequently stained with primary antibodies in blocking buffer overnight at 4 °C. After cells were washing three times with DPBS, cells were incubated with Alexa fluorophore-conjugated secondary antibodies in blocking buffer for 1 hour at RT, followed by three washes with DPBS.

### Epi-fluorescence imaging

Live-cell wide-field epi-fluorescence imaging was performed after incubating cells with 1 µM fluorescent ligand in DMEM for 1 hour at 37 °C in a 5.0% CO₂ incubator. Cells were then washed twice with DPBS and imaged in a live-cell imaging solution (ThermoFisher, A59688DJ).

Fixed-cell wide-field epi-fluorescence microscopy was performed after protective fixation and non-specific binding reduction treatments, with a final fluorescent probe concentration of 20 nM (for epi-fluorescence microscopy) and 100–500 nM (for confocal fluorescence microscopy, SIM imaging, and STED imaging) in DPBS.

Widefield epi-fluorescence imaging was performed using a home-built microscope based on an Olympus IX-73 body, equipped with a Lumencor SOLA light engine (SOLA U-nIR) and a CMOS Camera (Thorlabs, CS126MU). Objectives equipped include UPlanSApo 100×/1.40 oil, PlanApo 60×/1.40 oil (Olympus), UPlanFL N 100×/1.30 oil (Olympus), PlanApo 60×/1.4 oil (Olympus), LCPlanFl 40×/0.6 (Olympus), LCAch N 20×/0.40 (Olympus). Filter sets include 49000 Olympus IX3-FFXL Mounted, ET – DAPI, AT350/50x ET460/50m T400LP; 49001 Olympus IX3-FFXL Mounted, ET - CFP, AT436/20x ET480/40m T455LP; 49002 Olympus IX3-FFXL Mounted, ET - EGFP (FITC/Cy2), ET470/40x ET525/50m T495LPXR; 49003 Olympus IX3-FFXL Mounted, ET - EYFP, ET500/20x ET535/30m T515LPXR; 49004 Olympus IX3-FFXL Mounted, ET -

CY3/TRITC, ET545/25x ET605/70m T565lpxr; 49008 Olympus IX3-FFXL_Mounted, ET mCherry/Texas Red, ET560/40x ET630/75m T585lpxr; 49006 Olympus IX3-FFXL_Mounted, ET CY5 set, ET620/60x, ET700/75m, T660lpxr; 49007 Olympus BX3_Mounted, ET Cy7, ET710/75x ET810/90m T760lpxr.

### Single-molecule localization microscopy (SMLM) imaging

The SMLM setup was based on a Nikon Eclipse-Ti2 inverted microscope. A 405-nm (Coherent, OBIS 405 nm LX), a 488-nm laser (Coherent, Sapphire 488-500 LPX CDRH), a 560-nm laser (MPB Communications, 2RU-VFL-P-1500-560-B1R), and a 642-nm laser (MPB Communications, 2RU-VFL-P-2000-642-B1R) were introduced into the sample through the back port of the microscope. A translation stage enabled the laser beams to be shifted towards the edge of the objective, ensuring that the emerging light reached the sample at incidence angles slightly smaller than the critical angle of the glass-water interface, thereby selectively illuminating fluorophores within a few micrometers of the coverslip surface. A zt405/488/561/647/752rpc-UF2 dichroic mirror (Chroma) and a zet405/488/561/642/752m multi-band-pass emission filter (Chroma) were used for fluorescence detection.

For one-color three-dimensional (3D) SMLM imaging, a cylindrical lens (f = 1000 mm) was placed between the microscope side port and the EMCCD camera (Andor iXon Life 897, Andor Technology). This configuration elongated the images of single molecules in the *x* and *y* directions for molecules located on the proximal and distal sides of the focal plane (relative to the objective), respectively. The ellipticity of the single-molecule images was then used to determine the *z* position of the molecules.

Single-color PAINT imaging experiments were performed using U2OS cells after protective fixation, with a final fluorescent probe concentration of 0.05–1 nM in DPBS, depending on protein tags and the expression level of the target protein. For targets with high expression levels, PAINT movies consisting of 30,000 frames were acquired using continuous illumination with a 200 ms exposure time per frame, resulting in a total acquisition time of 100 min. For some target proteins with low expression levels, PAINT imaging with FLEXTAGs was performed using pulsed illumination. Each imaging cycle consisted of 1 s of recorded illumination (5 frames) followed by 4 s with no image acquisition (dark period), resulting in a total acquisition time of ∼8 hours for a 30,000-frame PAINT movie.

For three-color two-dimensional (2D) SMLM imaging, data were acquired sequentially in the following order: far-red–channel PAINT, red-channel PAINT, and far-red-channel STORM. First, 1 nM ET-JQ1-AF647 or 1 nM SLF’-HMSiR was added to image RTN4-FLEXTAG1 or RTN4-FLEXTAG3 using PAINT. After completion of far-red PAINT imaging, the sample was washed by 3 × 1 mL DPBS. Next, 1 nM TMP-TMR was added to label TOM20-FLEXTAG2, and red-channel PAINT imaging was performed under the same acquisition settings. The sample was then washed again with 3 × 1 mL DPBS prior to STORM imaging. For STORM imaging, the actin cytoskeleton was labeled by incubating cells with 0.5 µM phalloidin-AF647 for 30 min at room temperature. The sample was washed once with 1 mL DPBS to remove excess phalloidin-AF647, followed by far-red channel STORM imaging. Samples for PAINT, HMSiR-based STORM, and AF647-based STORM were imaged at laser power densities of 0.05 kW cm^-2^, 0.1 kW cm^-2^, and 2 kW cm^-2^, respectively, with an EMCCD camera gain of 200. Representative SMLM images were reconstructed from 30,000 frames with 200 ms exposure time per frame, resulting in total data acquisition time of 100 min. SMLM images for kinetic calculations were reconstructed from 10,000 frames with 100 ms exposure time per frame, resulting in total data acquisition time of 17 min.

Live-cell STORM was performed in live U2OS cells. Cells were incubated in live-cell imaging solution (Gibco, A59688DJ) containing 30 nM SLF’-HMSiR/HTL-JF630b for 10 min at RT. Prior to incubation, the SLF’-HMSiR solution was passed through a 0.22 µm filter and sonicated for 10 minutes to remove SLF’-HMSiR aggregates and ensure optimal probe performance.

### Stimulated emission depletion (STED) nanoscopy

STED nanoscopy was performed on U2OS cells following protective fixation using a Zeiss LSM 980 confocal microscope equipped with an Abberior STEDYCON add-on module. Two-color fluorescent labeling was achieved by conjugating FLEXTAG1, FLEXTAG2, or FLEXTAG3 ligands with JF635 and AF594. Excitation was performed using a 633 nm laser for JF635 and a 561 nm laser for AF594. Super-resolution was achieved using a 775 nm depletion laser for STED imaging.

### Structured illumination microscopy (SIM)

Structured illumination microscopy (SIM) was performed using a custom microscope built on an Olympus IX-83 body, equipped with the CrestOptics X-Light V3 spinning disk confocal unit in combination with DeepSIM X-light super-resolution system module. Image acquisition was carried out using an ORCA-Fusion BT Digital CMOS camera (Hamamatsu). The standard data acquisition setting was applied, where 37 raw images were captured per field of view, each with a structured illumination pattern at different phase shifts and orientations, to reconstruct a single super-resolved SIM image.

Three-color SIM imaging was performed in live U2OS cells, 100 nM fluorophores TMP-JF525, SLF’-MaP555, and ET-JQ1-JF635 were used, with excitation wavelengths of 470 nm, 555 nm, and 640 nm, respectively.

Photobleaching experiments were performed in live U2OS cells by sequentially imaging the same field of view 30 times, with 1-minute intervals between cycles to allow fluorescent probe self-renewal and potential recovery of fluorescence intensity.

### SMLM image processing and determination of event rate and apparent off-rate

SMLM image reconstruction and rendering were performed using Picasso v0.6.9. Single-molecule localizations were identified frame-by-frame using Least Squares (LQ) Gaussian fitting with following parameters: minimum net gradient of 10,000–15,000, baseline of 100, sensitivity of 1.00, and quantum efficiency of 1.00. Lateral drift was subsequently corrected using redundant cross-correlation (RCC) implemented in Picasso v0.6.9. Apparent off-rates (the average rate of disappearance of the single-molecule fluorescent spot), event rates (the average number of fluorescent ligand binding events per unit imaging area per unit time), single-molecule fitting precisions, and NeNA localization precisions, were determined from the reconstructed SMLM images using Picasso v0.6.9^5^.

### Determination of TMR/mEGFP fluorescence ratio

The TMR/mEGFP fluorescence ratio in the mitochondria of U2OS cells expressing TOM20 dually tagged with mEGFP and a FLEXTAG tag was quantified using CellProfiler to assess the relative fluorescence intensity of TMR compared to mEGFP. mEGFP was used as the reference channel to define the mitochondrial regions in the U2OS cells. For each GFP-channel fluorescence image, background subtraction was performed using the Circular Average Filter module in CellProfiler with the user-defined diameter set to 50 pixels (approximately 1.725 µm). After background subtraction, the filtered mEGFP image was thresholded based on fluorescence intensity to generate a binary mask of mitochondrial regions. The mitochondrial mask was then applied to both the TMR and mEGFP fluorescence channels, ensuring that fluorescence intensity measurements were confined to the mitochondrial regions defined by the mask. The mean fluorescence intensities within the mitochondrial mask were calculated for both the TMR and mEGFP channels. The ratio of TMR to mEGFP fluorescence intensity was subsequently computed to assess labeling efficiency of the FLEXTAG tag on mitochondria.

### Mitochondrial morphology assay

To assess the potential self-aggregation of FLEXTAG tags, cell fractions exhibiting normal or abnormal mitochondrial morphology were determined through visual inspection. Normal mitochondrial morphology was defined as elongated and variably sized filamentous structures that were non-clustered and distributed throughout the perinuclear region as well as parts of the cytosol. A dense distribution of mitochondria around the nucleus is not classified as abnormal unless accompanied by clustering.

Abnormal mitochondrial morphology was characterized by one or more of the following features: ⅰ) Mitochondria appeared clumped in localized regions; ⅱ) Mitochondria displayed excessive intertwining; ⅲ) Mitochondria were predominantly concentrated in the perinuclear region, with minimal extension into the peripheral cytoplasm.

It is important to note that abnormal mitochondrial morphology is not solely caused by the aggregation of protein tags. Other factors, such as toxicity from transient transfection reagents or cell stress due to excessive exogenous protein expression, may also contribute to abnormal mitochondria morphology. As a negative control, we measured the cell fraction exhibiting abnormal mitochondrial morphology in cells expressing mEGFP-tagged TOM20. We found that approximately 10.2 ± 2.0% of these cells displayed abnormal mitochondrial morphology.

### Mitochondrial and nuclear morphology-based cell viability assay

To assess the cytotoxicity of protective agents, we monitored cell number and morphological changes of mitochondria and the nucleus at 0, 1, 5, 12, and 24 hours following the addition of 100 µM protective agents, using epi-fluorescence microscopy. Abnormal nuclear morphology was defined by notable shrinkage or the appearance of multilobed structures, while abnormal mitochondrial morphology included fragmentation, rounding, or altered subcellular distribution compared to the typical elongated, filamentous network observed under normal conditions.

### MTT-based cell viability assay

Cell viability was measured using the MTT assay following standard protocols. Cells were seeded in 96-well plates at optimized densities and treated with HMBR/JQ1/MTX/ET-JQ1/TMP/SLF’ for 72 h at 37 °C. MTT solution was added and incubated for 4 h at 37 °C, after which formazan crystals were solubilized. Absorbance was measured using a BioTek ELx808 microplate reader, and relative viability was calculated against vehicle controls. IC_50_ values were determined by nonlinear regression analysis of dose-response curves using GraphPad Prism.

### Protein Purification

Day 1: *E. coli* (BL21) cells were transformed by heat shock (50 ng plasmid, 42 °C for 45 s), recovered in SOB for 1 h at 37 °C, and plated on LB agar containing kanamycin (50 µg/mL). Day 2: For expression, a single colony was inoculated into LB with kanamycin and grown overnight. The next day, 1.5 L LB cultures were inoculated (1:100) and grown at 37 °C until OD_600_ reached 0.6–0.8, followed by induction with 0.5 mM IPTG. Cultures were shifted to 18 °C for FLEXTAG1/FLEXTAG3, and 15 °C for FLEXTAG2, and expressed overnight (12–15 hours). Day 3: Cells were harvested by centrifugation (3000 rpm, 30 min, 4 °C) and resuspended in buffer (50 mM Tris, 250 mM NaCl, 1 mM PMSF, pH 7.4). Cell suspensions were lysed on ice by probe sonication (2 seconds on and 8 s off, total on-time 6 min, 60% amplitude). Lysates were clarified by centrifugation (16,000 rpm, 60 min, 4 °C).

The soluble lysate was then loaded to a HiTrap TALON Crude 5 mL column (Cytiva) pre-equilibrated with Buffer A (50 mM Tris, 250 mM NaCl, pH 7.4) using a Mini ÄKTA chromatography system. The column was washed with 5% imidazole in Buffer A at a flow rate of 4 mL/min to remove nonspecifically bound proteins. Bound protein was subsequently eluted over 8 column volumes using Buffer B (Buffer A supplemented with 150 mM imidazole). Elution fractions were collected automatically and analyzed by SDS–PAGE using 4–20% gradient gels run at 150 V for 50 min to identify protein-containing fractions. Fractions containing the target protein were pooled and concentrated using 3 kDa MWCO centrifugal concentrators at 3000 × g, followed by filtration through a 0.45-µm syringe filter. Protein concentration was measured on a NanoDrop spectrophotometer based on A280. Purified proteins were aliquoted, flash-frozen in liquid nitrogen, and stored at −80 °C until use. SDS-PAGE and ÄKTA chromatogram analyses of purified His-tagged FLEXTAG proteins are shown in Supplementary Data 1.

### Determination of binding affinities for FLEXTAG1-3

Isothermal titration calorimetry (ITC) experiments were performed on TA Affinity ITC auto instrument using ITCRun software version 3.8.4.24000 (TA instrument). The stir speed was 125 rpm at 20 °C. Small-molecule ligands and purified FLEXTAGs are prepared in buffer (50 mM Tris, 250 mM NaCl, pH 7.4) containing 1% DMSO. Matching 50 mM Tris, 250 mM NaCl, pH 7.4 + 1% DMSO buffer was used for all blank and reference titrations. Ligand concentrations were maintained at least 5-fold higher than protein concentrations to ensure proper saturation behavior. A 96-well sample plate was prepared with 485 μL material for each cell solution and 220 μL for each syringe solution, and the plate was covered to prevent evaporation. For each experiment, 25 injections of 2.0 μL volume were performed. Data analysis was performed using NanoAnalyze software version 4.0.2.0. The thermograms were fit using an independent model. (TA instrument, https://www.tainstruments.com/support/software-downloads-support/downloads/).

We also performed a cell-imaging-based fluorescence enhancement assay, in which H2B-mEGFP-FLEXTAG fusion proteins were overexpressed in U2OS cells and labeled with a concentration series (1, 5, 25, 125, 625, and 3125 nM) of ET-JQ1-JF635, TMP-JF635, or SLF’-JF635. By plotting the JF635/mEGFP fluorescence ratio as a function of ligand concentration, we derived apparent cellular *K_D_* values.

### ColabFold predictions

The dimer interface was predicted using AlphaFold2 (ColabFold v1.5.5) with MMseqs2 for multiple sequence alignment (MSA) generation. The ColabFold notebook (https://colab.research.google.com/github/sokrypton/ColabFold/blob/main/AlphaFold2.ipynb) was used with default settings. The model confidence was evaluated using the predicted local distance difference test (pLDDT) scores provided by AlphaFold2. Structural visualization and analysis were performed using PyMOL 2.5.4.

To identify potential residue interactions at the dimer interface, inter-protein contacts were identified by detecting any atomic pair with bond distances ≤ 3.0 Å or 3.5 Å.

The predicted structures of BRD4BD2 variants, eDHFR variants, and FKBP variants were aligned to the crystal structure of BRD4 complexed with JQ1 (PDB: 3MXF), eDHFR complexed with TMP (PDB: 6XG5), and FKBP^F36V^ complexed with SLF’ (PDB: 1BL4), respectively, to assess structural changes with designed mutations.

## Data availability

All data supporting the findings of this study are provided within the paper and its Supplementary Information. Source data of all data presented in graphs within the figures are provided with this paper.

## Code availability

The following software was used in this study: PyMOL 2.5.4, CellProfiler 4.2.5, and Picasso v0.6.9. Other custom MATLAB codes used in this study are available from the corresponding authors upon reasonable request.

## Acknowledgements

This work was supported by the National Institute of General Medical Sciences of the National Institutes of Health (R35GM142973 to R.Z. and R35GM154931 to M.F.), the Life Sciences Research Foundation, and startup funding from the Pennsylvania State University provided to R.Z. We thank Zeyu Xi for helpful discussion regarding HMSiR imaging. We are grateful to Dr. Luke D. Lavis for generously providing HTL-JF630b as a gift. We also acknowledge the Proteomics and Mass Spectrometry Core Facility of the Huck Institutes of the Life Sciences (RRID:SCR_024462) for access to the Bruker UltrafleXtreme MALDI-TOF/TOF instrument, and the Huck Institutes’ Biomolecular Interactions Core Facility (RRID:SCR_024458) for access to TA Affinity ITC auto instrument.

## Author contributions

R.Z. and H.Z. conceived the project, designed the experiments, interpreted the data, and wrote the manuscript. H.Z. and Y.W synthesized the fluorescent ligands. H.Z. and Y. Z. expressed and purified FLEXTAG proteins. H.Z. and S.Z. conducted the ITC experiments. S.Z. conducted the MTT assay. H.Z. and X.W. prepared imaging samples, performed conventional fluorescence imaging and SMLM imaging experiments. H.Z. performed SIM imaging experiments. H.Z., Y.Y., and M.F. designed and conducted STED imaging experiments. H.Z., Y.T., H.Y., and M.L. developed the software used for data acquisition and analysis. H.Z., X.W., Y.T., and H.Y. analyzed the data. R.Z. acquired funding and supervised the project.

## Competing interests

R.Z. and H.Z. are co-inventors on a patent application related to this work (US Provisional Patent App No.63/836,663). All other authors declare no competing interests.

## Supplementary Information for

### Supplementary Figures

**Supplementary Fig. 1.**
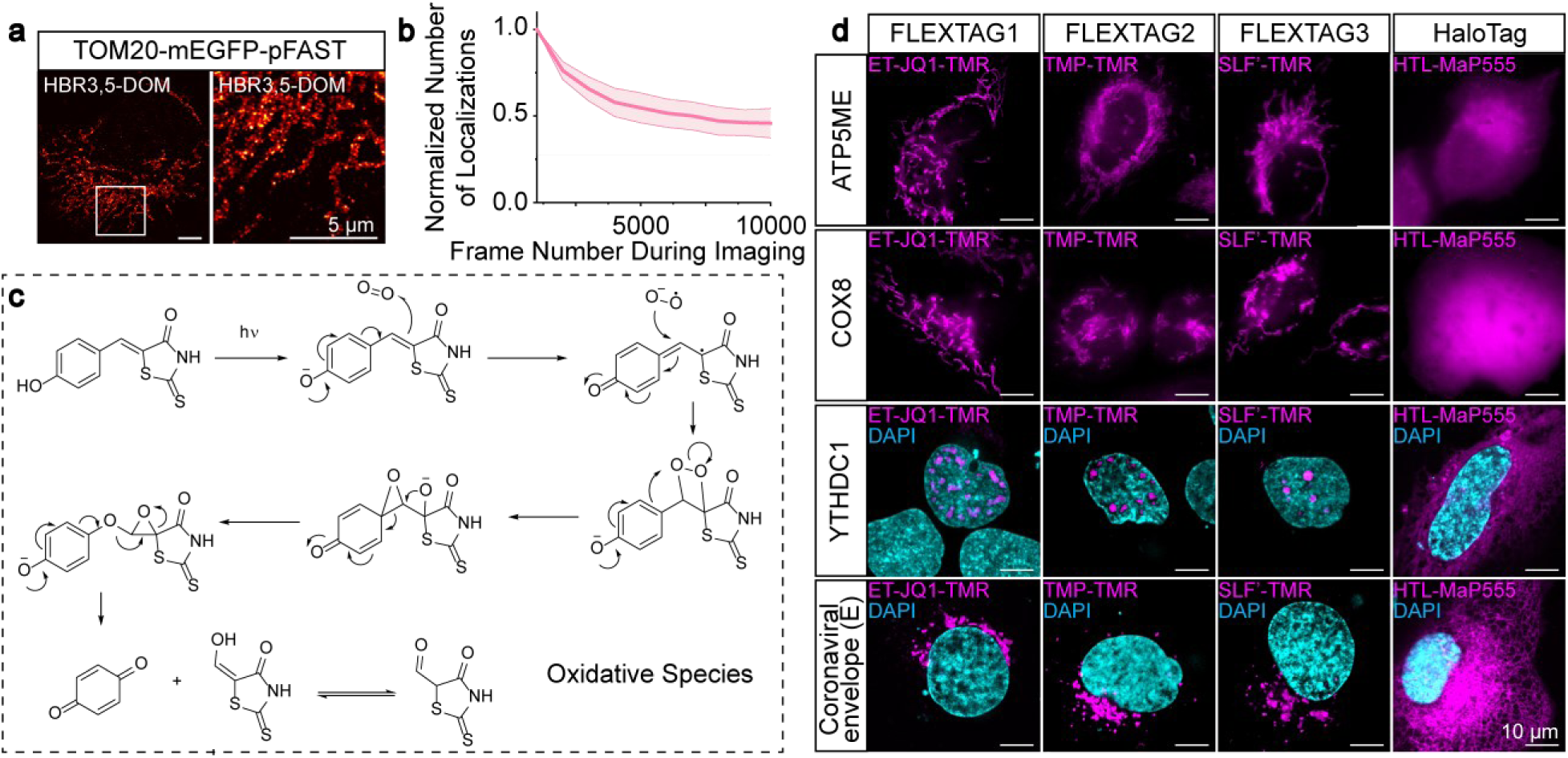
Limitations of GFP-chromophore-based PAINT imaging and large protein tags such as HaloTag. **a,** Representative PAINT images of fixed cells overexpressing TOM20-mEGFP-pFAST, imaged with 5 nM HBR-3,5DOM^1^. The image on the right is a magnified view of the boxed region in the left image. Scale bars: 5 μm. **b,** Quantification of survival rate of the single-molecule localizations during 30,000-frame PAINT imaging of 5 nM HBR-3,5DOM. Localization numbers are normalized to the first frame. fields of viewData are presented as mean ± s.e.m. *n* = 3 biological replicates; 3 fields of view (FOVs) were examined. **c,** Proposed photobleaching mechanism of GFP-chromophores (e.g., HBR derivatives)^2^. The photobleaching product quinone and aldehyde (enol) may react with pFAST and cause irreversible binding pocket blockage. **d,** Representative epi-fluorescence images showing morphology of ATP5ME, COX8, and confocal fluorescence microscopy showing morphology of YTHDC1, and the coronaviral envelope (E) protein when fused to FLEXTAG1, FLEXAG2, FLEXTAG3, or HaloTag^3^. ATP5ME, COX8, YTHDC1, and the coronaviral envelope (E) protein are expected to localize to the mitochondria, mitochondria, nucleus, Golgi apparatus, respectively. Scale bars: 10 μm.

**Supplementary Fig. 2.**
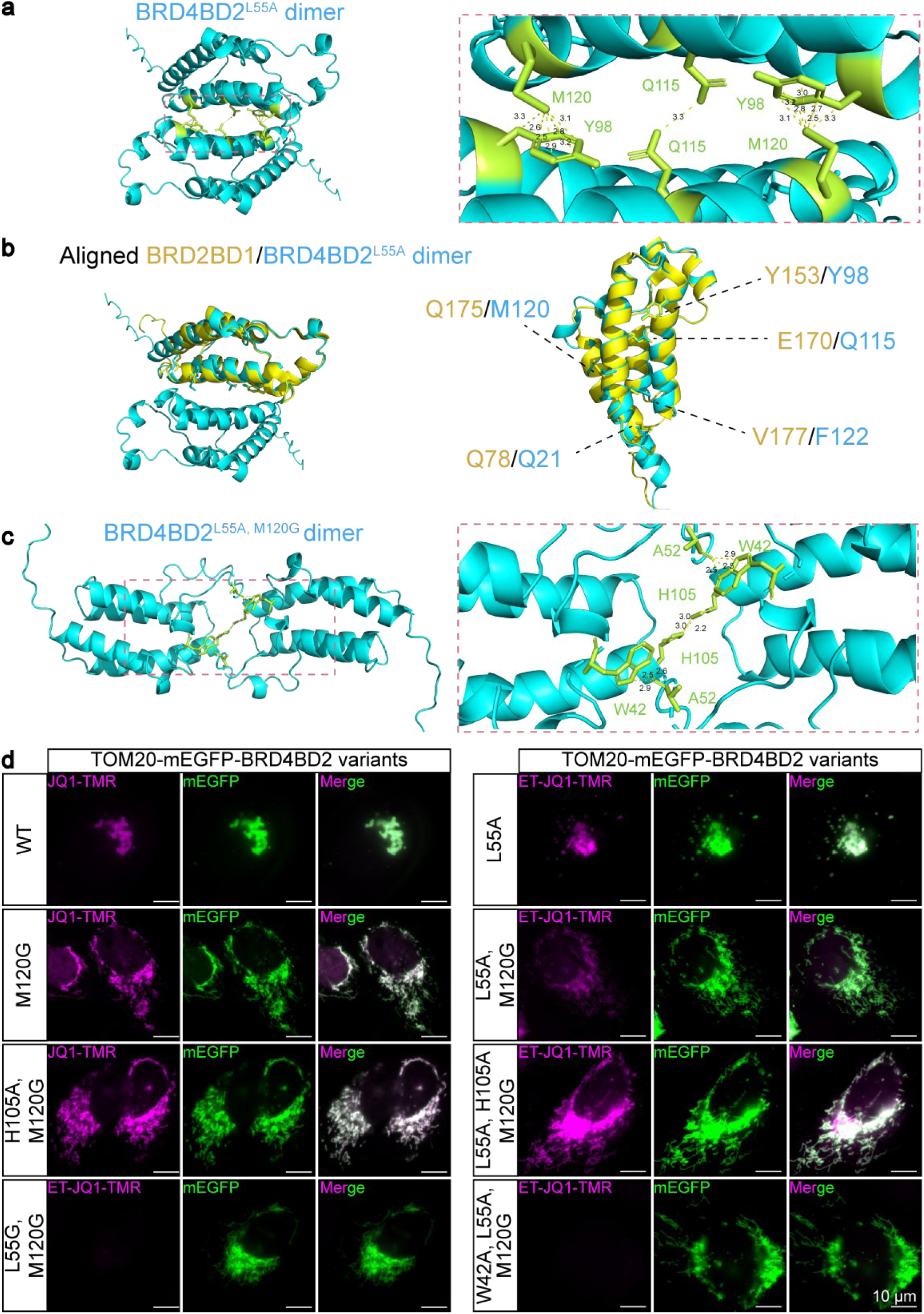
ColabFold-predicted dimer structures of BRD4BD2 variants and analysis of mitochondrial morphology in cells overexpressing TOM20-mEGFP-BRD4BD2 variants. **a,** ColabFold^4^ prediction of BRD4BD2^L55A^ dime interfaces. Key inter-protein interactions (≤ 3.5 Å bond distances) are highlighted in green, including Y98-M120 and Q115-Q115. Residue-residue interactions are depicted as yellow dashed lines, with interatomic distances measured in PyMOL and reported in angstroms (Å). **b,** Structural alignment of the ColabFold-predicted BRD4BD2^L55A^ and the crystal structure of BRD2BD1 (PDB: 3JVJ)^5^. **c,** ColabFold prediction of BRD4BD2^L55A, M120G^ dimer interfaces. Key inter-protein interactions (≤ 3.5 Å bond distances) are highlighted in green, including A52-W42 and H105-H105. Residue-residue interactions are depicted as yellow dashed lines, with interatomic distances measured in PyMOL and reported in angstroms (Å). **d,** Representative epi-fluorescence images of fixed cells overexpressing TOM20-mEGFP-BRD4BD2 variants. BRD4BD2 variants with L55A or L55G mutations were labeled with 1 µM ET-JQ1-TMR, while variants without the L55 mutation were labeled with 1 µM JQ1-TMR. Scale bars: 10 μm.

**Supplementary Fig. 3.**
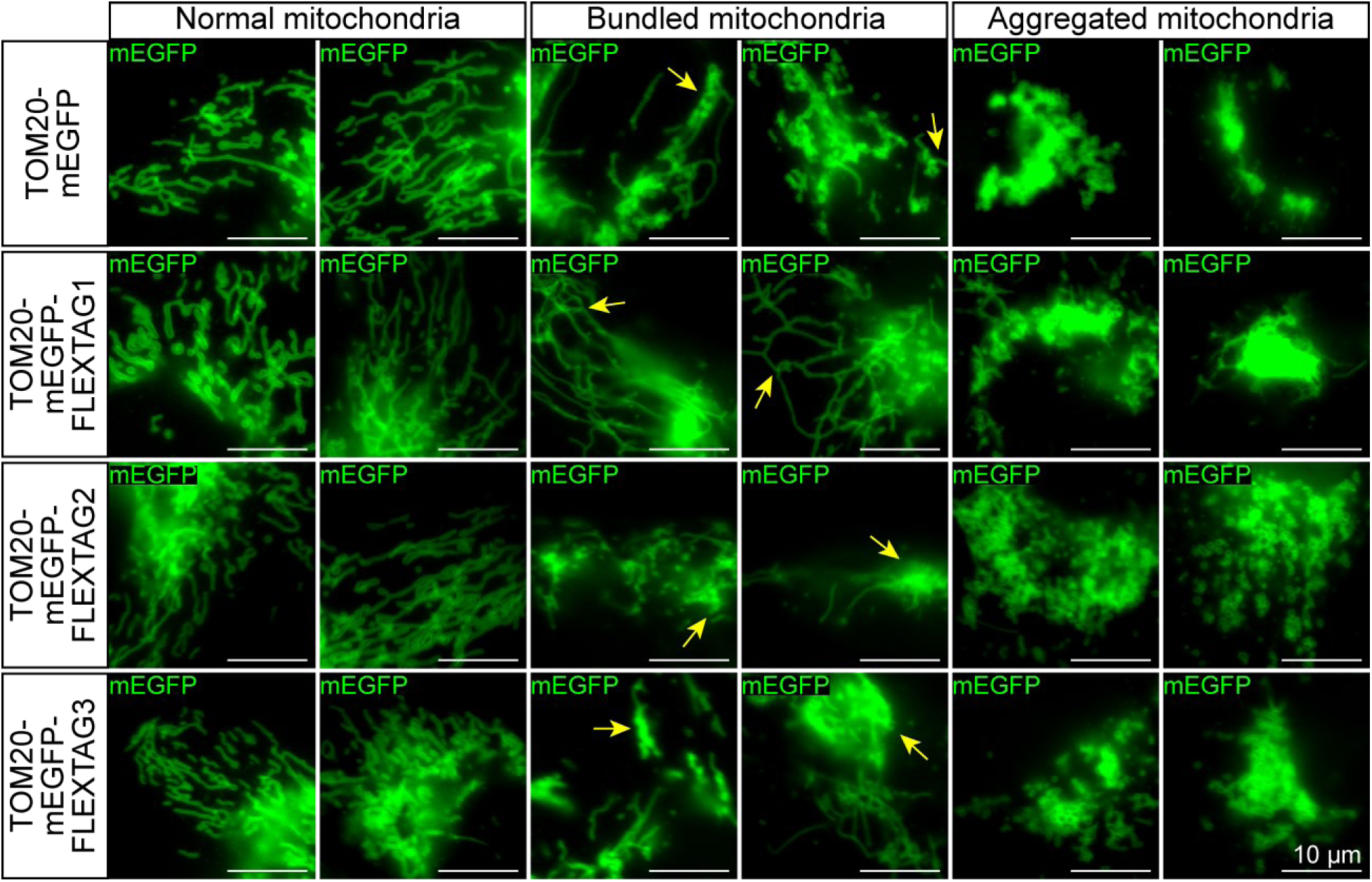
Mitochondrial morphology assay. Representative epi-fluorescence images showing normal, bundled, and aggregated mitochondria in cells overexpressing TOM20-mEGFP (control), TOM20-mEGFP-FLEXTAG1, TOM20-mEGFP-FLEXTAG2, or TOM20-mEGFP-FLEXTAG3. Abnormal mitochondrial morphology, classified as either bundled or aggregated, was defined by one or more of the following features: ⅰ) mitochondria appeared clumped in localized regions; ⅱ) mitochondria displayed excessive intertwining as indicated by yellow arrows; ⅲ) Mitochondria were predominantly clustered around the nucleus, with minimal extension into peripheral cytoplasmic regions. Scale bars: 10 μm.

**Supplementary Fig. 4.**
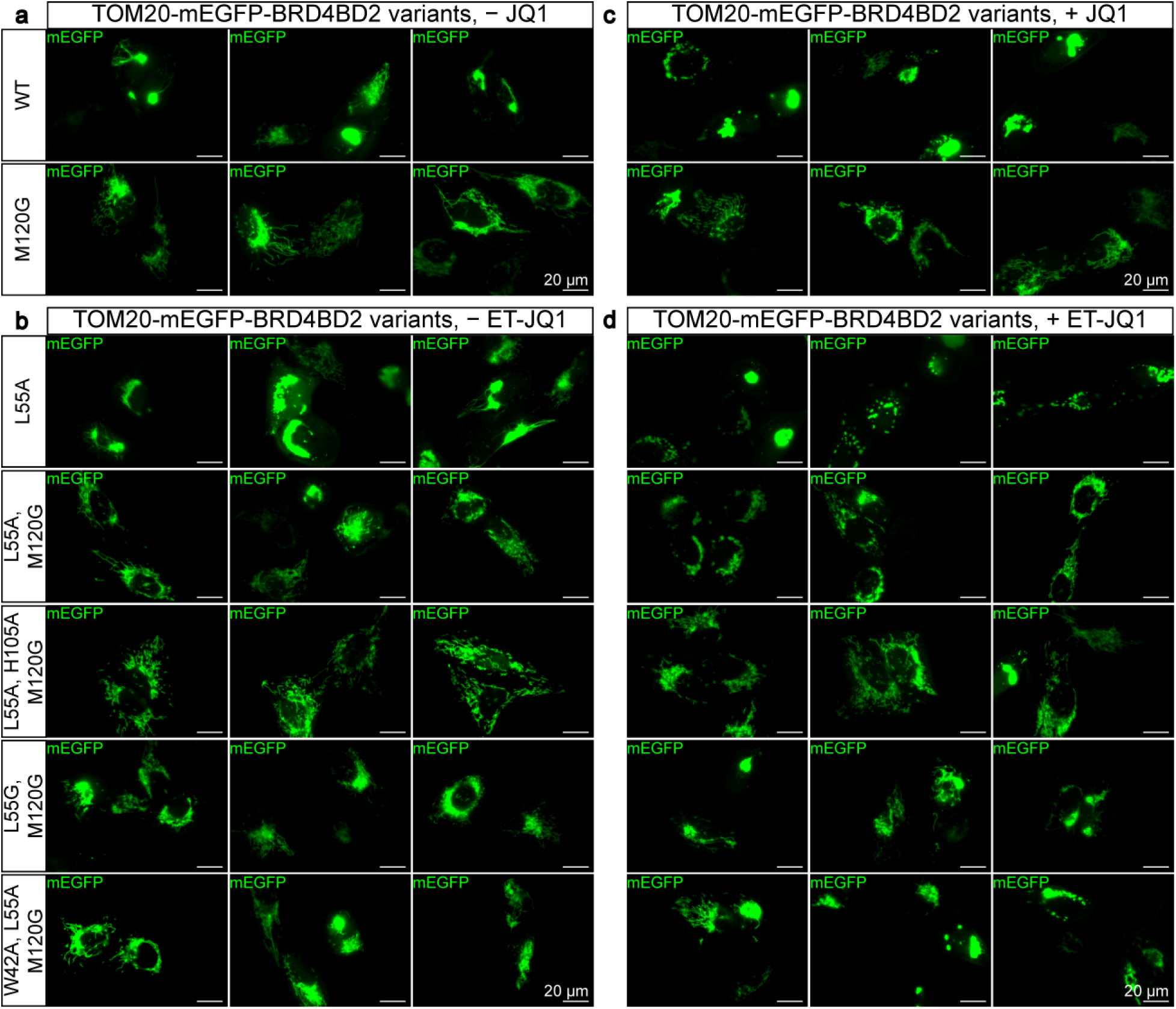
Analysis of mitochondrial morphology in cells overexpressing TOM20-mEGFP-BRD4BD2 variants. **a,** Representative epi-fluorescence images showing mitochondrial morphology of cells overexpressing TOM20-mEGFP-BRD4BD2 variants with the indicated mutations in the absence of ligandJQ1 or ET-JQ1. Scale bars: 20 μm. **b,** Representative epi-fluorescence images showing mitochondrial morphology of cells overexpressing TOM20-mEGFP-BRD4BD2 variants with the indicated mutations in the absence of ligand ET-JQ1. Scale bars: 20 μm. **c,** Same as **a**, but cells were imaged 1 hour following the addition of 100 µM JQ1. Scale bars: 20 μm. **d,** Same as **b**, but cells were imaged 1 hour following the addition of 100 µM ET-JQ1. Scale bars: 20 μm.

**Supplementary Fig. 5.**
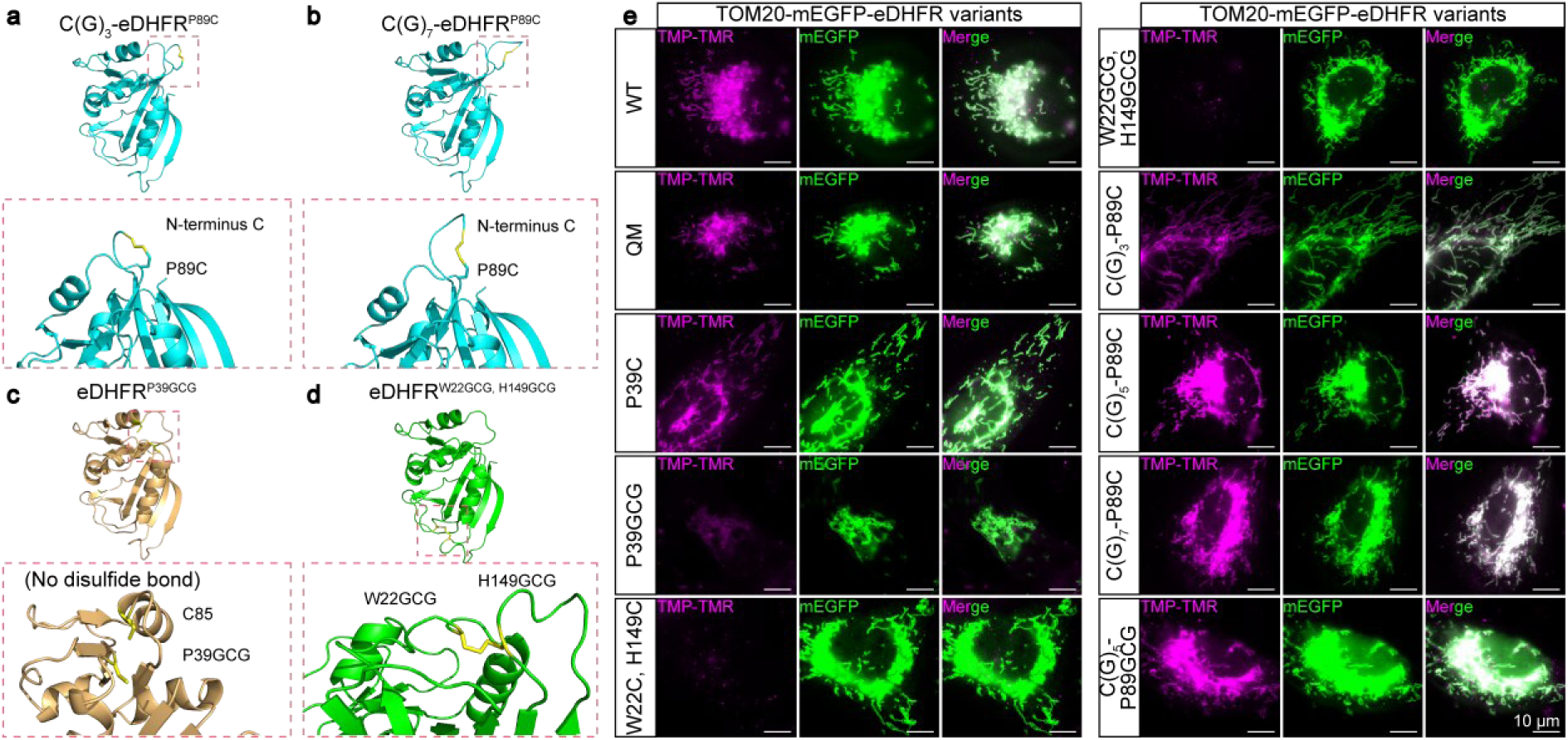
ColabFold-predicted structures and labeling efficiencies of eDHFR variants. **a,** ColabFold-predicted tructure of C(G)_3_-eDHFR^P89C^, showing a disulfide bond (highlighted in yellow) formed between the N-terminus cysteine and P89C. **b,** Same as **a**, but for C(G)_7_-eDHFR^P89C^. **c,** ColabFold-predicted structure of eDHFR^P39GCG^, showing no disulfide bond formed between P39GCG and C85. **d,** ColabFold-predicted structure of eDHFR^W22GCG, H149GCG^, showing a disulfide bond (highlighted in yellow) formed between the W22GCG and H149GCG. **e,** Representative epi-fluorescence images of cells overexpressing the indicated TOM20-mEGFP-eDHFR variants, labeled with 1 µM TMP-TMR. Scale bars: 10 μm.

**Supplementary Fig. 6.**
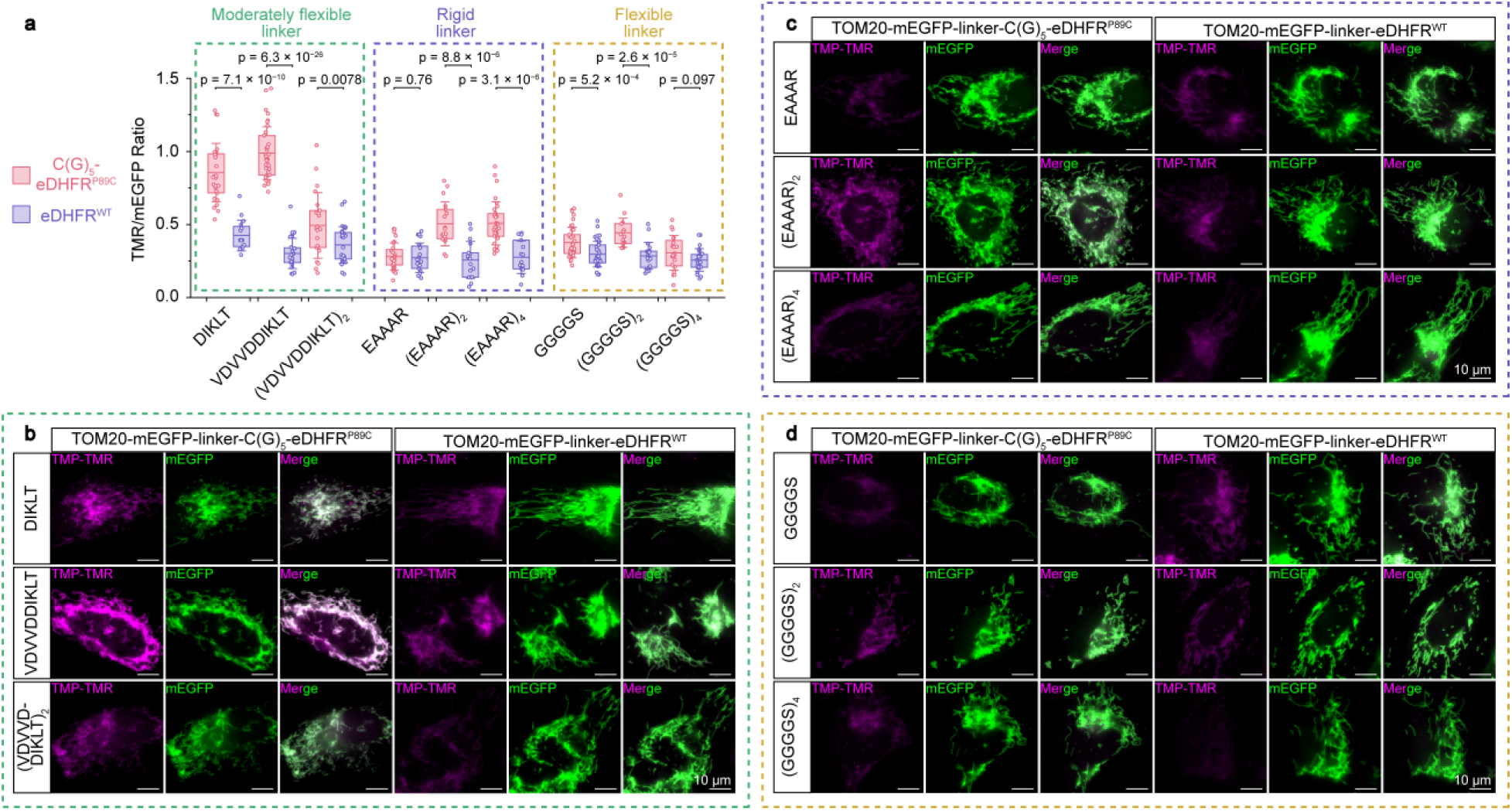
Screening of linkers to improve labeling efficiency on eDHFR variants. **a,** Quantification of the TMR/mEGFP fluorescence ratio upon TMP-TMR labeling of mEGFP-linker-C(G)_5_-eDHFR^P89C^/eDHFR^WT^. Linkers tested include moderately flexible (green boxes), rigid (purple boxes), and flexible (yellow boxes) types. Each data point represents the average TMR/mEGFP ratio of mEGFP-masked regions in a single FOV. *n* = 15–40 FOVs per condition were examined. Data are presented as mean ± s.e.m. Boxplots show the mean and boundaries (first and third quartiles); whiskers denote s.d. *p*-values calculated with two-sided unpaired Student’s *t*-test. **b,** Representative epi-fluorescence images of cells overexpressing TOM20-mEGFP-(moderately flexible linker)-eDHFR^P89C^/eDHFR^WT^, labeled by 1 µM TMP-TMR. Scale bars: 10 μm. **c,** Same as **b**, but for TOM20-mEGFP-(rigid linker)-eDHFR^P89C^/eDHFR^WT^. Scale bars: 10 μm. **d,** Same as **b**, but for TOM20-mEGFP-(flexible linker)-eDHFR^P89C^/eDHFR^WT^. Scale bars: 10 μm.

**Supplementary Fig. 7.**
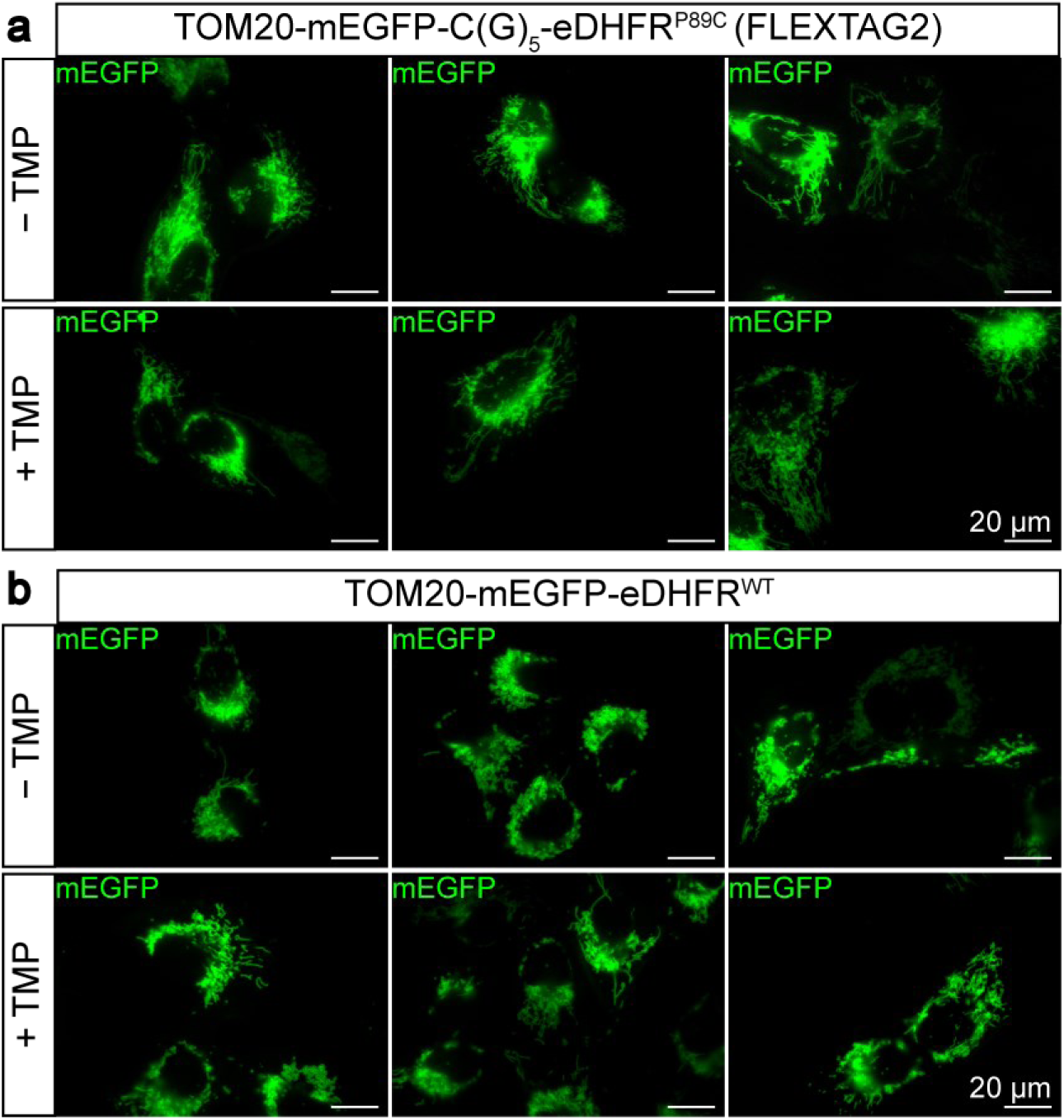
Analysis of mitochondrial morphology in cells overexpressing TOM20-mEGFP-eDHFR variants. **a,** Representative epi-fluorescence images showing the mitochondrial morphology of TOM20-mEGFP-C(G)_5_-eDHFR^P89C^ in the absence of or 1 hour after the addition of 100 µM TMP. Scale bars: 20 μm. **b,** Same as **a**, but for TOM20-mEGFP-eDHFR^WT^ Scale bars: 20 μm.

**Supplementary Fig. 8.**
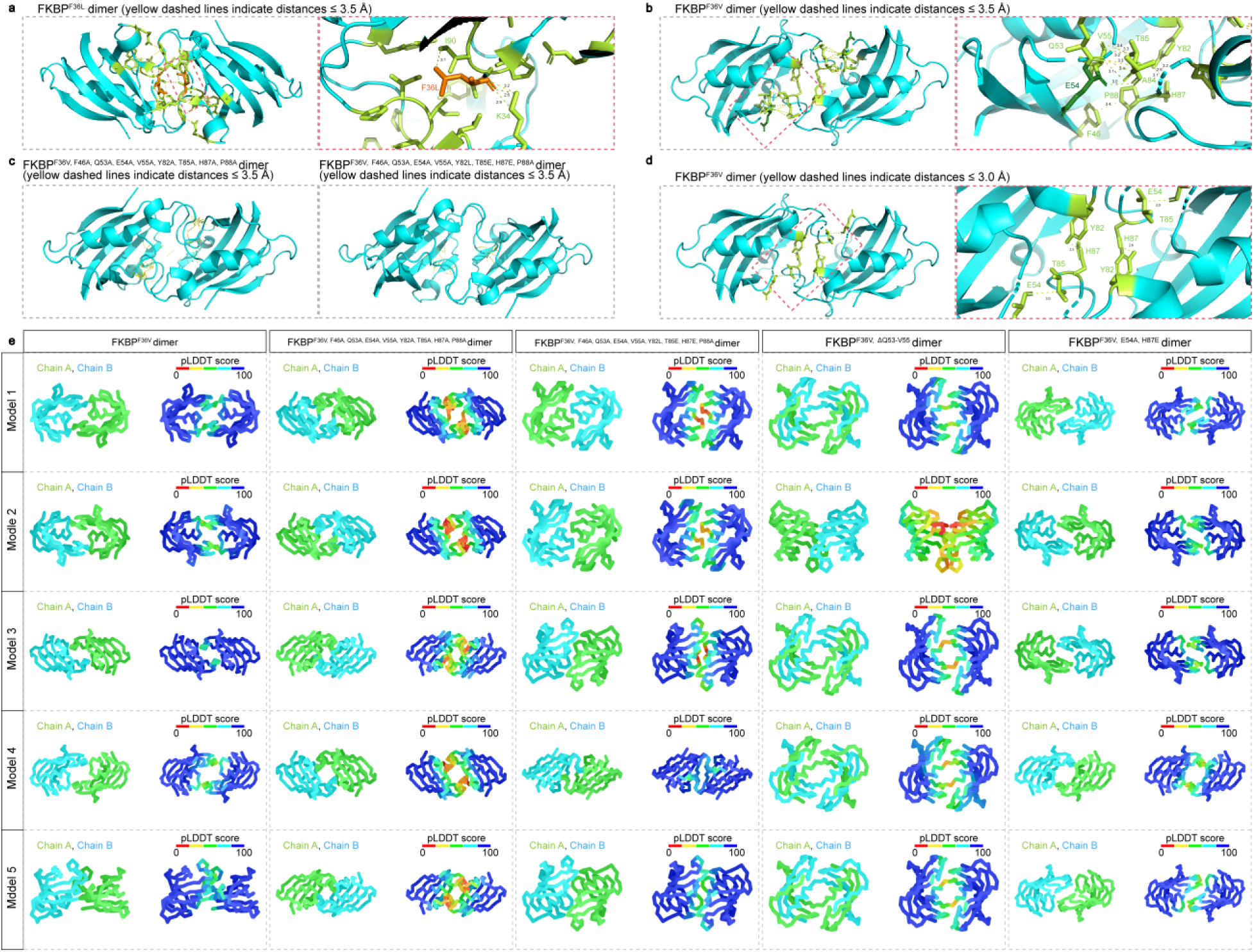
ColabFold-predicted dimer structures of FKBP variants. **a,** ColabFold prediction of the dimer interface for FKBP^F36L^. Interatomic distances ≤ 3.5 Å are shown as yellow dashed lines, including interactions between Y26-G89, K34-L36, K34-D37, L36-I90, D37-P92, R42-P92, F46-P88, Y82-I90, and I90-I91. Residues involved in dimer interface formation are highlighted in green. The F36L mutation is positioned near K34 and I90, directly contributing to dimer stabilization. **b,** ColabFold prediction of the dimer interface for FKBP^F36V^. Interatomic distances ≤ 3.5 Å are shown as yellow dashed lines, with distances measured in PyMOL. Residues involved in dimer interface formation are highlighted in green, including F46-P88, Q53/E54/V55-T85, and Y82-H87. F36V is not directly involved in the dimer interface. **c,** ColabFold prediction of the dimer interfaces for FKBP^F36V, F46A, Q53A, E54A, V55A, Y82A, T85A, H87A, P88A^ (left) and FKBP^F36V, F46A, Q53A, E54A, V55A, Y82L, T85E, H87E, P88A^ (right). Alanine (A) mutations were introduced to disrupt key interactions by reducing the size of side chains, while substitutions to glutamic acid (E) or leucine (L) were designed to weaken electrostatic interactions between oppositely charged side chains at the key positions on the dimer interfaces. Interatomic distances ≤ 3.5 Å are shown as yellow dashed lines. **d,** ColabFold prediction of the dimer interface for FKBP^F36V^. Interatomic distances ≤ 3.0 Å are shown as yellow dashed lines, including the interaction between E54 side chain and T85 backbone, and the side chain interactions between Y82 and H87. **e,** ColabFold-predicted dimer structures of five rationally designed FKBP^F36V^ variants, showing mutations intended to disrupt the FKBP^F36V^dimer interface lead to folding instability near the binding pocket of FKBP^F36V^. Variants including FKBP^F36V^, FKBP^F36V, F46A, Q53A, E54A, V55A, Y82A, T85A, H87A, P88A^, FKBP^F36V, F46A, Q53A, E54A, V55A, Y82L, T85E, H87E, P88A^ and FKBP^F36V, E54A, H87E^ retained dimerization tendencies. FKBP^F36V, F46A, Q53A, E54A, V55A, Y82A, T85A, H87A, P88A^ and FKBP^F36V, F46A, Q53A, E54A, V55A, Y82L, T85E, H87E, P88A^ exhibited compromised folding, indicated by reduced pLDDT scores in specific regions. The deletion mutant FKBP^F36V, ΔQ53–V55^ failed to fold correctly, evidenced by intertwined Chain A and Chain B.

**Supplementary Fig. 9.**
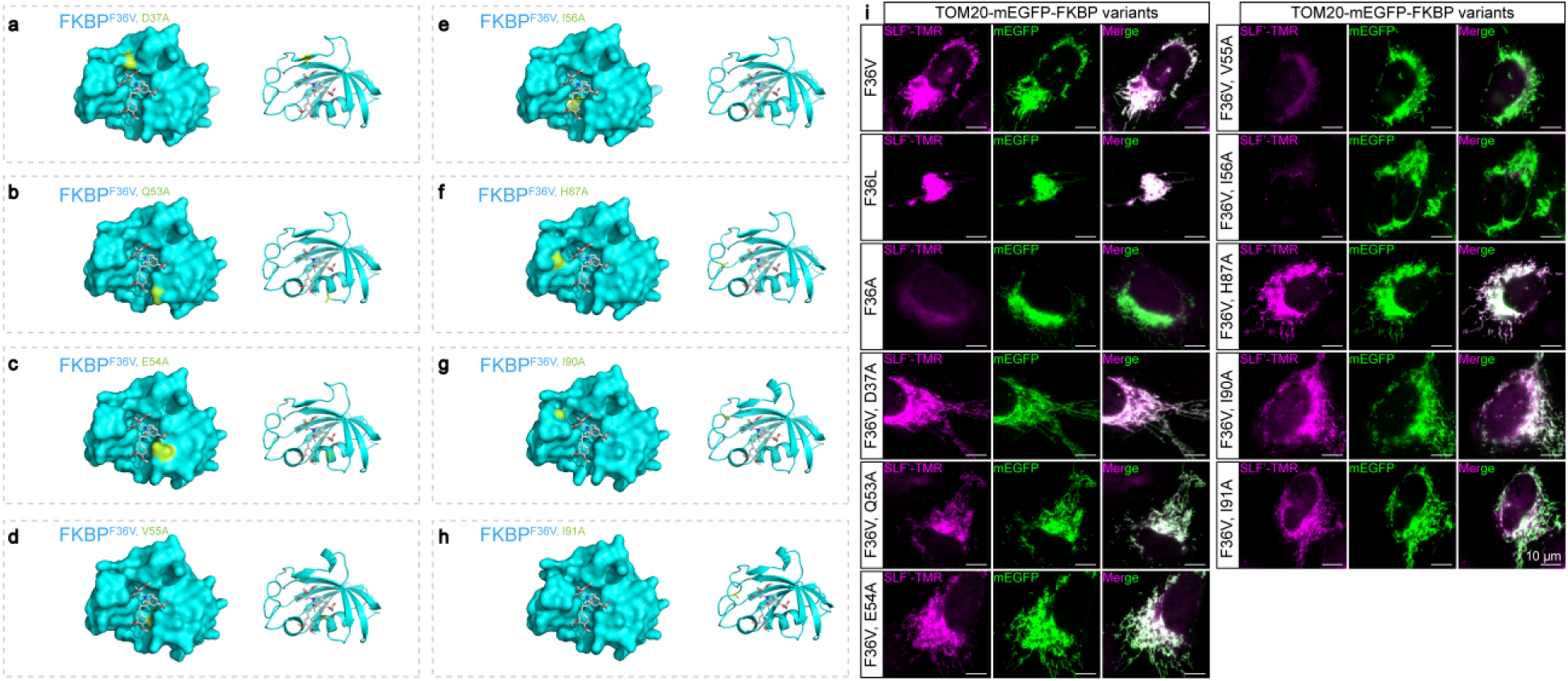
ColabFold-predicted structures and labeling efficiencies of FKBP variants. **a,** ColabFold prediction of FKBP^F36V, D37A^ structure. D37A is highlighted in green. **b,** ColabFold prediction of FKBP^F36V, Q53A^ structure. Q53A is highlighted in green. **c,** ColabFold prediction of FKBP^F36V, E54A^ structure. E54A is highlighted in green. **d,** ColabFold prediction of FKBP^F36V, V55A^ structure. V55A is highlighted in green. **e,** ColabFold prediction of FKBP^F36V, I56A^ structure. I56A is highlighted in green. **f,** ColabFold prediction of FKBP^F36V, H87A^ structure. H87A is highlighted in green. **g,** ColabFold prediction of FKBP^F36V, I90A^ structure. I90A is highlighted in green. **h,** ColabFold prediction of FKBP^F36V, I91A^ structure. I91A is highlighted in green. **i,** Representative epi-fluorescence images of fixed cells overexpressing TOM20-mEGFP-FKBP variants, labeled with 1 µM SLF’-TMR. Scale bars: 10 μm.

**Supplementary Fig. 10.**
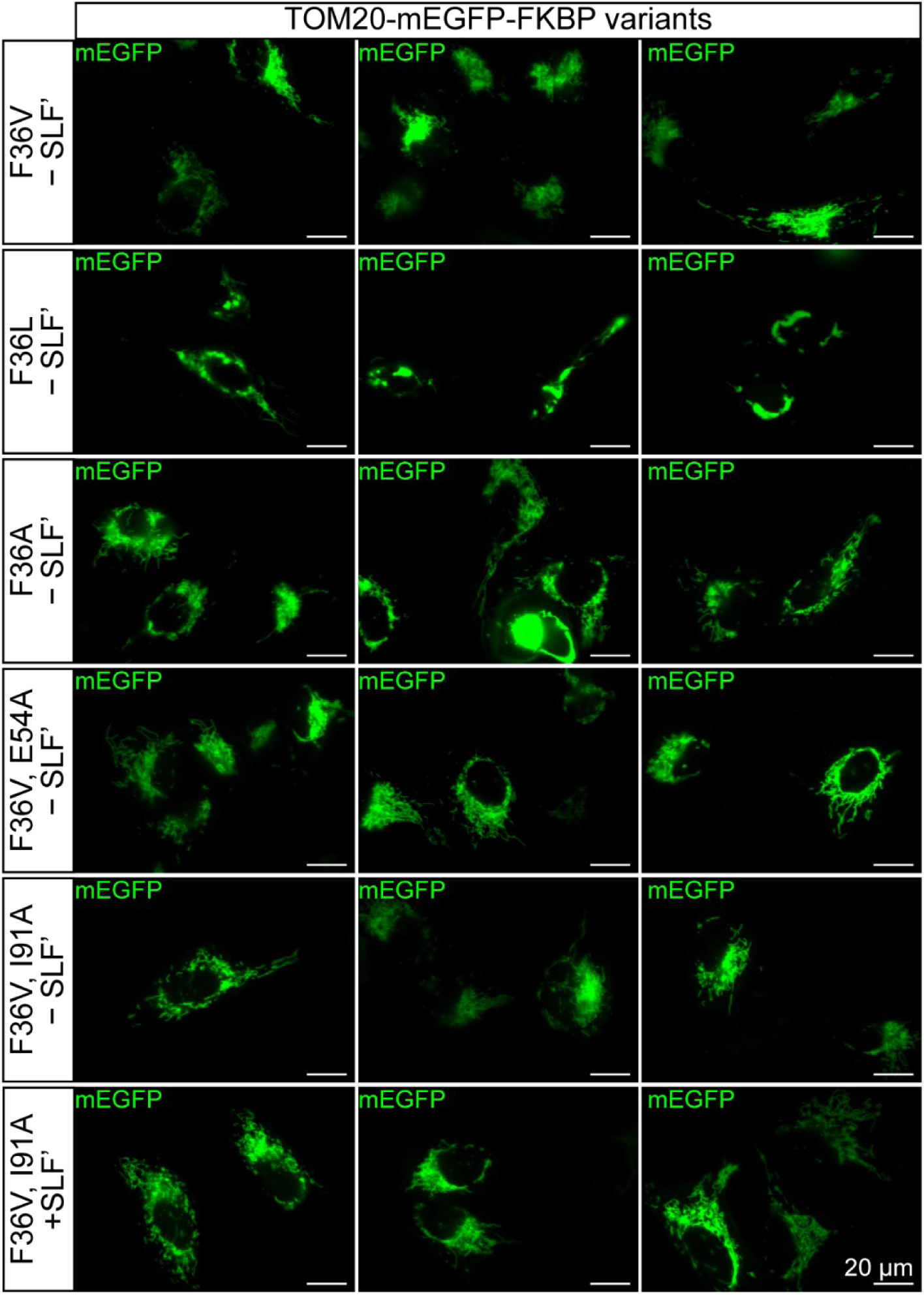
Analysis of mitochondrial morphology in cells overexpressing TOM20-mEGFP-FKBP variants. Mitochondrial morphology was assessed in cells overexpressing TOM20-mEGFP-FKBP^F36V, I91A^ (FLEXTAG3) in absence or 1 hour after treatment with 100 µM SLF’. All other FKBP variants were analyzed only in the absence of SLF’. Scale bars: 20 μm.

**Supplementary Fig. 11.**
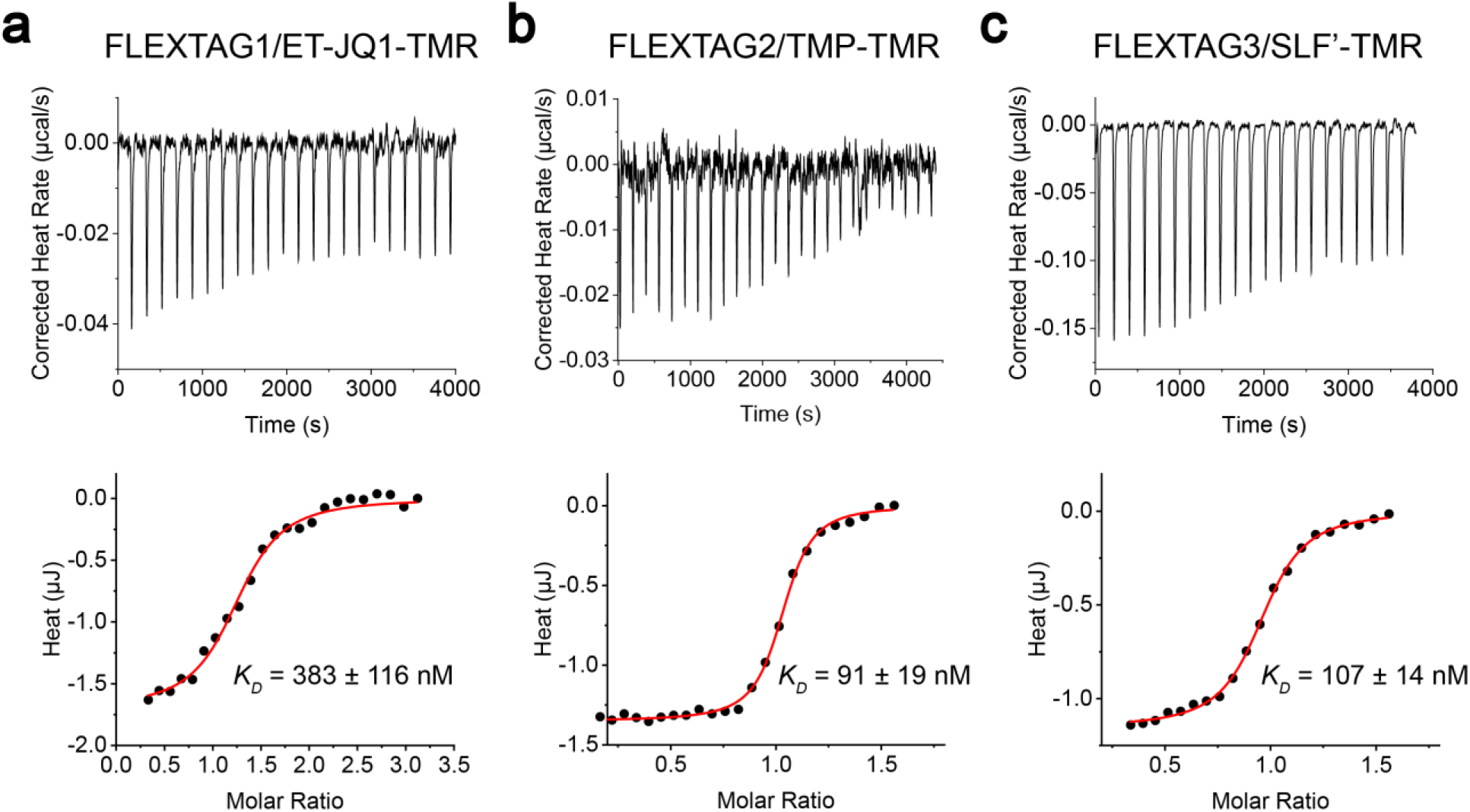
Isothermal titration calorimetry (ITC) analysis for determination of apparent. ***K_D_*** values of FLEXTAG-ligand interactions *in vitro*. a, Representative ITC thermograms (top) and corresponding binding isotherms (bottom) for FLEXTAG1. 5 µM ET-JQ1-TMR in the sample cells was titrated with 100 μM FLEXTAG1 at 20 °C. **b,** Representative ITC thermograms (top) and corresponding binding isotherms (bottom) for FLEXTAG2. 15 µM TMP-TMR in the sample cells was titrated with 100 μM FLEXTAG2 at 20 °C. **c,** Representative ITC thermograms (top) and corresponding binding isotherms (bottom) for FLEXTAG2. 10 µM FLEXTAG3 in the sample cells was titrated with 100 μM SLF’-TMR at 20 °C.

**Supplementary Fig. 12.**
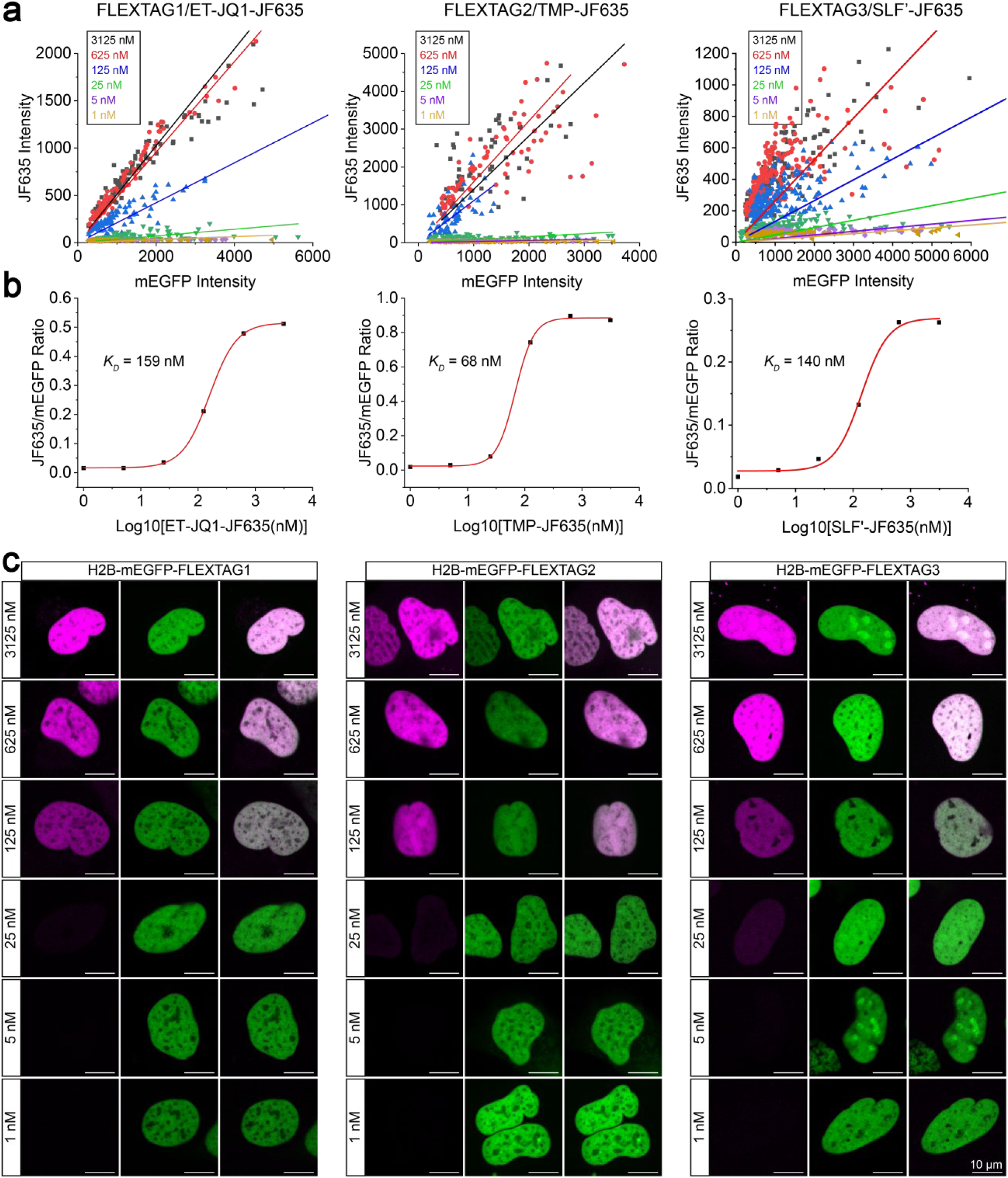
Cell-imaging-based fluorescence enhancement assay for determination of apparent. ***K_D_*** values of FLEXTAG-ligand interactions *in vivo*. a, Scatter plots of JF635 fluorescence intensity versus mEGFP fluorescence intensity in the nuclei of individual U2OS cells expressing H2B-mEGFP-FLEXTAG1, H2B-mEGFP-FLEXTAG2, and H2B-mEGFP-FLEXTAG3, labeled with increasing concentrations (1, 5, 25, 125, 625, 3125 nM) of ET-JQ1-JF635 (*n* = 85–152 nuclei), TMP-JF635 (*n* = 56–84 nuclei), or SLF’-JF635 (*n* = 116–212 nuclei), respectively. For each ligand concentration, the JF635 and mEGFP fluorescence signals were quantified within the mEGFP-defined nuclear mask, and theaverage JF635/mEGFP ratio were obtained by linear fitting of the scatter plots. **b,** Binding curves of the three FLEXTAGs. Apparent *K_D_* values were obtained by plotting the average JF635/mEGFP ratios determined from **a** against ligand concentration and fitting this does-response cruves to a nonlinear regression model. The resulting *K_D_* values for FLEXTAG1–3 were 159 nM, 68 nM, and 140 nM, respectively. **c,** Representative confocal fluorescence images of H2B-mEGFP-FLEXTAGs labeled with JF635-conjugated ligands at increasing ligand concentrations. Scale bars, 10 μm.

**Supplementary Fig. 13.**
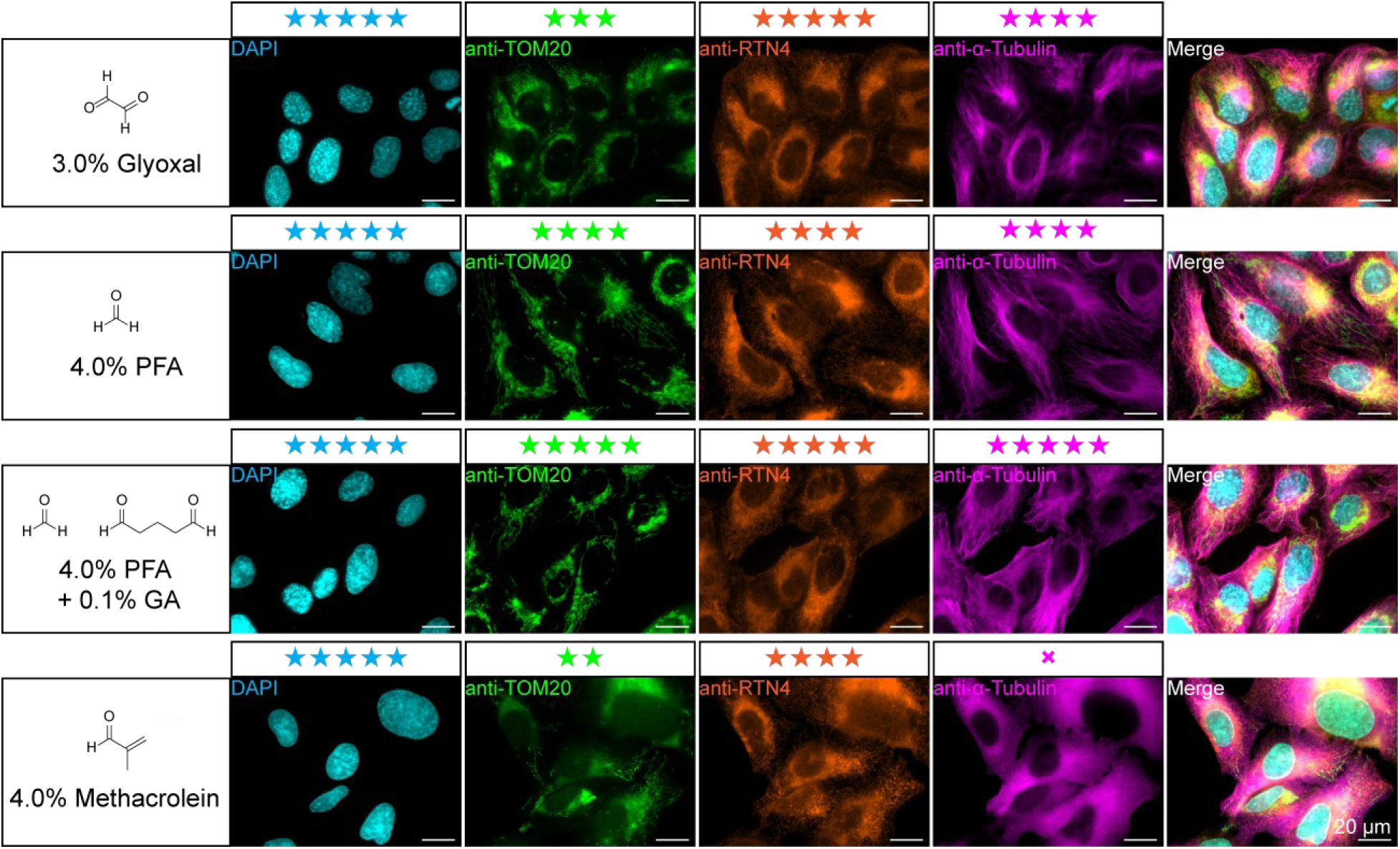
Evaluation of fixation efficiency of aldehyde-based fixatives across multiple organelles. Efficiency was assessed based on a comprehensive evaluation on immunofluorescence signal intensity, degree of signal diffusion, and the morphological preservation of the fixed organelles. The number of stars denotes the relative effectiveness of each fixation condition, Glyoxal fixation resulted in diffused mitochondrial signals, suggesting partial dissociation of TOM20 from its native localization during fixation. PFA-fixed samples showed discontinuous labeling of both mitochondria and the ER in some regions. Methacrolein fixation resulted in widespread signal diffusion for TOM20, RTN4, and α-tubulin, indicating poor structural preservation. In contrast, PFA + GA fixation yielded consistently preserved morphology and yielded strong, well-localized signals across all tested organelles. Scale bars: 20 μm.

**Supplementary Fig. 14.**
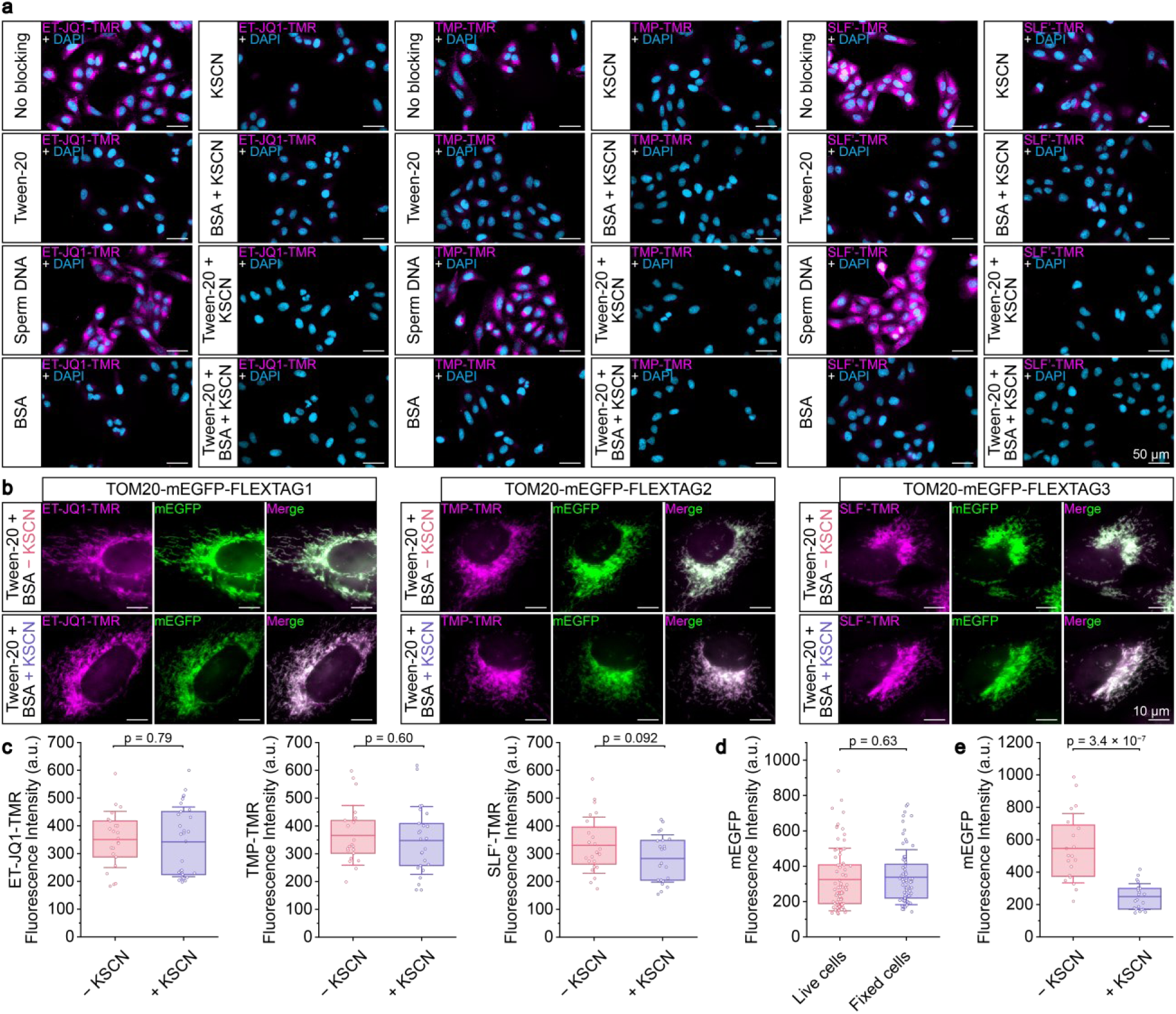
Effects of blocking agents on non-specific binding and specific labeling efficiencies. **a,** Representative epi-fluorescence images showing non-specific binding of FLEXTAG ligands in the presence of various blocking agents. Scale bars: 50 μm. **b,** Representative epi-fluorescence images of fixed cells overexpressing TOM20-mEGFP-FLEXTAG1/FLEXTAG2/FLEXTAG3, incubated with the blocking buffers containing 0.1% Tween-20 + 3.0% BSA, or 0.1% Tween-20 + 3.0% BSA + 300 mM KSCN, and labeled with the indicated FLEXTAG ligand. Scale bars: 10 μm. **c,** Effect of KSCN on the labeling efficiencies of FLEXTAG ligands. Fixed cells overexpressing TOM20-mEGFP-FLEXTAG1/FLEXTAG2/FLEXTAG3 were labeled with the corresponding TMR conjugated FLEXTAG ligand. TMR fluorescence intensities in fixed cells were quantified after incubation and imaging in buffers containing either 0.1% Tween-20 + 3.0% BSA or 0.1% Tween-20 + 3.0% BSA + 300 mM KSCN. *n* = 21 –30 FOVs per condition examined. **d,** Effect of fixation on mEGFP fluorescence intensity. mEGFP fluorescence intensities in live versus fixed cells were quantified, showing no substantial change before and after fixation. *n* = 72 FOVs for live cells, *n* = 77 FOVs for fixed cells. **e,** Effect of KSCN on mEGFP fluorescence intensity in fixed cells. mEGFP fluorescence intensities in fixed cells were quantified after incubation and imaging in buffers containing either 0.1% Tween-20 + 3.0% BSA or 0.1% Tween-20 + 3.0% BSA + 300 mM KSCN. The fluorescence intensity of mEGFP decreased by half in the presence of KSCN, possibly due to the chaotropic nature of thiocyanate, which may destabilize the GFP chromophore and lead to a fluorescence quenching. *n* = 21 FOVs for the mEGFP fluorescence intensity without the presence of KSCN, *n* = 23 FOVs for the mEGFP fluorescence intensity with the addition of KSCN. Each data point represents the average fluorescence intensity measured in a single FOV. Boxplots show the mean and boundaries (first and third quartiles); whiskers denote s.d. *p*-values calculated with two-sided unpaired Student’s *t*-test.

**Supplementary Fig. 15.**
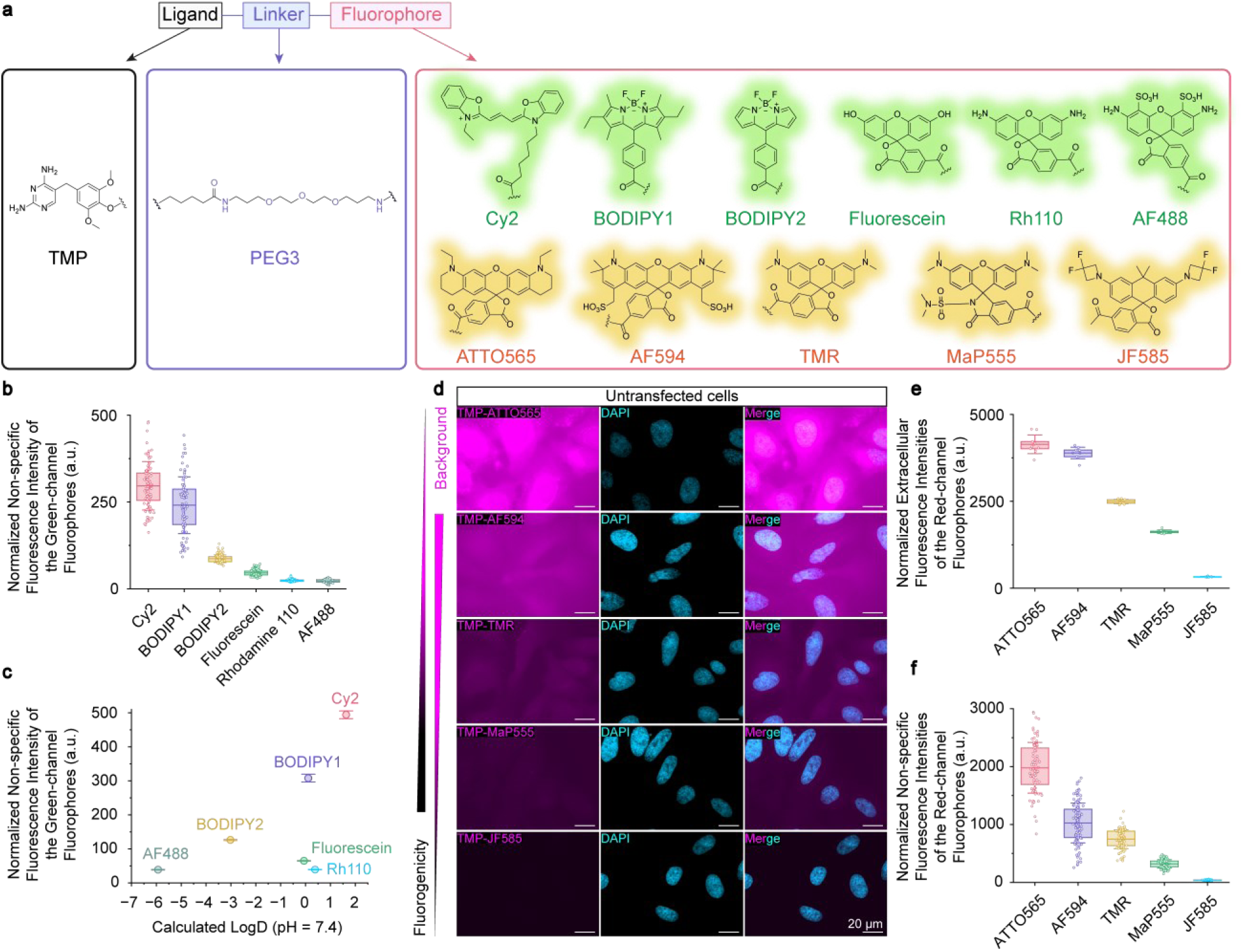
Investigation of the relationship between non-specific binding and the hydrophobicity or fluorogenicity of fluorophores. **a,** Chemical structures of TMP-conjugated fluorophores. The green-channel fluorophores (Cy2, BODIPY1^6^, BODIPY2^7^, Fluorescein, Rh110, and AF488), ranked by decreasing hydrophobicity (as indicated by calculated LogD), were used to evaluate the effect of hydrophobicity on non-specific binding. The red-channel fluorophores (ATTO565, AF594, TMR, MaP555, and JF585), ranked by increasing fluorogenicity, were used to evaluate the effect of fluorophore fluorogenicity on non-specific binding fluorescence (defined as the fluorescence measured within cell contours due to non-specific binding of TMP-conjugated fluorophores to protein surfaces in fixed cells) and extracellular fluorescence (defined as the fluorescence measured outside the cell contours caused by freely diffusing TMP-conjugated fluorophores in the imaging buffer). The fluorogenicity of rhodamine-based fluorophores arises from their ability to reversibly transition between a non-fluorescent, closed lactone form and a fluorescent, open zwitterionic form in solution. For fluorogenic imaging applications, an ideal rhodamine fluorophore remains “closed” in its unbound state, resulting in minimal background fluorescence. Upon binding to a target protein or entering a specific cellular environment, the equilibrium shifts toward the zwitterionic form, leading to a substantial fluorescence turn-on and a high signal-to-noise ratio. **b,** Quantifications of non-specific binding fluorescence intensity of the green-channel fluorophores. Non-specific binding fluorescence intensities were normalized by the brightness of each fluorophore. Each data point represents the average fluorescence intensity inside a single cell. *n* = 88–201 cells per condition examined. **c,** Correlaton between non-specific binding fluorescence and LogD. Non-specific binding of TMP-conjugated fluorophores showed a general positive correlation with calculated LogD (pH = 7.4). Notably, rhodamine-based fluorophores exhibited consistently lower levels of non-specific binding fluorescence compared to cyanine and BODIPY dyes. **d,** Representative epi-fluorescence images of fixed cells showing the non-specific binding fluorescence and extracellular fluorescence of the red-channel fluorophores. These red-channel rhodamine fluorophores, are in a known fluorogenic dye family, which can undergo a conformational shift from a non-fluorescent lactone (closed) form to a fluorescent zwitterionic (open) form upon target binding. TMP was conjugated to five red-channel rhodamine fluorophores with various fluorogenicity ranging from non-fluorogenic ATTO565 and AF594 to the highly fluorogenic JF585. Scale bars: 20 μm. **e,** Quantifications of extracellular fluorescence intensities of the red-channel fluorophores, normalized by each fluorophore’s brightness, revealing a negative correlation with fluorogenicity. Each data point represents the average background fluorescence intensity of a single FOV. *n* = 10 FOVs per condition examined. **f,** Quantifications of non-specific fluorescence intensities of the red-channel fluorophores, normalized by each fluorophores’ brightness, revealing a negative correlation with fluorogenicity.. Each data point represents the average fluorescence intensity inside a single cell. *n* = 82–124 cells per condition examined. Data are presented as mean ± s.d. Boxplots show the mean and boundaries (first and third quartiles); whiskers denote s.d.

**Supplementary Fig. 16.**
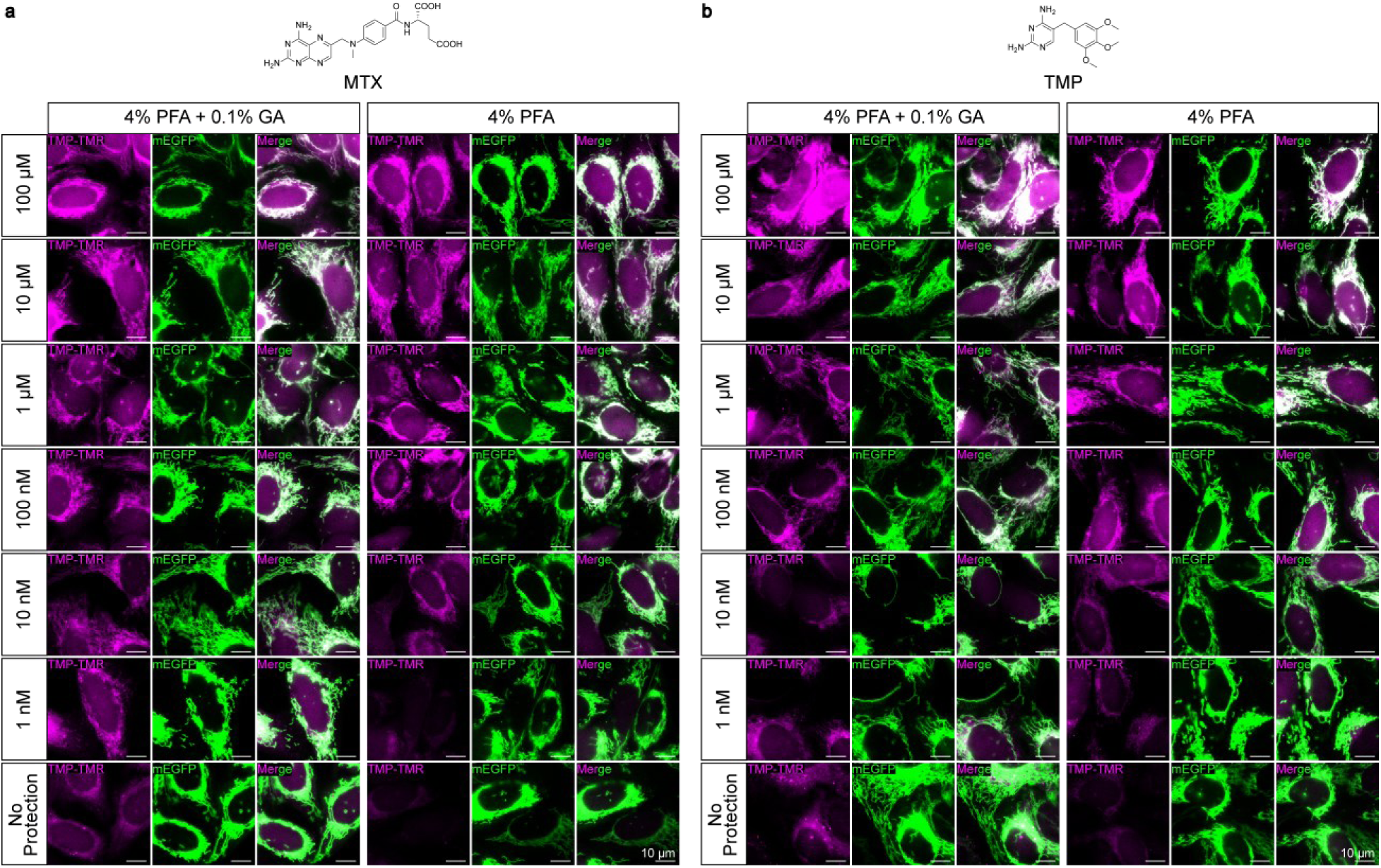
Protective fixation with MTX or TMP. **a,** Representative epi-fluorescence images of cells stably expressing TOM20-mEGFP-C(G)_5_-eDHFR^P89C^ (FLEXTAG2), labeled with 20 nM TMP-TMR under no-wash conditions following protective fixation. Cells were incubated with a gradient of MTX concentrations for 12 hours, followed by fixation with either PFA + GA (left) or PFA alone (right), each containing the same concentration of MTX as in the incubation step. MTX concentrations are indicated along the left side of the image panels. Scale bars: 10 μm. **b,** Representative epi-fluorescence images of cells stably expressing TOM20-mEGFP-C(G)_5_-eDHFR^P89C^ (FLEXTAG2), labeled with 20 nM TMP-TMR under no-wash conditions following protective fixation. Cells were incubated with a gradient of TMP concentrations for 1 hour, followed by fixation with either PFA + GA (left) or PFA alone (right), each containing the same concentration of TMP as in the incubation step. TMP concentrations are indicated along the left side of the image panels. Scale bars: 10 μm.

**Supplementary Fig. 17.**
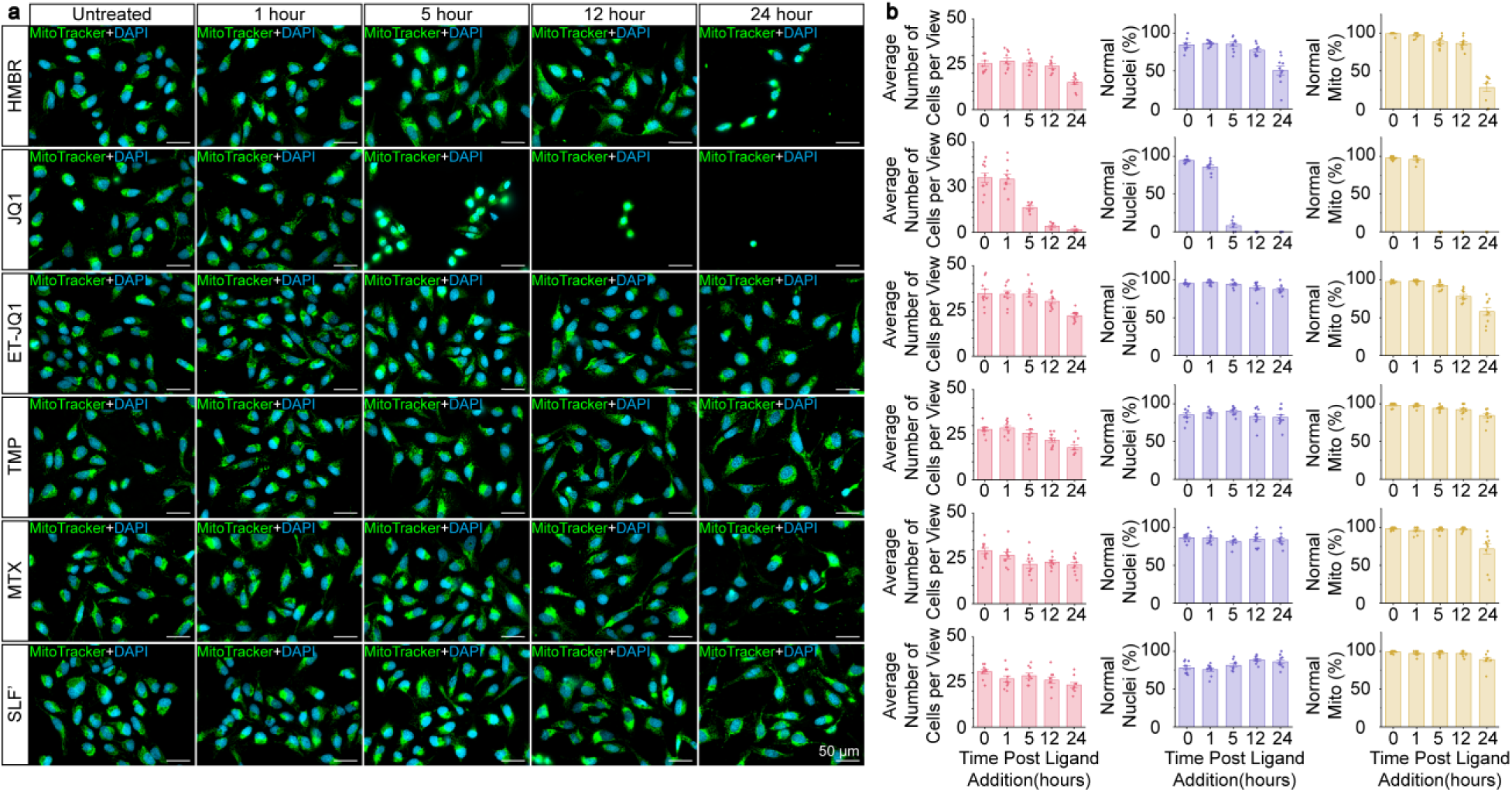
Evaluation of the cytotoxicity of protective agents via cell viability assay. **a,** Representative epi-fluorescence images of cells at 0, 1, 5, 12, and 24 hours after the addition of 100 µM HMBR, JQ1, ET-JQ1, TMP, MTX, or SLF’. Nuclei and mitochondria were labeled with 0.1 µg/mL DAPI and 50 nM MitoTracker Green, respectively. All time points were imaged from the same sample, with DAPI and MitoTracker Green added 30 minutes prior to the addition of the protective agents and maintained without restaining throughout the 24-hour period. Scale bars: 50 μm. **b,** Quantification of the cell number per FOV, and the fraction of cells with normal nucleus and mitochondria at 0, 1, 5, 12, and 24 hours after the addition of 100 µM HMBR, JQ1, ET-JQ1, TMP, MTX, or SLF’. Definitions of abnormal mitochondria and nuclear morphology are provided in the Method section. Each data point represents measurements from a single FOV. Data are presented as mean ± s.e.m. *n* = 10–11 FOVs per condition examined. Boxplots show the mean and boundaries (first and third quartiles); whiskers denote s.d.

**Supplementary Fig. 18.**
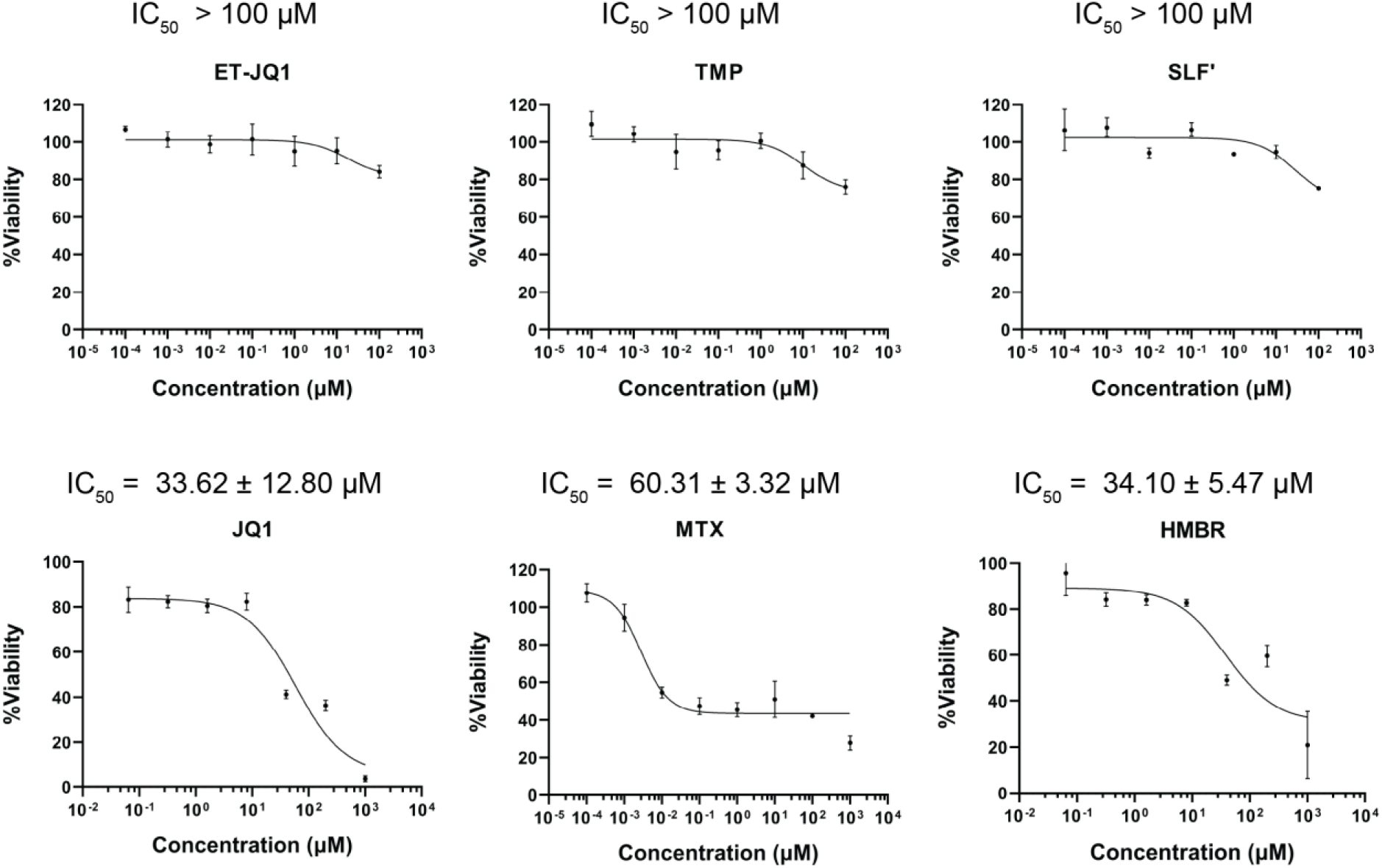
Evaluation of the cytotoxicity of protective agents via the MTT assay. U2OS cells were treated with increasing concentrations of ET-JQ1, TMP, SLF’, JQ1, MTX, or HMBR for 72 h at 37 °C, and cell viability was quantified using the MTT assay. Viability values were plotted as a function of compound concentration. ET-JQ1, TMP, and SLF’ exhibited minimal cytotoxicity with IC_50_ values >100 µM. JQ1, MTX, and HMBR displayed notable cytotoxicity with IC_50_ values of 33.62 ± 12.80 µM, 60.31 ± 3.32 µM, and 34.10 ± 5.47 µM, respectively. Data are shown as mean ± s.d. from three independent experiments. IC_50_ values were determined by nonlinear regression analysis of dose-response curves.

**Supplementary Fig. 19.**
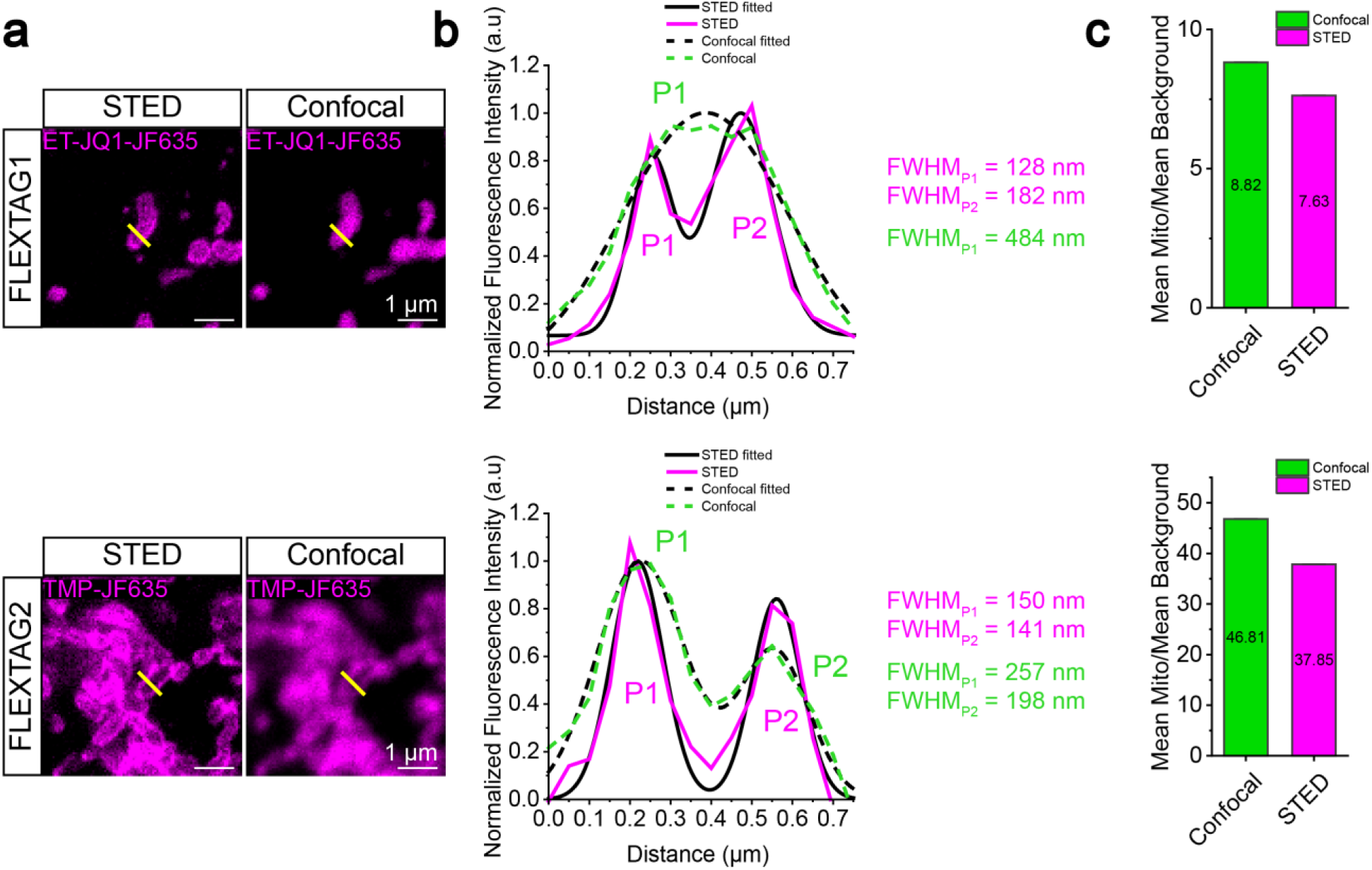
Comparison between STED and conventional confocal fluorescence microscopy. **a,** Representative STED and confocal fluorescence images of TOM20-mEGFP-FLEXTAG1 (top) and TOM20-mEGFP-FLEXTAG2 (bottom) labeled with 200 nM ET-JQ1-JF635 and TMP-JF635, respectively. The yellow lines indicate mitochondrial outer-membrane regions used for line-profile analysis. **b,** Fluorescence intensity profiles of ET-JQ1-JF635 (top) and TMP-JF635 (bottom) along the indicated lines for STED (magenta) and confocal (green) imaging. Solid lines represent raw intensity traces; dashed lines represent corresponding Gaussian fits. STED profiles exhibit reduced full width at half maximum (FWHM) relative to confocal. **c,** Quantified signal-to-noise ratios for confocal and STED images in **a**. Scale bars: 1 μm.

**Supplementary Fig. 20.**
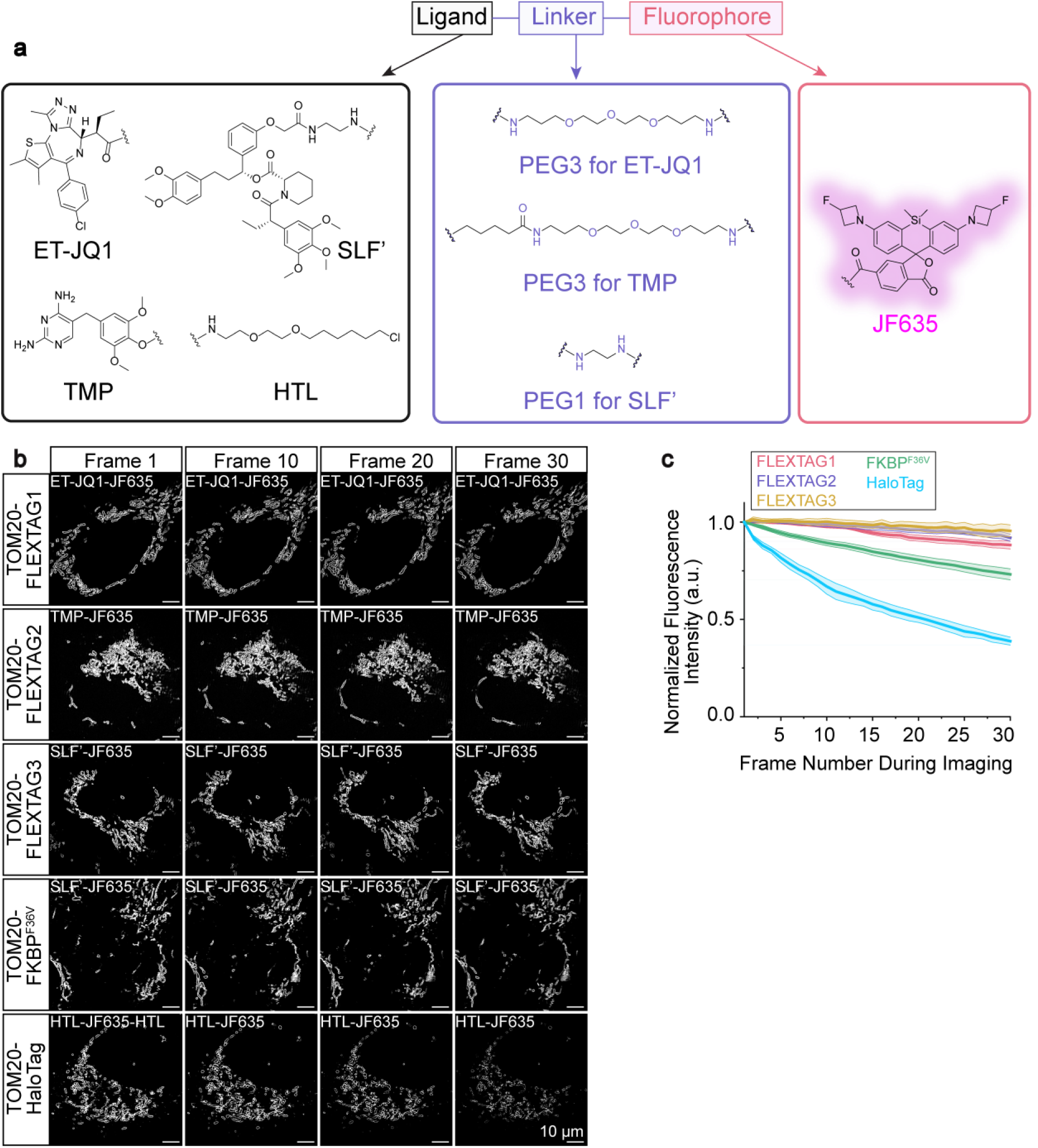
Evaluation of the anti-fade performance of FLEXTAG1, FLEXTAG2, and FLEXTAG3 using JF635-conjugated ligands. **a,** Chemical structures of JF635 conjugated to ET-JQ1, TMP, SLF’, and HaloTag Ligand (HTL)-. The HTL molecule contains both warhead targeting the HaloTag binding pocket and a PEG-based linker, allowing direct conjugation to JF635 without the need for additional linkers. **b,** Representative time-lapse SIM images of a single live cell overexpressing TOM20-mEGFP-FLEXTAG1/FLEXTAG2/FLEXTAG3/ FKBP^F36V^/HaloTag, labeled with 500 nM JF635-conjugated ET-JQ1, TMP, SLF’, or HTL. Images were shown at frames 1, 10, 20, and 30. Scale bars: 10 μm. **c,** Quantification of fluorescence intensity of JF635-labeled FLEXTAG1/FLEXTAG2/FLEXTAG3/ FKBP^F36V^/HaloTag over 30-frame live-cell SIM imaging. Data are presented as mean ± s.e.m. *n* = 3 biological replicates; 3 FOVs were examined for each protein tag.

**Supplementary Fig. 21.**
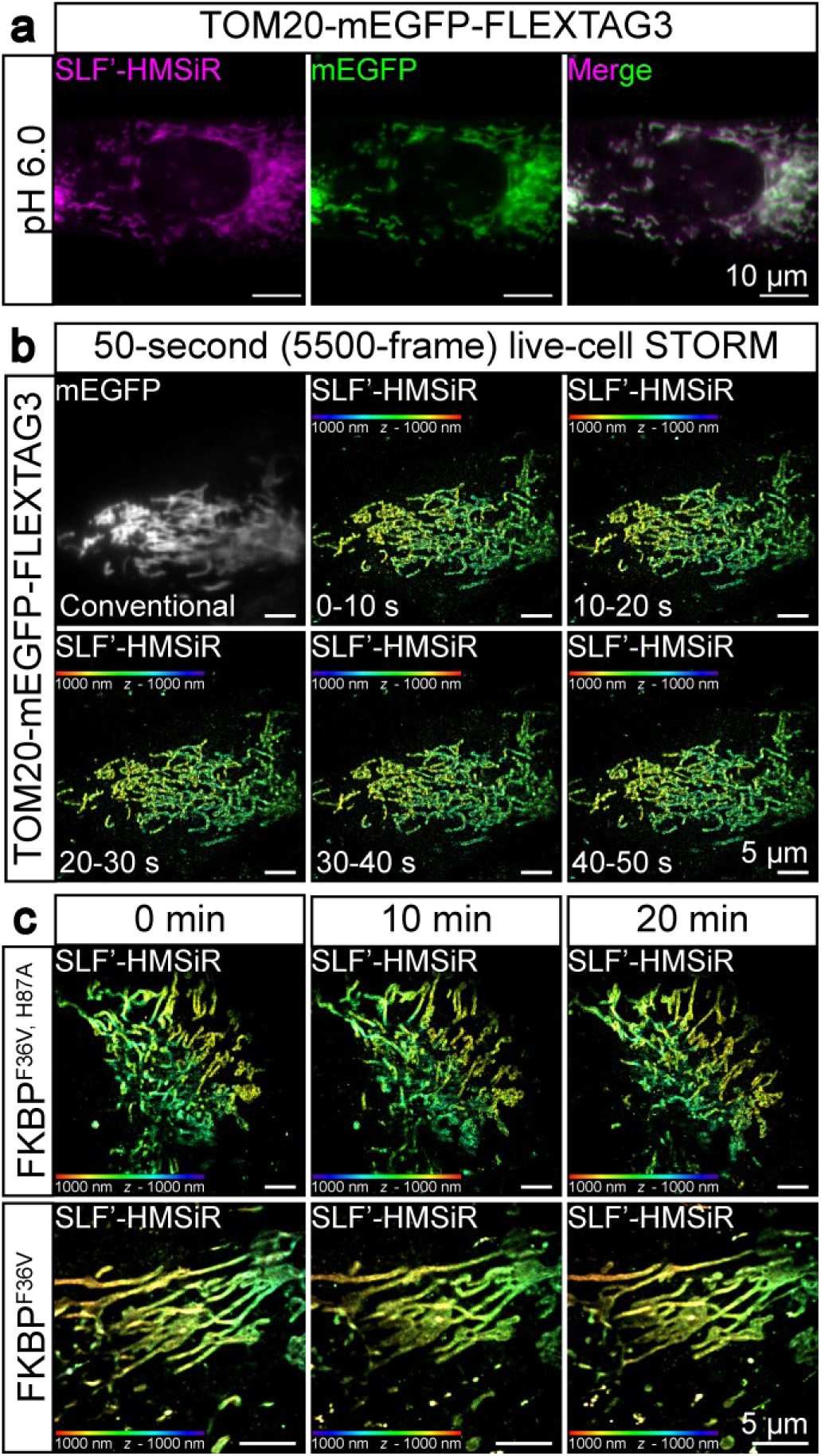
Conventional and super-resolution imaging with FLEXTAG3 and SLF’-HMSiR. **a,** Epi-fluorescnece images of fixed cells overexpressing TOM20-mEGFP-FLEXTAG3, labeled with 100 nM SLF’-HMSiR. The pH of the imaging buffer (DPBS) was adjusted to 6.0 to enhance the fluorescence intensity of HMSiR, as the equilibrium between its non-fluorescent “closed” form and fluorescent “open” form is highly pH dependent. Under acidic conditions, protonation of the carboxylic acid group shifts the equilibrium toward the open, zwitterionic state, thereby turning on fluorescence and improving signal intensity. Scale bars: 10 μm. **b,** Conventional and 3D-STORM images of live cells overexpressing TOM20-mEGFP-FLEXTAG3, labeled with 30 nM SLF’-HMSiR, in the same FOV over 0–50 seconds (5,500 frames at 110 Hz). Notably, SLF’-HMSiR may tend to aggregate under certain conditions. To minimize this effect, the probe solution was passed through a 0.22 µm filter and sonicated prior to imaging to remove aggregated fluorophores. Scale bars: 5 μm. **c,** Representative 3D-STORM images of live cells overexpressing TOM20-mEGFP-FKBP^F36V, H87A^ and TOM20-mEGFP-FKBP^F36V^, labeled with 30 nM SLF’-HMSiR, from three imaging cycles. Each cycle consisted of 4.5 minutes of data acquisition (30,000 frames, 110 Hz) followed by 5.5 minutes of dark recovery. Scale bars: 5 μm.

### Supplementary Tables

**Supplementary Table 1.** Calculated single-molecule fitting precision and NeNA precision of FLEXTAG-based PAINT.

| Protein Tag | Fluorescent Ligand | Color Channel | Excitation Wavelength (nm) | Single-molecule Fitting Precision (nm) | NeNA (nm) |
| --- | --- | --- | --- | --- | --- |
| FLEXTAG1 | ET-JQ1-AF488 | Green | 488 | 3.53 | 27.5 |
|  | ET-JQ1-TMR | Red | 560 | 3.39 | 21.5 |
|  | ET-JQ1-AF647 | Far-red | 642 | 5.03 | 30.9 |
| FLEXTAG 2 | TMP-AF488 | Green | 488 | 2.61 | 21.3 |
|  | TMP-TMR | Red | 560 | 2.97 | 33.2 |
|  | TMP-JF635 | Far-red | 642 | 3.49 | 21.0 |
| FLEXTAG 3 | SLF'-AF488 | Green | 488 | 4.03 | 34.8 |
|  | SLF'-MaP555 | Red | 560 | 3.91 | 34.6 |
|  | SLF'-HMSiR | Far-red | 642 | 3.05 | 24.7 |

**Supplementary Table 2.**
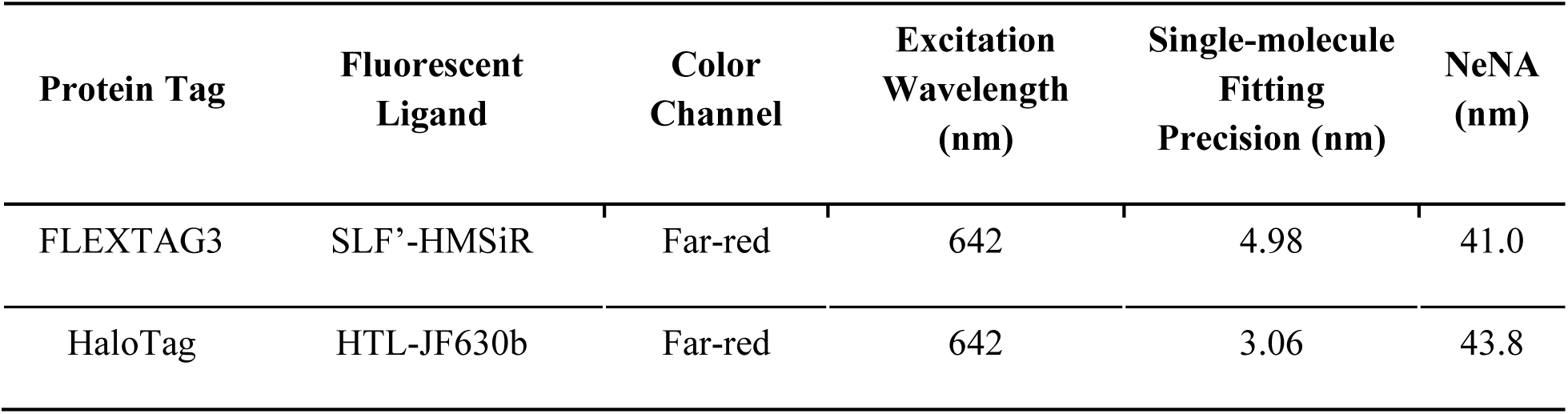
Calculated fitting precision and NeNA precision of FLEXTAG-based and HaloTag-based live-cell STORM.

| Protein Tag | Fluorescent Ligand | Color Channel | Excitation Wavelength (nm) | Single-molecule Fitting Precision (nm) | NeNA (nm) |
| --- | --- | --- | --- | --- | --- |
| FLEXTAG3 | SLF'-HMSiR | Far-red | 642 | 4.98 | 41.0 |
| HaloTag | HTL-JF630b | Far-red | 642 | 3.06 | 43.8 |

**Supplementary Table 3.** Percentages of cells exhibiting normal mitochondrial morphology when overexpressing TOM20-mEGFP-FLEXTAGs.

| Protein Tag | FLEXTAG1 |  | FLEXTAG2 |  | FLEXTAG3 |  | mEGFP (baseline) |
| --- | --- | --- | --- | --- | --- | --- | --- |
| Ligand | – ET-JQ1 | + ET-JQ1 | – TMP | + TMP | – SLF’ | + SLF’ | N.A. |
| % Normal Mito (mean ± s.e.m) | 88.6 ± 2.5 | 80.8 ± 2.5 | 92.1 ± 1.3 | 92.4 ± 1.2 | 75.8 ± 3.3 | 75.9 ± 2.0 | 89.8 ± 1.6 |

### Supplementary Methods

NMR spectra and HRMS data of the below synthesized compounds are provided in Supplementary Data 2 and Supplementary Data 3, respectively.

#### Synthesis of HMSiR

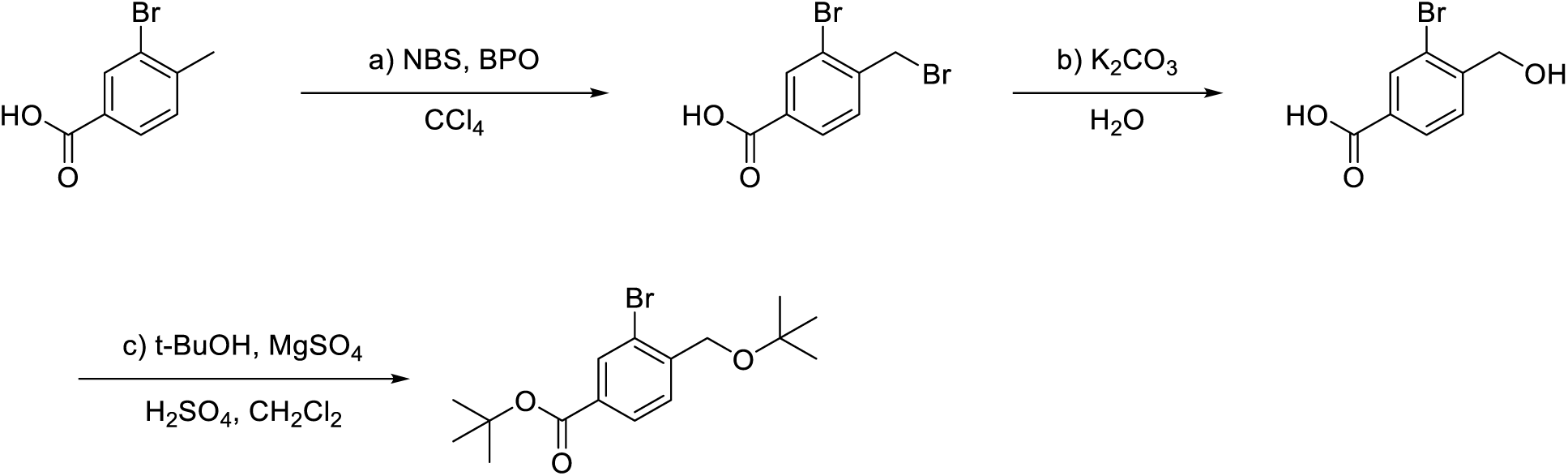

a) 2.15 g (10 mmol, 1 eq.) 3-Bromo-4-methylbenzoic acid was suspended in 10 mL carbon tetrachloride. 1.96 g (11 mmol, 1.1 eq) N-bromosuccinimide and 120 mg (0.5 mmol, 0.05 eq.) benzoyl peroxide were added to the reaction mixture. The reaction was stirred at 100 ℃ in an nitrogen or argon atmosphere for 2 hours. The reaction mixture was chilled to 0 ℃, filtered, the filtrate was collected and recrystallized from diethyl ether. Product was obtained as a white solid, yield 50%.

b) 586 mg (2 mmol, 1 eq.) 3-Bromo-4-(bromomethyl)benzoic acid and 1.1 g (8 mmol, 4 eq.) potassium carbonate was suspended in 20 mL water. The reaction mixture was stirred at 100 ℃ for 5 hours. The solution was added 1M hydrochloric acid and the precipitated white solid was collected by filtration. Product was obtained as a white solid, yield 90%.

c) 960 mg (8 mmol, 8 eq.) Magnesium sulfate was suspended in 20 mL dichloromethane and 196 mg (2 mmol, 2 eq.) sulfuric acid was added to the mixture while stirring. 230 mg (1 mmol, 1 eq.) 3-Bromo-4-(hydroxymethyl)benzoic acid and 740 mg (10 mmol, 10 eq.) tert-butanol were subsequently added, and the reaction was stirred at RT overnight. The reaction was quenched with saturated sodium bicarbonate solution, extracted with ethyl acetate, washed with brine, and dried by sodium sulfate. Solvent was evaporated and the residue was purified by chromatography column (5% ethyl acetate in hexane). The product was obtained as a colorless oil, yield 45%^10^.

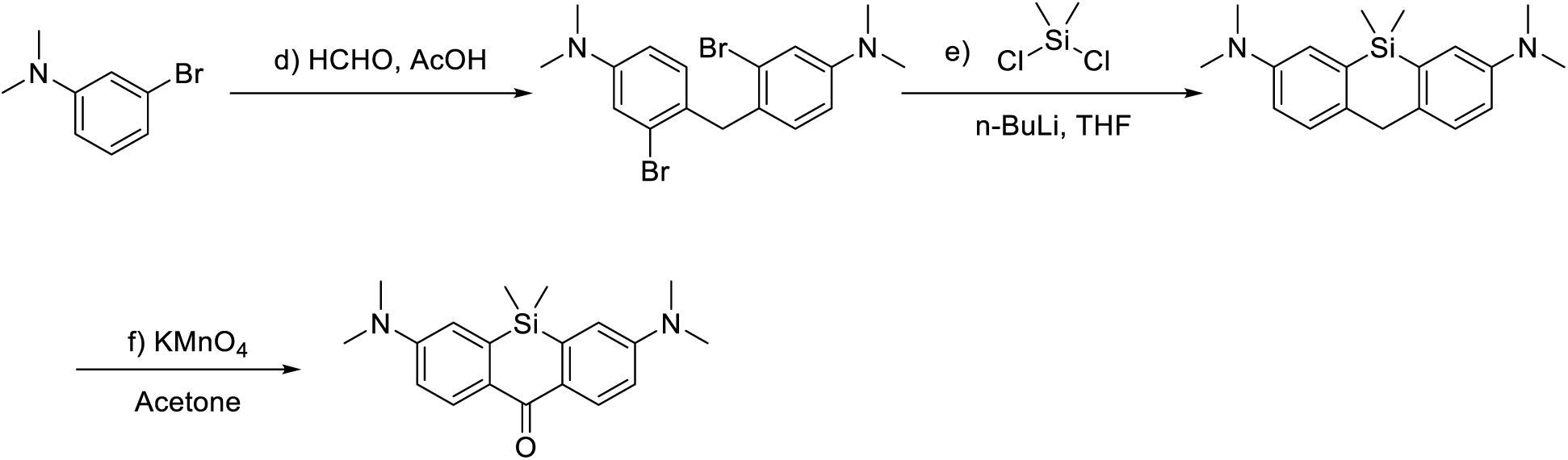

d) 1 g (5 mmol, 1 eq.) 3-Bromo-N, N-dimethylaniline was suspended in 10 mL acetic acid. 1.5 g (50 mmol, 10 eq.) paraformaldehyde was added, and the reaction mixture was stirred at 60 ℃ for 2 hours. Excess acetic acid was removed in vacuo and the remaining acetic acid was neutralized by saturated sodium bicarbonate solution. The mixture was extracted with ethyl acetate, washed with brine, and dried by sodium sulfate. Solvent was evaporated and the residue was purified by chromatography column (5% ethyl acetate in hexane). The product was obtained as a white solid, yield 69%.

e) 400 mg (1 mmol, 1 eq.) product from last step was suspended in anhydrous tetrahydrofuran, chilled to −78 ℃, and 1mL (2.5 mmol, 2.5 eq.) n-butyl lithium (2.5 M in hexane) was added drop wisely while stirring in an nitrogen or argon atmosphere. The solution was maintained at −78 ℃, and stirred for 30 min. 120 mg (0.5 mmol, 0.5 eq.) dichlorodimethylsilane was added drop wisely while stirring, and the reaction gradually warmed up to RT, and stirred for 2 hours. The reaction was quenched with saturated ammonium chloride solution, extracted with ethyl acetate, washed with brine, and dried by sodium sulfate. The solvent was evaporated, and the resulting residue was used immediately without purification.

f) 5mL acetone was added to the residue, followed by adding 470 mg (3 mmol, 3 eq.) potassium permanganate in portion. The reaction was stirred for 2 hours, and the solvent was removed in vacuo. The residue was purified by chromatography column (n-hexane/dichloromethane/ethyl acetate, 5:1:1, (vol/vol)). The product was obtained as a light-yellow solid with green fluorescence, yield 17%^11^.

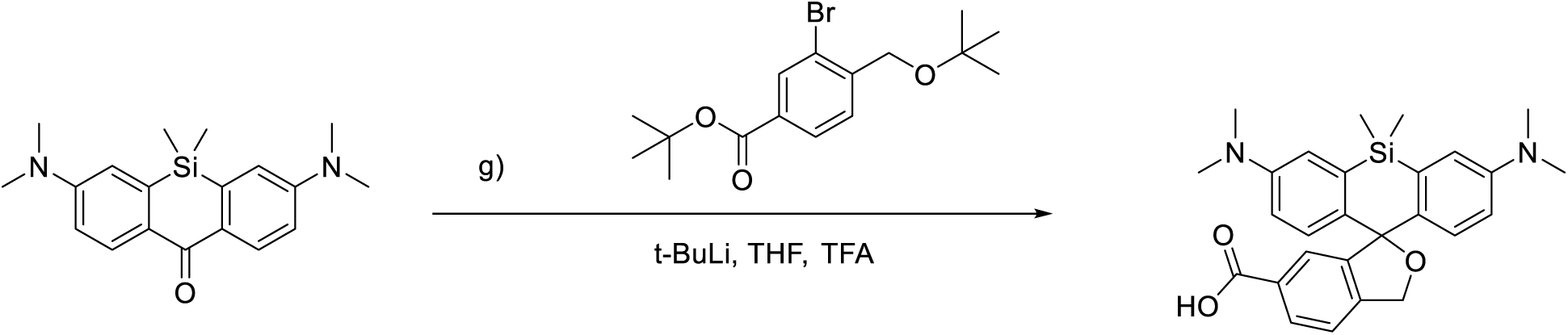

g) 171 mg (0.5 mmol, 3 eq.) benzoic acid, 3-bromo-4-[(1,1-dimethylethoxy)methyl]-, 1,1-dimethylethyl ester was suspended in 5 mL anhydrous tetrahydrofuran, chilled to −78 ℃, and 300 µL (0.5 mmol, 3 eq.) tert-butyl lithium (1.7 M in pentane) was added drop wisely while stirring in an nitrogen or argon atmosphere. The solution was maintained at −78 ℃ and stirred for 30 min. 50 mg (0.16 mmol, 1 eq.) 2,7-Bis(dimethylamino)-9,9-dimethyl-9-silaanthracen-10(9H)-one was dissolved in 1 mL anhydrous tetrahydrofuran, and was added drop wisely while stirring. The reaction was gradually warmed up to RT, and stirred overnight. The reaction was quenched with dropwise addition of acetic acid, and the solvent was evaporated in vacuo. The resulting residue was suspended in trifluoroacetic acid and was stirred at RT overnight. The reaction mixture was neutralized by saturated sodium bicarbonate solution, extracted with dichloromethane, dried by sodium sulfate, and purified by chromatography column (10% methanol in dichloromethane). Product obtained as a light blue solid, yield 35%.

^1^H NMR (500 MHz, Chloroform-*d*) δ 8.02 (dd, J = 7.9, 1.4 Hz, 2H), 7.78 (s, 1H), 7.38 (d, J = 7.9 Hz, 2H), 7.02 – 6.99 (m, 4H), 6.65 (dd, J = 8.9, 2.9 Hz, 2H), 5.34 (s, 2H), 2.95 (s, 12H), 0.67 (s, 3H), 0.55 (s, 3H). ^13^C NMR (126 MHz, Chloroform-*d*) δ 170.74, 148.67, 145.19, 138.38, 135.24, 130.20, 129.39, 129.00, 128.21, 126.34, 121.29, 117.13, 114.20, 50.85, 40.66, 0.49, −0.94. HRMS [M+H]^+^: Calcd 459.2104, found 459.2096.

#### Synthesis of HBR-3,5DOM

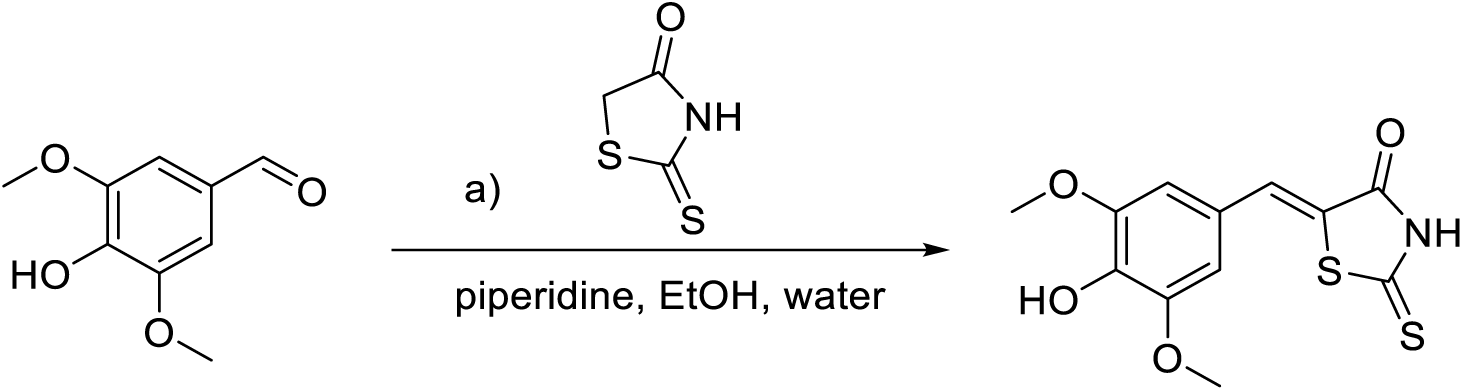

200 mg (1.5 mmol, 1 eq.) rhodanine and 300 mg (1.65 mmol, 1.1 eq.) 4-hydroxy-3,5-dimethoxybenzaldehyde were suspended in 100 mL water. 4.5 mL ethanol and 128 mg (1.5 mmol, 1 eq.) piperidine were added, and the reaction mixture was stirred at RT overnight. The reaction mixture was then neutralized by addition of 1 M hydrochloric acid solution. After standing in cold room (2 ℃) overnight, the precipitate was filtered and collected. The product ((Z)-5-(4-hydroxy-3,5-dimethoxybenzylidene)-2-thioxothiazolidin-4-one) was obtained as a yellow crystal, yield 66%^12^.

^1^H NMR (500 MHz, DMSO-*d_6_*) δ 9.48 (s, 1H), 7.58 (s, 1H), 6.89 (s, 2H), 3.84 (s, 6H).

^13^C NMR (126 MHz, DMSO-*d_6_*) δ 195.88, 169.91, 148.78, 139.62, 133.46, 123.70, 121.95, 108.94.

#### Synthesis of HMBR

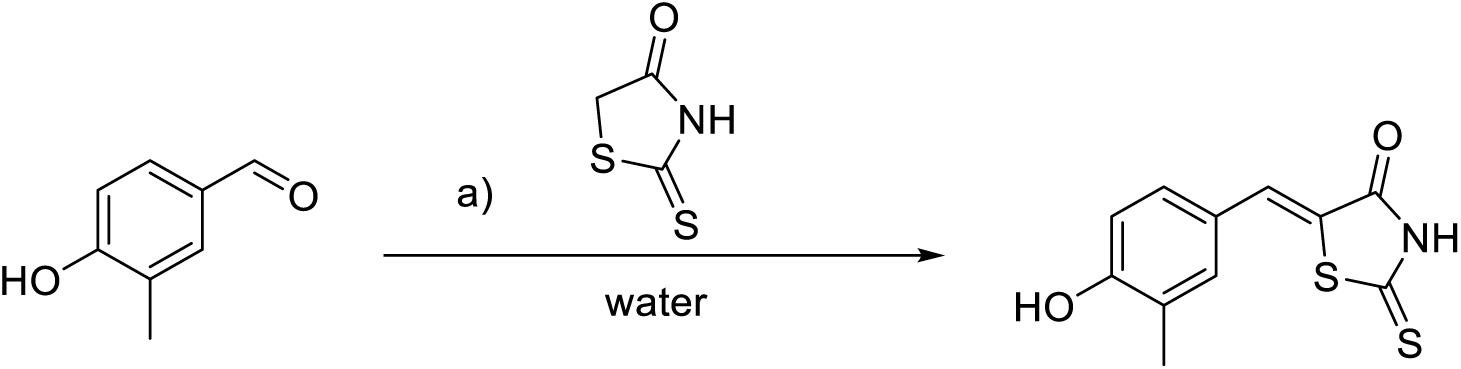

200 mg (1.5 mmol, 1 eq.) rhodanine and 225 mg (1.65 mmol, 1.1 eq.) 4-hydroxy-3-methylbenzaldehyde were suspended in 100 mL water. The reaction was stirred at 80℃ for 7 days. The reaction was allowed to stand in cold room (2 ℃) overnight, and the precipitate was filtered and collected. The product ((Z)-5-(4-hydroxy-3-methylbenzylidene)-2-thioxothiazolidin-4-one) was obtained as a yellow crystal, yield 55%^1^.

^1^H NMR (500 MHz, DMSO-*d_6_*) δ 10.38 (s, 1H), 7.52 (s, 1H), 7.34 (s, 1H), 7.31 (d, J = 8.4 Hz, 1H), 6.94 (d, J = 8.4 Hz, 1H), 2.54 (s, 3H).

^13^C NMR (126 MHz, DMSO-*d_6_*) δ 196.04, 169.98, 159.19, 134.13, 133.10, 131.10, 125.96, 124.31, 121.10, 116.08, 16.36.

#### Synthesis of BODIPYs

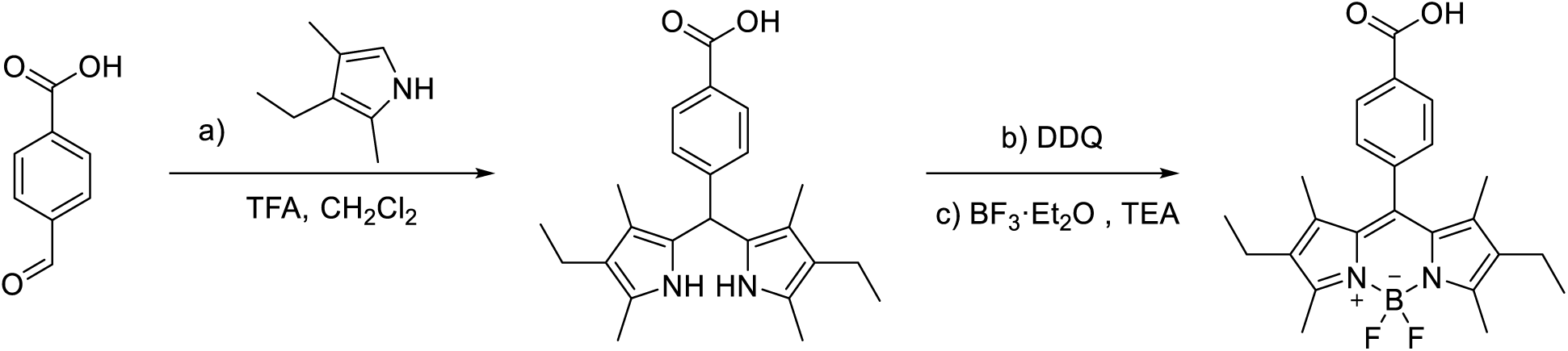

a) 500 mg (4 mmol, 2 eq.) 3-ethyl-2,4-dimethylpyrrole, 300 mg (2 mmol, 1 eq.) 4-carboxybenzaldehyde and catalytic amount of trifluoroacetic acid were suspended in 100 mL dichloromethane. The reaction mixture was stirred at RT overnight under nitrogen/argon atmosphere.

b) 450 mg (2 mmol, 1 eq.) 2,3-dichloro-5,6-dicyano-*p*-benzoquinone in 20 mL dichloromethane was added in portion. The reaction mixture was stirred at RT for 1 hour.

c) 4 mL (28 mmol, 14 eq.) triethylamine was added, followed by addition of 4 mL BF_3_·Et_2_O (32 mmol, 16 eq.). The reaction mixture was stirred at RT for 2 hours. The reaction mixture was then washed by water and brine, dried by sodium sulfate, and purified by chromatography column (hexane: ethyl acetate = 1: 1 (vol/vol)). Product was obtained as a dark red solid, 3-step yield 45%^6^.

^1^H NMR (500 MHz, Chloroform-*d*) δ 8.34 – 8.24 (m, 2H), 7.53 – 7.44 (m, 2H), 2.57 (s, 6H), 2.38 – 2.28 (m, 4H), 1.30 (s, 6H), 1.01 (s, 6H).

^13^C NMR (126 MHz, Chloroform-*d*) δ 171.49, 154.40, 141.74, 138.43, 133.16, 130.89, 130.22, 129.75,

128.91, 119.28, 113.93, 17.08, 14.61, 12.58, 11.87.

HRMS [M-H]^-^: Calcd 423.2061, found 423.2060.

#### Synthesis of TMP-linker-boc

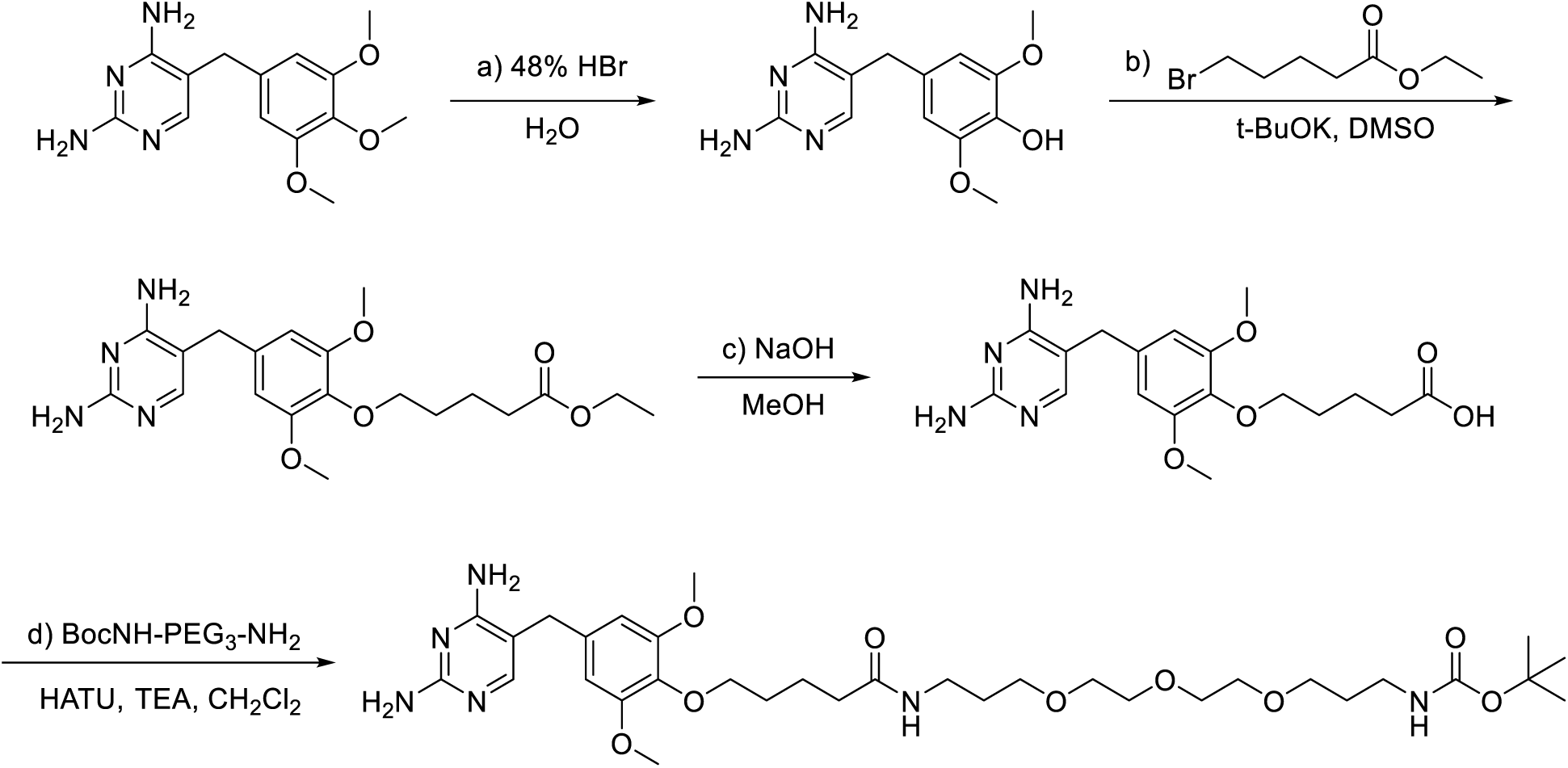

a) 5 g (17.2 mmol, 1 eq.) trimethylprim was suspended in 60 mL (excess) 48% hydrobromic acid. The reaction was heated to 100 ℃ and stirred 20–30 min (no exceed 30 min). The reaction was cooled to RT, and 50% sodium hydroxide solution was added in portion. The reaction mixture was allowed to stand in cold room (2 ℃) overnight to precipitate. The precipitated solid was collected by filtration, washed by 100 mL ice-cold water, and re-dissolved in 200 mL boiling water. pH was adjusted to 7 with 28% ammonia solution and the solution was allowed to stand in cold room overnight to recrystallize. The precipitated solid was collected by filtration. Product (4-[(2,4-Diamino-5-pyrimidinyl)methyl]-2,6-dimethoxyphenol) was obtained as a light pink solid, yield 60%.

b) 2.3 g (8.3 mmol, 1 eq) 4-[(2,4-Diamino-5-pyrimidinyl)methyl]-2,6-dimethoxyphenol was suspended in 30 mL DMSO, and 2.8 g (25 mmol, 3 eq.) Potassium tert-butoxide was added under nitrogen/argon atmosphere. The reaction mixture was stirred at RT for 30 min. 1.9 g (9.13 mmol, 1.1 eq.) Ethyl 5-bromovalerate was added in portion, and the reaction mixture was stirred overnight. Solvent was removed under high-vac, and the residue was purified by chromatography column (10% methanol in dichloromethane). The product (Ethyl 5-[4-[(2,4-diamino-5-pyrimidinyl)methyl]-2,6-dimethoxyphenoxy]pentanoate) was obtained as a white solid, yield 47%^13^.

^1^H NMR (400 MHz, Chloroform-*d*) δ 7.75 (s, 1H), 6.37 (s, 2H), 5.16 – 4.99 (m, 2H), 4.80 (s, 2H), 4.12 (q, J = 7.0 Hz, 2H), 3.94 (t, J = 5.6 Hz, 2H), 3.78 (s, 6H), 3.64 (s, 2H), 2.38 (t, J = 7.0 Hz, 2H), 1.88 – 1.73 (m, 4H), 1.25 (t, J = 7.0 Hz, 3H).

HRMS [M+H]^+^: Calcd 405.2129, found 405.2132.

c) 404 mg (1mmol, 1 eq.) Ethyl 5-[4-[(2,4-diamino-5-pyrimidinyl)methyl]-2,6-dimethoxyphenoxy]pentanoate was suspended in 15 mL 1M sodium hydroxide in methanol. The reaction mixture was stirred at RT overnight. The reaction was neutralized by 1M hydrochloric acid solution, and the solvent was evaporated. The crude was used immediately in the next step without further purification.

d) The crude from the last step was suspended in 8 mL dichloromethane. 480 mg (1.5 mmol, 1.5 eq.) N-Boc-4,7,10-trioxa-1,13-tridecanediamine and 550 mg (5 mmol, 5 eq.) triethylamine was added. 456 mg (1.2 mmol, 1.2 eq.) HATU was suspended in 2 mL dichloromethane, and was added in portion. The reaction mixture was stirred at RT overnight. Solvent was removed in vacuo, and the resulting residue was purified by chromatography column (10% methanol in dichloromethane). The product (tert-butyl (19-(4-((2,4-diaminopyrimidin-5-yl)methyl)-2,6-dimethoxyphenoxy)-15-oxo-4,7,10-trioxa-14-azanonadecyl)carbamate) was obtained as a white solid, yield 29%.

^1^H NMR (500 MHz, Chloroform-*d*) δ 7.61 (s, 1H), 6.38 (s, 2H), 3.93 (t, J = 6.0 Hz, 2H), 3.77 (s, 6H), 3.63 – 3.60 (m, 8H), 3.59 – 3.57 (m, 4H), 3.52 (dd, J = 6.0, 2.6 Hz, 4H), 3.32 (q, J = 6.2 Hz, 2H), 3.21 – 3.17 (m, 2H), 2.25 (t, J = 7.2 Hz, 2H), 1.76 (dq, J = 17.7, 6.6 Hz, 9H), 1.42 (s, 9H).

^13^C NMR (126 MHz, Chloroform-*d*) δ 173.61, 163.03, 160.96, 153.64, 133.54, 106.79, 105.25, 79.03, 72.90, 70.40, 70.07, 69.31, 56.10, 38.27, 37.46, 36.15, 34.27, 29.62, 29.39, 29.04, 28.43, 22.42.

HRMS [M+H]^+^: Calcd 679.4025, found 679.4007.

#### Synthesis of SLF’-linker-boc

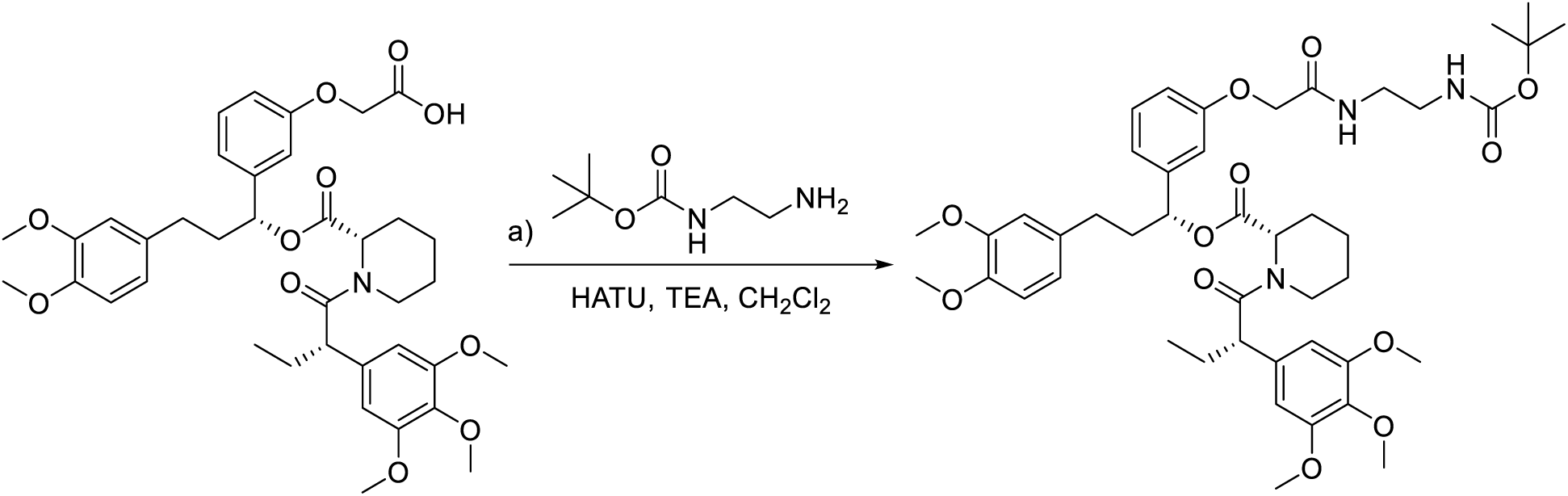

a) 13.8 mg (0.02 mmol, 1 eq.) 2-(3-((R)-3-(3,4-dimethoxyphenyl)-1-(((S)-1-((S)-2-(3,4,5-trimethoxyphenyl)butanoyl)piperidine-2-carbonyl)oxy)propyl)phenoxy)acetic acid was suspended in 2 mL dichloromethane. 4.8 mg (0.03 mmol, 1.5 eq.) tert-butyl (2-aminoethyl)carbamate and 55 mg (0.5 mmol, 25 eq.) triethylamine was added. 9.1 mg (0.024 mmol, 1.2 eq.) HATU was suspended in 1 mL dichloromethane, and was added in portion. The reaction mixture was stirred at RT overnight. Solvent was removed in vacuo, and the resulting residue was purified by preparative TLC plate (5% methanol in dichloromethane). The product ((R)-1-(3-(2-(2-((tert-butoxycarbonyl)amino)ethoxy)-2-oxoethoxy)phenyl)-3-(3,4-dimethoxyphenyl)propyl (S)-1-((S)-2-(3,4,5-trimethoxyphenyl)butanoyl)piperidine-2-carboxylate) was obtained as a white solid, yield 81%.

^1^H NMR (500 MHz, Chloroform-*d*) δ 7.35 – 7.30 (m, 1H), 7.19 (t, *J* = 7.9 Hz, 2H), 6.81 – 6.79 (m, 2H), 6.79 – 6.75 (m, 5H), 6.68 (d, *J* = 8.0 Hz, 3H), 6.66 (d, *J* = 9.1 Hz, 5H), 6.43 (s, 1H), 6.41 (s, 4H), 5.83 (t, *J* = 6.8 Hz, 1H), 5.63 (dd, *J* = 8.2, 5.4 Hz, 2H), 5.47 (d, *J* = 5.3 Hz, 2H), 4.47 (s, 3H), 3.79 (s, 6H), 3.68 (s, 9H), 3.48 (d, *J* = 5.7 Hz, 5H), 2.80 (td, *J* = 13.3, 3.0 Hz, 2H), 2.56 (td, *J* = 9.3, 4.6 Hz, 2H), 2.47 (ddd, *J* = 14.4, 9.3, 6.8 Hz, 2H), 2.36 – 2.27 (m, 2H), 2.15 – 2.05 (m, 5H), 1.93 (tt, *J* = 10.5, 5.3 Hz, 2H), 1.72 (hept, *J* = 6.8, 6.3 Hz, 9H), 1.42 (s, 10H), 0.90 (t, *J* = 7.2 Hz, 7H).

^13^C NMR (126 MHz, Chloroform-*d*) δ 172.66, 170.59, 168.87, 157.36, 156.56, 153.50, 153.17, 148.88, 147.35, 142.31, 136.63, 135.30, 133.35, 129.78, 120.20, 119.88, 112.91, 111.70, 111.28, 104.99, 104.60, 75.62, 67.30, 60.79, 56.31, 55.97, 55.84, 52.06, 50.86, 43.48, 38.21, 31.31, 28.36, 26.79, 25.33, 20.91, 12.56.

HRMS [M+Na]^+^: Calcd 858.4147, found 858.4130.

#### Synthesis of SLF’’-linker-boc

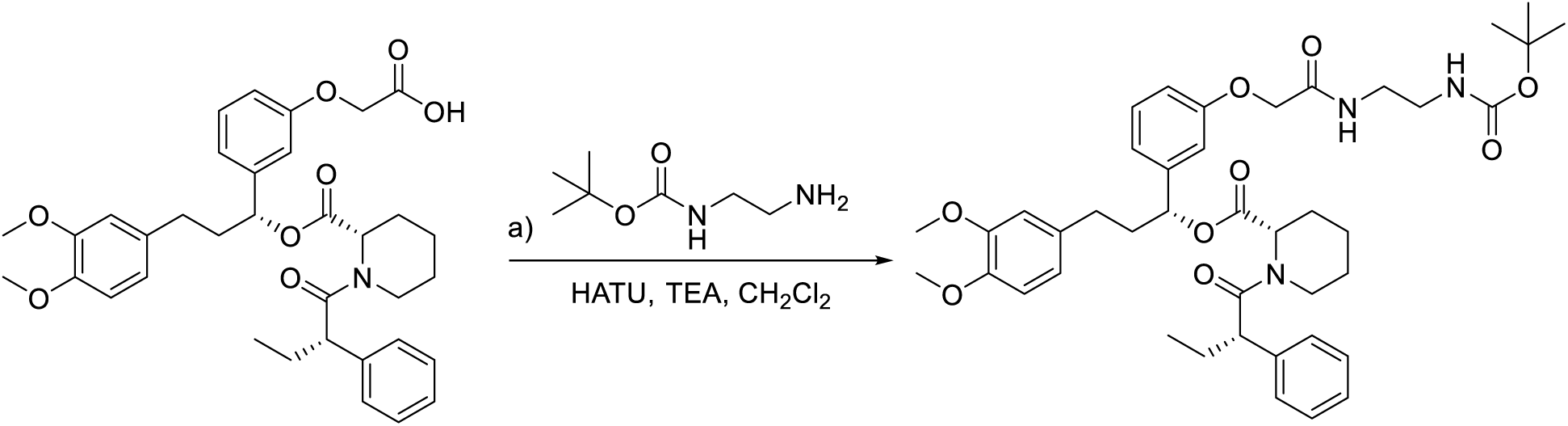

a) 12.1 mg (0.02 mmol, 1 eq.) 2-(3-((R)-3-(3,4-dimethoxyphenyl)-1-(((S)-1-((S)-2-phenylbutanoyl)piperidine-2-carbonyl)oxy)propyl)phenoxy)acetic acid was suspended in 2 mL dichloromethane. 4.8 mg (0.03 mmol, 1.5 eq.) tert-butyl (2-aminoethyl)carbamate and 55 mg (0.5 mmol, 25 eq.) triethylamine was added. 9.1 mg (0.024 mmol, 1.2 eq.) HATU was suspended in 1 mL dichloromethane, and was added in portion. The reaction mixture was stirred at RT overnight. Solvent was removed in vacuo, and the resulting residue was purified by preparative TLC plate (5% methanol in dichloromethane). The product ((R)-1-(3-(2-(2-((tert-butoxycarbonyl)amino)ethoxy)-2-oxoethoxy)phenyl)-3-(3,4-dimethoxyphenyl)propyl (S)-1-((S)-2-phenylbutanoyl)piperidine-2-carboxylate) was obtained as a white solid, yield 86%.

^1^H NMR (500 MHz, Chloroform-*d*) δ 7.33 – 7.13 (m, 6H), 7.00 – 6.52 (m, 6H), 5.86 – 5.62 (m, 1H), 5.53 – 5.34 (m, 1H), 5.07 – 4.55 (m, 1H), 4.44 (d, J = 53.4 Hz, 2H), 3.85 (s, 6H), 3.70 – 3.27 (m, 5H), 2.80 (s, 2H), 2.70 – 2.54 (m, 2H), 2.51 – 1.50 (m, 10H), 1.42 (s, 9H), 1.29 – 1.13 (m, 2H), 0.88 (t, J = 7.2 Hz, 3H).

^13^C NMR (126 MHz, Chloroform-*d*) δ 172.43, 170.69, 170.16, 168.76, 168.62, 157.45, 157.16, 148.93, 148.84, 147.47, 147.30, 142.23, 142.03, 140.16, 139.68, 133.53, 133.19, 130.02, 129.82, 128.96, 128.68, 128.11, 127.56, 127.04, 126.84, 120.18, 120.07, 119.89, 114.03, 113.52, 113.35, 113.03, 111.75, 111.61, 111.34, 111.30, 79.65, 76.25, 75.76, 67.21, 67.11, 55.92, 55.83, 55.51, 51.99, 51.09, 50.60, 43.62, 40.31, 40.26, 40.12, 40.01, 39.63, 38.61, 38.12, 31.50, 31.25, 29.69, 28.38, 28.35, 28.26, 25.97, 25.39, 24.39, 20.97, 20.46, 12.64, 12.36.

#### Synthesis of ET-JQ1 linker-boc

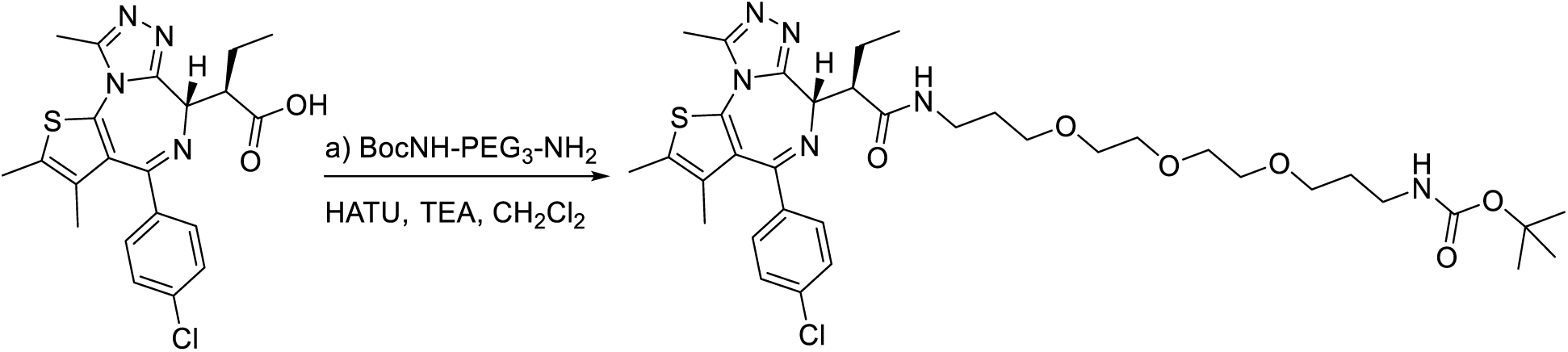

a) 17.2 mg (0.04 mmol, 1 eq.) (R)-2-((S)-4-(4-chlorophenyl)-2,3,9-trimethyl-6H-thieno[3,2-f][1,2,4]triazolo[4,3-a][1,4]diazepin-6-yl)butanoic acid was suspended in 2 mL dichloromethane. 19.2 mg (0.06 mmol, 1.5 eq.) N-Boc-4,7,10-trioxa-1,13-tridecanediamine and 55 mg (0.5 mmol, 12.5 eq.) triethylamine was added. 18.2 mg (0.048 mmol, 1.2 eq.) HATU was suspended in 1 mL dichloromethane, and was added in portion. The reaction mixture was stirred at RT overnight. Solvent was removed in vacuo, and the resulting residue was purified by preparative TLC plate (5% methanol in dichloromethane). The product (tert-butyl ((R)-16-((S)-4-(4-chlorophenyl)-2,3,9-trimethyl-6H-thieno[3,2-f][1,2,4]triazolo[4,3-a][1,4]diazepin-6-yl)-15-oxo-4,7,10-trioxa-14-azaoctadecyl)carbamate) was obtained as a white solid, yield 72%.

^1^H NMR (500 MHz, Chloroform-*d*) δ 7.38 – 7.29 (m, 4H), 6.95 (s, 1H), 5.14 (s, 1H), 4.25 (d, *J* = 9.4 Hz, 1H), 3.67 – 3.56 (m, 17H), 3.51 (q, *J* = 6.2 Hz, 3H), 3.46 (dd, *J* = 12.6, 6.3 Hz, 2H), 3.40 – 3.33 (m, 1H), 3.20 (s, 2H), 2.99 (s, 3H), 2.66 (s, 3H), 2.41 (s, 3H), 2.14 (s, 1H), 2.07 – 2.01 (m, 1H), 1.90 – 1.84 (m, 2H), 1.77 – 1.71 (m, 4H), 1.67 (s, 3H), 1.42 (s, 9H), 1.04 (t, *J* = 7.2 Hz, 3H).

^13^C NMR (126 MHz, Chloroform-*d*) δ 173.58, 163.03, 156.12, 155.03, 149.66, 136.66, 132.00, 130.94, 130.84, 130.48, 129.90, 128.59, 70.70, 70.52, 70.31, 70.18, 70.15, 70.04, 69.79, 69.48, 69.32, 59.26, 50.99, 45.35, 38.38, 37.52, 29.72, 29.60, 29.35, 29.25, 28.45, 23.23, 14.47, 13.12, 12.03, 11.84.

HRMS [M+Na]^+^: Calcd 753.3177, found 753.3163.

#### Synthesis of JQ1 linker-boc

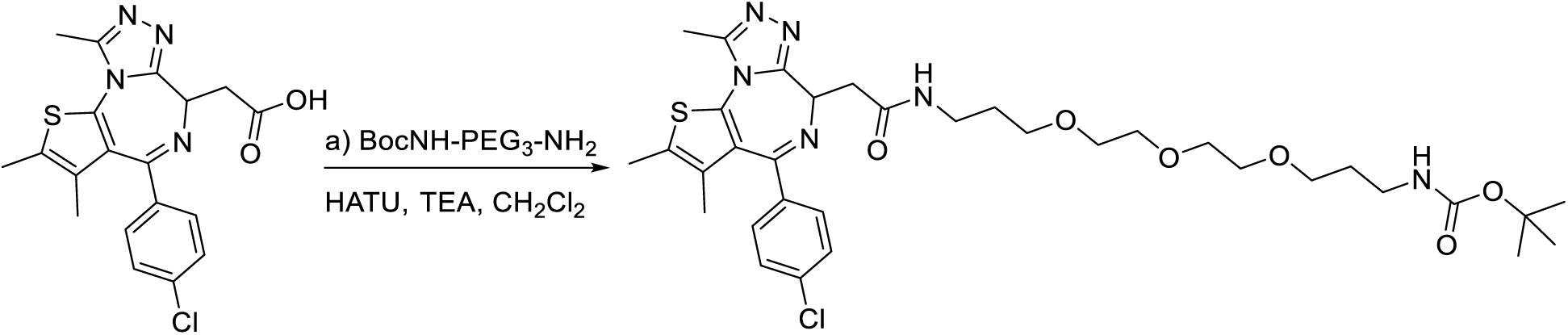

a) 16 mg (0.04 mmol, 1 eq.) 2-(4-(4-chlorophenyl)-2,3,9-trimethyl-6H-thieno[3,2-f][1,2,4]triazolo[4,3-a][1,4]diazepin-6-yl)acetic acid was suspended in 2 mL dichloromethane. 19.2 mg (0.06 mmol, 1.5 eq.) N-Boc-4,7,10-trioxa-1,13-tridecanediamine and 55 mg (0.5 mmol, 12.5 eq.) triethylamine was added. 18.2 mg (0.048 mmol, 1.2 eq.) HATU was suspended in 1 mL dichloromethane, and was added in portion. The reaction mixture was stirred at RT overnight. Solvent was removed in vacuo, and the resulting residue was purified by preparative TLC plate (5% methanol in dichloromethane). The product (tert-butyl (1-(4-(4-chlorophenyl)-2,3,9-trimethyl-6H-thieno[3,2-f][1,2,4]triazolo[4,3-a][1,4]diazepin-6-yl)-2-oxo-7,10,13-trioxa-3-azahexadecan-16-yl)carbamate) was obtained as a white solid, yield 74%.

^1^H NMR (500 MHz, Chloroform-*d*) δ 7.38 (dd, *J* = 41.0, 8.0 Hz, 4H), 7.08 (s, 1H), 5.30 (s, 1H), 4.63 (t, *J* = 6.5 Hz, 1H), 3.64 (d, *J* = 24.9 Hz, 8H), 3.55 (dd, *J* = 20.7, 5.4 Hz, 4H), 3.47 (s, 2H), 3.44 – 3.37 (m, 2H), 3.34 – 3.29 (m, 1H), 3.24 (q, *J* = 7.2 Hz, 4H), 3.20 (s, 2H), 2.67 (s, 3H), 2.41 (s, 3H), 1.84 – 1.79 (m, 2H), 1.76 – 1.72 (m, 2H), 1.68 (s, 3H), 1.42 (s, 9H), 1.35 (t, *J* = 7.2 Hz, 6H).

^13^C NMR (126 MHz, Chloroform-*d*) δ 170.66, 163.96, 156.22, 155.68, 149.96, 136.80, 136.59, 132.01, 131.06, 130.93, 130.49, 129.88, 128.71, 78.97, 70.07, 70.04, 69.46, 54.27, 53.46, 50.70, 47.36, 38.96, 38.61, 38.30, 37.53, 29.58, 28.99, 28.44, 14.39, 13.10, 11.79, 8.75.

HRMS [M+H]^+^: Calcd 703.3045, found 703.3050.

#### General strategies for fluorophore coupling

ET-JQ1-linker-boc, JQ1-linker-boc, TMP-linker-boc, or SLF’-linker-boc were suspended in 4M hydrochloric acid in 1,4-dioxane. The reaction was stirred at RT for 30 min. The solvent was removed in vacuo and the resulting residue was immediately used in the next step without purification.

#### Strategy A was applied to fluorophores in the NHS ester form

Deprotected ET-JQ1-linker-NH_2_, JQ1-linker- NH_2_, TMP-linker- NH_2_, or SLF’-linker- NH_2_ (2 eq.) was suspended in 2 mL N, N-dimethylformamide. 50 µL triethylamine (excess amount) was added, and the mixture was stirred for 10 min. 1−5 mg (1 eq.) fluorophore-NHS ester was suspended in 1 mL anhydrous DMF and was added to the reaction mixture. Additional 50 µL triethylamine was added, and the reaction mixture was stirred overnight (protected from light with aluminum foil). The solvent was removed in vacuo, and the resulting residue was purified by preparative TLC plate (1−10% methanol in dichloromethanedepending on fluorophore hydrophobicity). Yield 50%−90%.

#### Strategy B was applied to fluorophores in the free carboxylic acid form

1−5 mg (1 eq.) fluorophore carboxylic acid, deprotected ET-JQ1-linker-NH_2_, JQ1-linker- NH_2_, TMP-linker- NH_2_, or SLF’-linker- NH_2_ (2 eq.), and 50 µL triethylamine (excess amount) were suspended in 3 mL anhydrous dichloromethane. While stirring, HATU (1.1 eq.) in 1 mL dichloromethane was added in portion, and the reaction mixture was stirred overnight (protected from light with aluminum foil). The solvent was removed in vacuo, and the resulting residue was purified by preparative TLC plate (1−10% methanol in dichloromethane depending on fluorophore hydrophobicity). Yield 30%−80%.

##### ET-JQ1-TMR

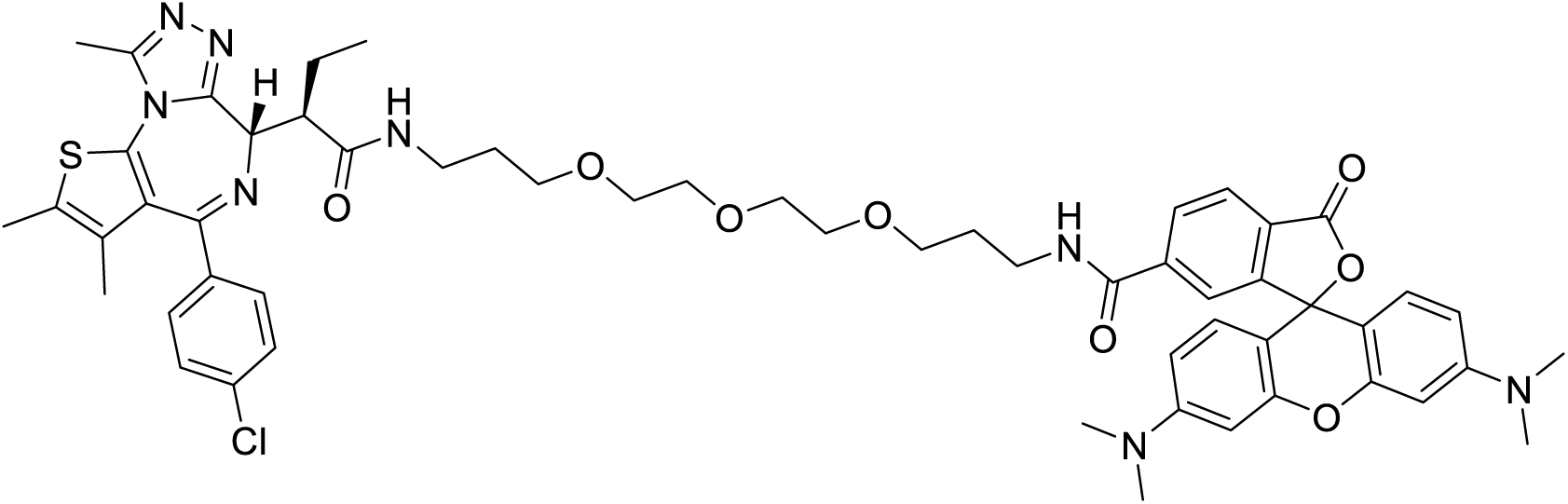

ET-JQ1-TMR was synthesized by strategy B, using 6-TAMRA. The product was obtained as a magenta solid.

^1^H NMR (500 MHz, Chloroform-*d*) δ 8.05 (d, *J* = 8.1 Hz, 1H), 7.98 (d, *J* = 8.0 Hz, 1H), 7.64 (s, 1H), 7.59 – 7.53 (m, 1H), 7.33 (d, *J* = 17.9 Hz, 4H), 6.90 (s, 1H), 6.57 (dd, *J* = 8.7, 2.9 Hz, 2H), 6.48 – 6.46 (m, 2H), 6.37 (d, J = 7.9 Hz, 2H), 4.22 (d, J = 9.7 Hz, 1H), 3.61 – 3.38 (m, 22H), 3.35 (s, 2H), 2.98 (s, 3H), 2.97 (s, 12H), 2.60 (s, 3H), 2.41 (s, 3H), 2.05 – 1.89 (m, 1H), 1.88 – 1.73 (m, 4H), 1.67 (s, 3H), 0.99 (t, J = 7.3 Hz, 3H).

^13^C NMR (126 MHz, Chloroform-*d*) δ 165.81, 162.97, 157.35, 154.99, 152.84, 152.12, 149.63, 136.67, 131.97, 130.95, 129.90, 128.78, 128.59, 128.42, 124.82, 123.02, 108.77, 106.36, 98.53, 70.33, 70.26, 70.23, 70.13, 69.98, 59.32, 53.43, 51.09, 40.28, 38.90, 37.71, 29.70, 29.27, 28.70, 23.17, 14.49, 13.13, 12.05, 11.78.

HRMS [M+Na]^+^: Calcd 1065.4070, found 1065.4066, [M+2H]^2+^: Calcd 522.2162, found 522.2157,

[M+H+Na]^2+^: Calcd 533.2072, found 533.2068.

##### ET-JQ1-JF635

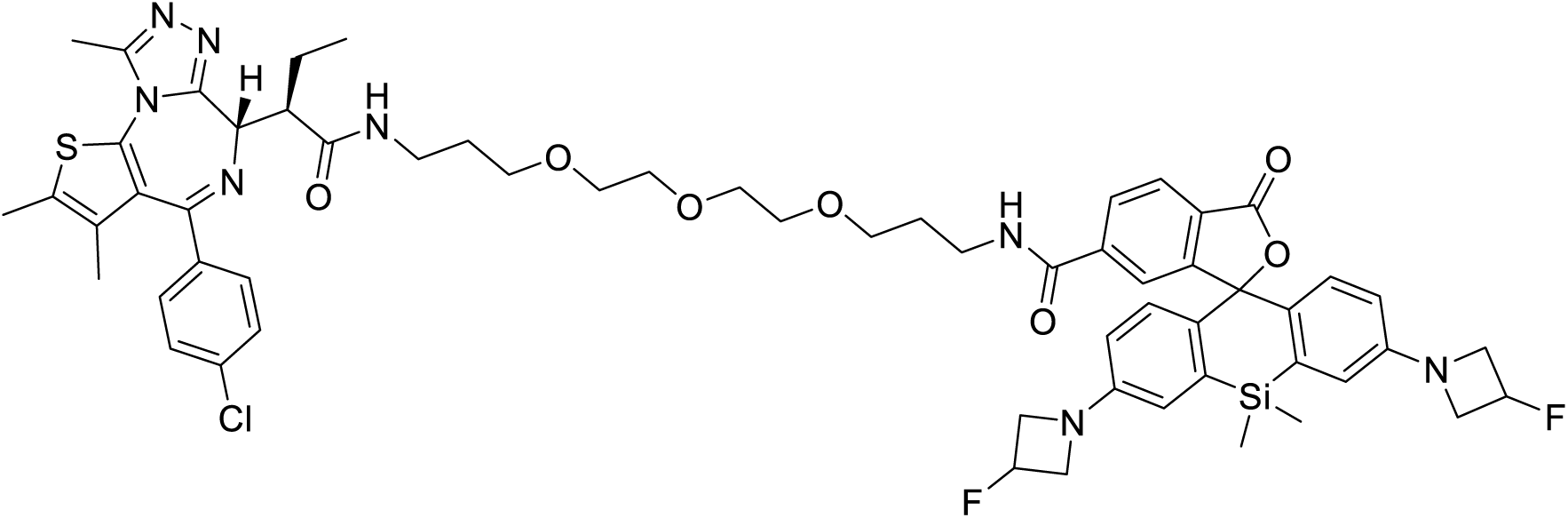

ET-JQ1-JF635 was synthesized by strategy A, using JF635 NHS ester. The product was obtained as a white solid.

^1^H NMR (500 MHz, Chloroform-*d*) δ 8.00 – 7.92 (m, 2H), 7.85 (s, 1H), 7.74 (t, *J* = 5.1 Hz, 1H), 7.38 – 7.31 (m, 4H), 6.92 (t, *J* = 5.6 Hz, 1H), 6.82 (dd, *J* = 12.8, 8.7 Hz, 2H), 6.73 – 6.69 (m, 2H), 6.30 (td, *J* = 8.8, 2.7 Hz, 2H), 5.50 – 5.34 (m, 2H), 5.14 (s, 1H), 4.26 (d, *J* = 9.9 Hz, 1H), 4.24 – 4.17 (m, 4H), 4.03 – 3.94 (m, 4H), 3.59 (qt, J = 6.4, 4.6, 3.4 Hz, 10H), 3.53 – 3.45 (m, 6H), 2.57 (s, 3H), 2.42 (s, 3H), 1.88 (q, J = 6.3 Hz, 4H), 1.69 (s, 2H), 1.03 (t, J = 7.4 Hz, 3H), 0.68 (s, 3H), 0.61 (s, 3H).

^13^C NMR (126 MHz, Chloroform-*d*) δ 173.31, 165.83, 162.89, 154.99, 154.47, 149.87, 149.60, 140.18, 136.79, 136.68, 136.65, 133.35, 131.90, 130.95, 130.83, 130.53, 129.91, 128.56, 128.46, 127.79, 127.49, 125.70, 123.85, 116.26, 112.95, 112.88, 91.63, 83.50, 81.87, 70.43, 70.24, 70.18, 59.56, 59.54, 59.41, 59.37, 51.19, 39.07, 37.94, 29.70, 29.25, 28.76, 23.10, 14.49, 13.13, 12.05, 11.72, −1.31.

HRMS [M+H]^+^: Calcd 1145.4352, found 1145.4344. [M+Na]^+^: Calcd 1167.4171, found 1167.4167.

##### ET-JQ1-AF647

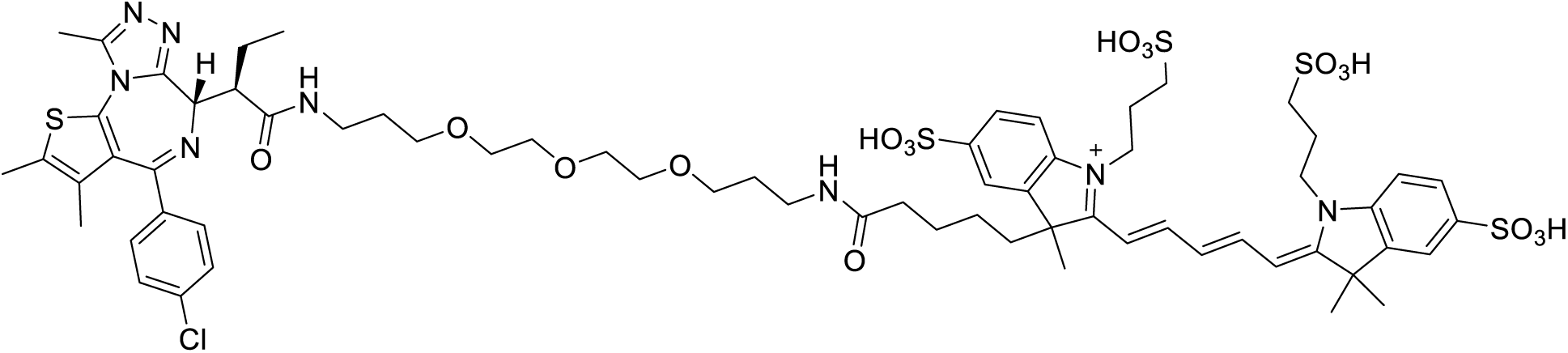

ET-JQ1-AF647 was synthesized by strategy A, using AF647 NHS ester. The product was obtained as a blue solid.

^1^H NMR (500 MHz, DMSO-*d*_6_) δ 8.36 (q, *J* = 11.6, 11.0 Hz, 2H), 8.29 (t, *J* = 4.6 Hz, 1H), 7.76 (d, *J* = 27.4 Hz, 2H), 7.67 – 7.64 (m, 1H), 7.61 (d, *J* = 7.9 Hz, 2H), 7.47 (d, *J* = 8.6 Hz, 2H), 7.40 – 7.34 (m, 4H), 6.56 (d, *J* = 12.2 Hz, 1H), 6.45 (d, *J* = 9.1 Hz, 2H), 5.02 (s, 1H), 4.27 (s, 4H), 3.70 (dd, *J* = 16.2, 4.2 Hz, 2H), 3.53 – 3.43 (m, 16H), 3.04 (d, *J* = 11.6 Hz, 2H), 3.01 – 2.89 (m, 4H), 2.58 (s, 3H), 2.40 (s, 3H), 2.00 – 1.95 (m, 4H), 1.85 (s, 4H), 1.74 – 1.71 (m, 2H), 1.67 (s, 6H), 1.60 (s, 3H), 1.45 (d, *J* = 6.5 Hz, 2H), 0.87 (t, *J* = 7.2 Hz, 3H).

^13^C NMR (126 MHz, DMSO-*d*_6_) δ 172.97, 172.13, 162.53, 156.72, 154.75, 150.15, 143.45, 145.70, 137.10, 135.69, 132.58, 131.37, 130.54, 130.10, 128.91, 126.60, 120.14, 110.36, 77.02, 70.21, 70.18, 70.00, 69.92, 68.62, 68.47, 59.68, 59.14, 55.78, 53.69, 51.51, 49.93, 49.40, 48.94, 48.38, 48.24, 36.18, 36.08, 35.79, 30.13, 29.79, 27.53, 22.88, 22.56, 14.54, 13.16, 12.12, 11.77, 8.58.

HRMS [M-2H]^2-^: Calcd 727.2090, found 727.2116. [M-3H]^3-^: Calcd 484.4702, found 484.4721.

##### JQ1-TMR

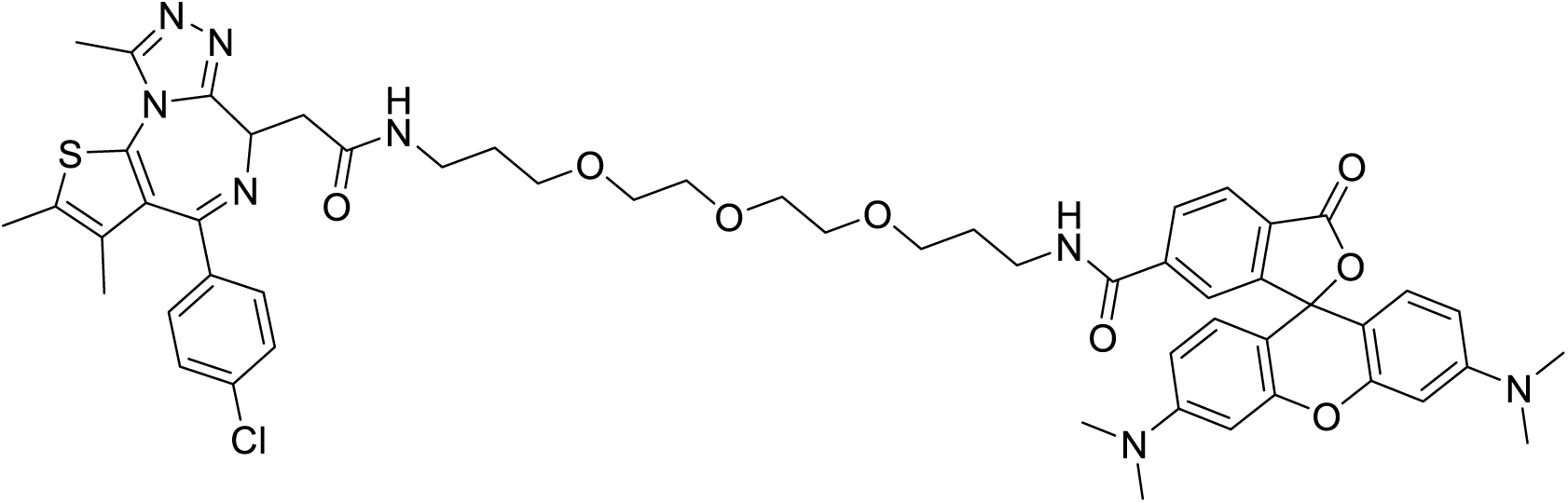

JQ1-TMR was synthesized by strategy B, using 6-TAMRA. The product was obtained as a magenta solid.

^1^H NMR (500 MHz, DMSO-*d_6_*) δ 8.68 (t, *J* = 5.5 Hz, 1H), 8.19 (t, *J* = 5.7 Hz, 1H), 8.15 (d, *J* = 8.8 Hz, 1H), 8.05 (d, *J* = 8.1 Hz, 1H), 7.63 (s, 1H), 7.48 (d, J = 8.6 Hz, 2H), 7.41 (d, J = 8.4 Hz, 2H), 6.54 – 6.48 (m, 6H), 4.51 (d, J = 6.2 Hz, 1H), 4.13 (s, 2H), 3.49 – 3.43 (m, 14H), 3.25 (s, 2H), 2.94 (s, 12H), 2.90 (s, 3H), 2.59 (s, 3H), 2.40 (s, 3H), 1.70 – 1.65 (m, 4H).

13C NMR (126 MHz, DMSO-*d_6_*) δ 169.87, 168.73, 164.92, 163.47, 155.55, 153.30, 152.60, 152.40, 150.26, 141.05, 137.20, 135.68, 132.71, 131.15, 130.56, 130.27, 130.01, 129.62, 128.93, 125.12, 122.67, 109.51, 106.09, 98.42, 85.23, 77.03, 70.18, 69.98, 69.94, 68.64, 68.48, 55.78, 54.35, 49.06, 38.10, 37.37, 36.22, 29.90, 29.57, 14.51, 13.14, 11.76.

HRMS [M+H]^+^: Calcd 1015.3938, found 1015.3939. [M+Na]^+^: Calcd 1037.3757, found 1037.3764,

[M+2H]^2+^: Calcd 508.2005, found 508.2005, [M+H+Na]^2+^: Calcd 519.1915, found 519.1915.

##### TMP-TMR

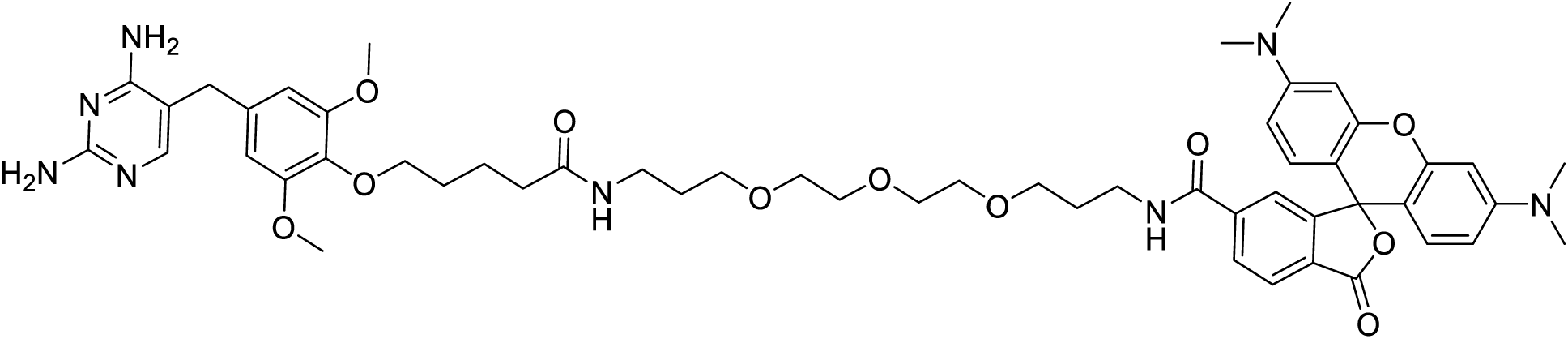

TMP-TMR was synthesized by strategy B, using 6-TAMRA. The product was obtained as a magenta solid.

^1^H NMR (500 MHz, Chloroform-*d*) δ 8.36 (s, 1H), 8.20 (d, *J* = 7.9 Hz, 1H), 7.63 (d, *J* = 22.1 Hz, 2H), 7.27 (d, *J* = 2.0 Hz, 1H), 7.23 (d, *J* = 7.9 Hz, 1H), 6.62 (d, *J* = 8.9 Hz, 2H), 6.49 (s, 2H), 6.41 (d, *J* = 9.0 Hz, 2H), 6.37 (s, 2H), 5.18 (s, 4H), 3.92 (t, *J* = 5.9 Hz, 2H), 3.75 (s, 6H), 3.70 – 3.60 (m, 12H), 3.53 – 3.50 (m, 2H), 3.48 (s, 2H), 3.30 (q, *J* = 6.0 Hz, 2H), 3.00 (s, 12H), 2.23 (t, *J* = 7.2 Hz, 2H), 1.96 – 1.90 (m, 2H), 1.79 (dd, *J* = 14.1, 6.7 Hz, 2H), 1.75 – 1.67 (m, 4H).

^13^C NMR (126 MHz, Chloroform-*d*) δ 173.14, 169.28, 165.82, 162.87, 153.69, 153.45, 152.71, 136.44, 135.91, 133.69, 133.55, 125.08, 123.46, 109.26, 107.05, 106.90, 98.27, 72.92, 70.54, 70.41, 70.37, 70.35, 70.06, 69.76, 56.16, 50.75, 40.28, 39.05, 37.54, 36.25, 34.48, 29.39, 29.15, 28.92, 22.45.

HRMS [M+H]^+^: Calcd 991.4924, found 991.4926.

##### TMP-MaP555

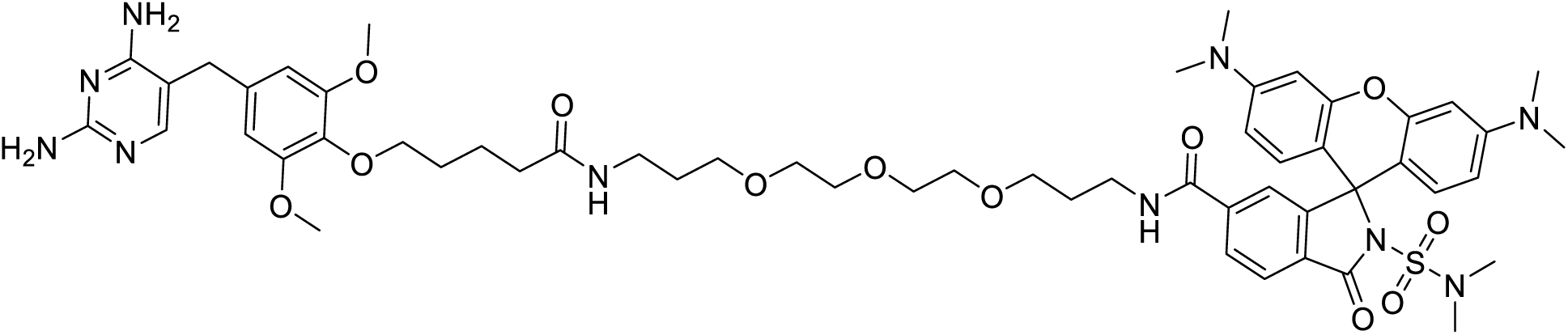

TMP-MaP555 was synthesized by strategy A, using MaP555 NHS ester. The product was obtained as a pink solid.

^1^H NMR (400 MHz, Methanol-*d_4_*) δ 8.44 (s, 1H), 7.97 – 7.87 (m, 2H), 7.43 – 7.40 (m, 1H), 7.36 (s, 1H), 6.45 (d, *J* = 8.8 Hz, 2H), 6.41 (s, 2H), 6.37 (d, *J* = 2.5 Hz, 2H), 6.33 (dd, *J* = 8.8, 2.6 Hz, 2H), 4.48 (s, 2H), 3.77 (t, *J* = 6.2 Hz, 2H), 3.66 (s, 6H), 3.53 (s, 2H), 3.39 – 3.36 (m, 4H), 3.27 (ddd, *J* = 8.9, 5.6, 2.5 Hz, 6H), 3.07 (t, *J* = 6.8 Hz, 2H), 3.03 (dt, *J* = 3.3, 1.6 Hz, 1H), 2.85 (s, 12H), 2.58 (s, 6H), 2.11 (t, *J* = 7.3 Hz, 2H), 1.70 – 1.61 (m, 4H), 1.56 (q, *J* = 7.3, 6.5 Hz, 4H).

^13^C NMR (151 MHz, Methanol-*d_4_*) δ 174.57, 168.87, 166.63, 153.44, 153.34, 140.48, 135.35, 128.39, 128.08, 123.39, 123.14, 108.19, 106.67, 105.40, 98.49, 72.50, 69.93, 69.76, 69.71, 69.44, 68.72, 68.48, 60.14, 55.17, 53.38, 39.10, 37.70, 36.84, 36.36, 35.39, 33.11, 29.11, 28.98, 28.61, 22.28.

HRMS [M+H]^+^: Calcd 1097.5135, found 1097.5125. [M+2H]^2+^: Calcd 549.2599, found 549.2606.

##### TMP-ATTO565

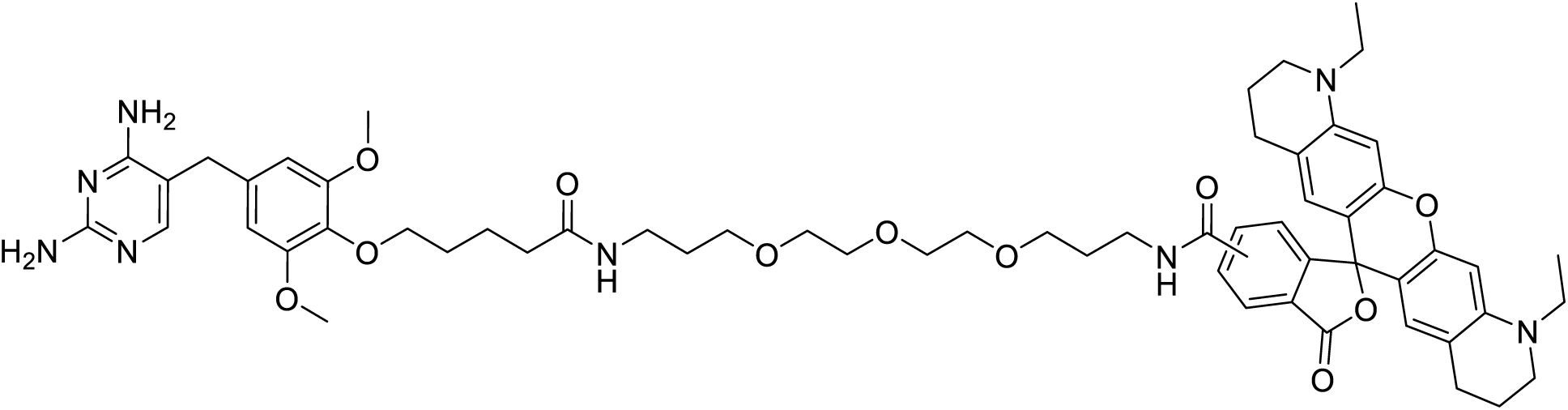

TMP-ATTO565 was synthesized by strategy A, using ATTO565 NHS ester (a mixture of 5 and 6 isomer). The product was obtained as a magenta solid.

^1^H NMR (500 MHz, Chloroform-*d*) δ 8.18 (d, J = 8.1 Hz, 1H), 7.91 (d, J = 8.0 Hz, 1H), 7.69 (s, 1H), 7.60 – 7.56 (m, 1H), 6.86 (s, 1H), 6.66 (s, 2H), 6.49 (s, 2H), 6.37 (s, 2H), 5.12 (s, 1H), 3.93 (t, J = 5.9 Hz, 2H), 3.76 (s, 6H), 3.68 – 3.64 (m, 3H), 3.61 (s, 6H), 3.57 (dd, J = 4.8, 2.9 Hz, 4H), 3.48 – 3.43 (m, 8H), 3.36 (d, J = 6.7 Hz, 4H), 3.26 (q, J = 6.3 Hz, 2H), 2.57 (tt, J = 17.0, 10.0 Hz, 4H), 2.20 (t, J = 7.0 Hz, 2H), 1.88 (ddd, J = 16.3, 11.0, 5.5 Hz, 8H), 1.71 (dt, J = 12.7, 5.4 Hz, 8H), 0.86 (dt, J = 18.3, 6.8 Hz, 6H).

^13^C NMR (126 MHz, Chloroform-*d*) δ 173.41, 166.23, 153.70, 128.70, 127.16, 105.15, 95.02, 73.01, 71.10, 70.49, 70.37, 70.33, 69.87, 69.68, 56.14, 55.99, 50.89, 48.90, 46.32, 37.12, 36.06, 34.64, 29.70, 29.38, 29.27, 28.61, 27.50, 22.37, 21.33, 10.92.

HRMS [M+H]^+^: Calcd 1071.5550, found 1071.5558. [M+2H]^2+^: Calcd 536.2811, found 536.2814.

##### TMP-JF585

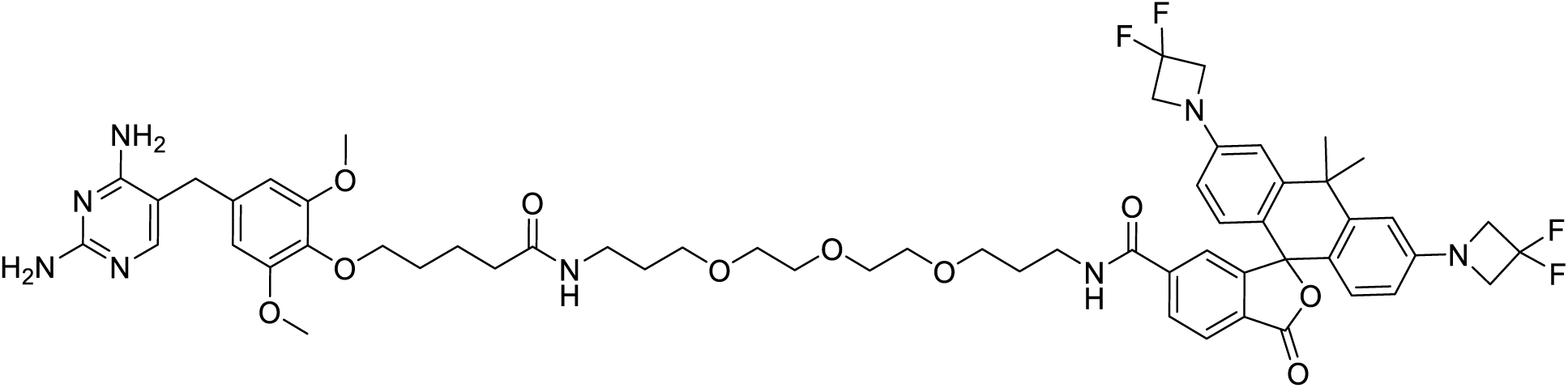

TMP-JF585 was synthesized by strategy A, using JF585 NHS ester. The product was obtained as a magenta solid.

^1^H NMR (400 MHz, Methanol-*d_4_*) δ 8.43 (s, 1H), 7.95 (s, 2H), 7.33 (t, *J* = 1.0 Hz, 1H), 7.29 (s, 1H), 6.73 (d, *J* = 2.4 Hz, 2H), 6.49 (d, *J* = 8.6 Hz, 2H), 6.44 (s, 1H), 6.42 (s, 2H), 6.31 (dd, *J* = 8.6, 2.4 Hz, 2H), 4.15 (t, *J* = 12.0 Hz, 8H), 3.78 (t, *J* = 6.2 Hz, 2H), 3.69 (s, 2H), 3.66 (s, 6H), 3.39 (q, *J* = 2.6, 2.0 Hz, 8H), 3.36 – 3.33 (m, 2H), 3.33 – 3.26 (m, 4H), 3.10 (t, *J* = 6.9 Hz, 2H), 2.11 (t, *J* = 7.3 Hz, 2H), 1.78 (s, 3H), 1.69 (dd, *J* = 12.6, 6.4 Hz, 4H), 1.65 (s, 3H), 1.61 – 1.55 (m, 4H).

^13^C NMR (101 MHz, Methanol-*d*_4_) δ 174.57, 170.31, 166.71, 163.77, 158.92, 155.88, 153.54, 150.77, 150.74, 150.72, 146.77, 140.97, 135.52, 135.43, 134.16, 133.82, 128.57, 128.52, 128.30, 124.70, 120.96, 118.84, 116.12, 113.41, 109.51, 105.57, 87.94, 72.50, 69.98, 69.94, 69.73, 69.66, 69.03, 68.71, 68.47, 68.25, 63.10, 62.84, 62.59, 55.18, 38.13, 37.53, 36.38, 35.40, 34.04, 32.86, 31.81, 29.14, 29.01, 28.75, 22.27. HRMS [M+H]^+^: Calcd 1113.5067, found 1113.5086. [M+Na]^+^: Calcd 1135.4887, found 1135.4878.

[M+2H]^2+^: Calcd 557.2570, found 557.2575. [M+H+Na]^2+^: Calcd 568.2480, found 568.2485.

##### TMP-AF594

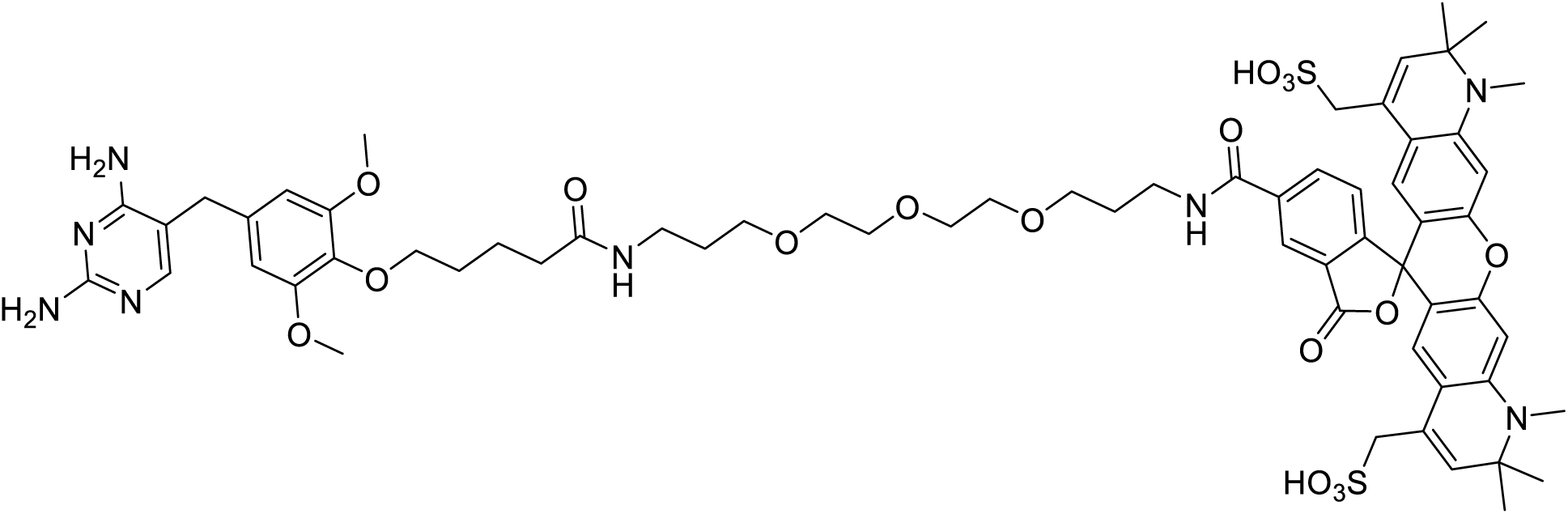

TMP-AF594 was synthesized by strategy A, using AF594 NHS ester. The product was obtained as a purple solid. ^1^H NMR (500 MHz, DMSO-*d_6_*) δ 8.96 – 8.87 (m, 1H), 8.38 (s, 1H), 8.17 (dd, *J* = 8.1, 1.3 Hz, 1H), 7.76 (t, *J* = 5.5 Hz, 1H), 7.51 (s, 1H), 7.22 (d, *J* = 8.0 Hz, 1H), 6.53 (d, *J* = 8.0 Hz, 4H), 6.29 (s, 2H), 6.06 (s, 2H), 5.66 (s, 2H), 5.53 (s, 2H), 5.02 (s, 1H), 3.76 (d, *J* = 6.1 Hz, 2H), 3.70 (s, 6H), 3.55 – 3.46 (m, 16H), 3.17 (s, 6H), 3.09 – 3.04 (m, 4H), 3.04 (s, 2H), 2.79 (s, 6H), 2.08 (t, J = 7.1 Hz, 2H), 1.85 – 1.78 (m, 2H), 1.61 (q, J = 6.7 Hz, 4H), 1.58 – 1.51 (m, 2H), 1.26 (d, J = 4.3 Hz, 12H).

^13^C NMR (126 MHz, DMSO-*d_6_*) δ 172.38, 165.33, 162.71, 162.63, 156.20, 153.31, 152.10, 147.61, 136.15, 135.29, 124.93, 123.27, 119.20, 106.29, 105.56, 72.49, 70.23, 70.04, 68.77, 68.56, 56.78, 56.31, 53.34, 49.06, 37.39, 36.16, 35.51, 33.44, 31.38, 29.87, 29.70, 28.21, 27.80, 22.31.

##### TMR-JF635

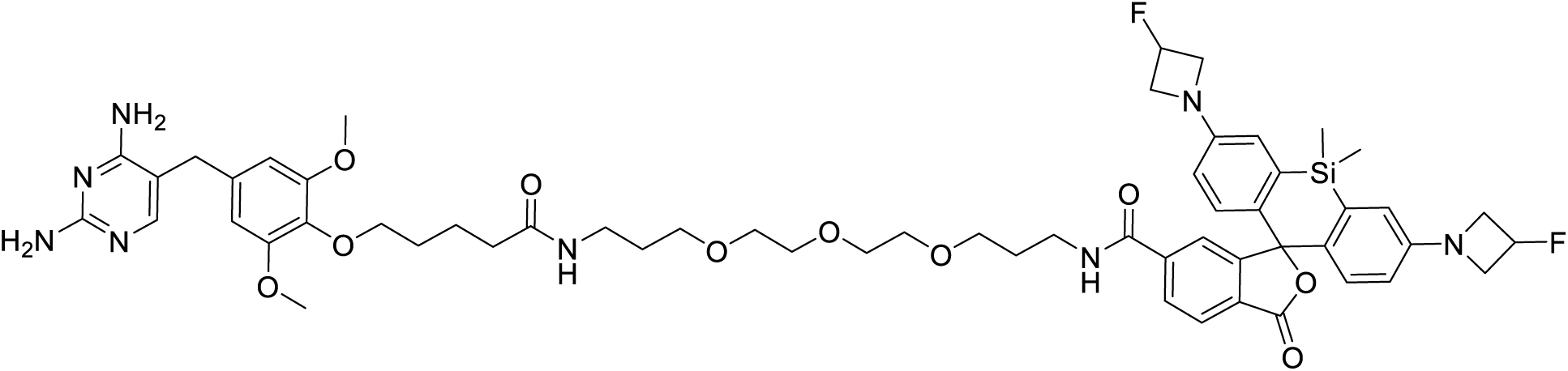

TMP-JF635 was synthesized by strategy A, using JF635 NHS ester. The product was obtained as a white solid.

^1^H NMR (500 MHz, Chloroform-*d*) δ 7.94 (s, 2H), 7.84 (s, 1H), 7.53 (s, 1H), 7.41 (s, 1H), 6.80 (d, *J* = 8.6 Hz, 2H), 6.72 (s, 2H), 6.33 (s, 2H), 6.31 (d, *J* = 8.6 Hz, 2H), 6.26 (s, 1H), 5.49 (s, 2H), 5.38 (s, 2H), 4.22 (dt, *J* = 16.4, 7.0 Hz, 4H), 4.04 – 3.94 (m, 6H), 3.79 (s, 6H), 3.67 – 3.63 (m, 3H), 3.61 (s, 4H), 3.48 – 3.44 (m, 4H), 3.32 (d, *J* = 5.9 Hz, 2H), 2.28 – 2.23 (m, 2H), 1.92 – 1.88 (m, 2H), 1.82 – 1.71 (m, 8H), 0.93 – 0.88 (m, 2H), 0.68 (s, 3H), 0.61 (s, 3H).

^13^C NMR (126 MHz, Chloroform-*d*) δ 170.00, 154.06, 149.95, 136.87, 133.11, 127.80, 127.26, 125.79, 116.27, 112.92, 105.27, 91.90, 83.50, 81.87, 72.94, 70.70, 70.29, 70.23, 70.15, 69.89, 69.62, 59.56, 59.37, 56.20, 37.39, 36.21, 29.71, 29.26, 28.68, 22.47, −1.38.

HRMS [M+H]^+^: Calcd 1093.5025, found 1093.4981

##### TMP-Cy2

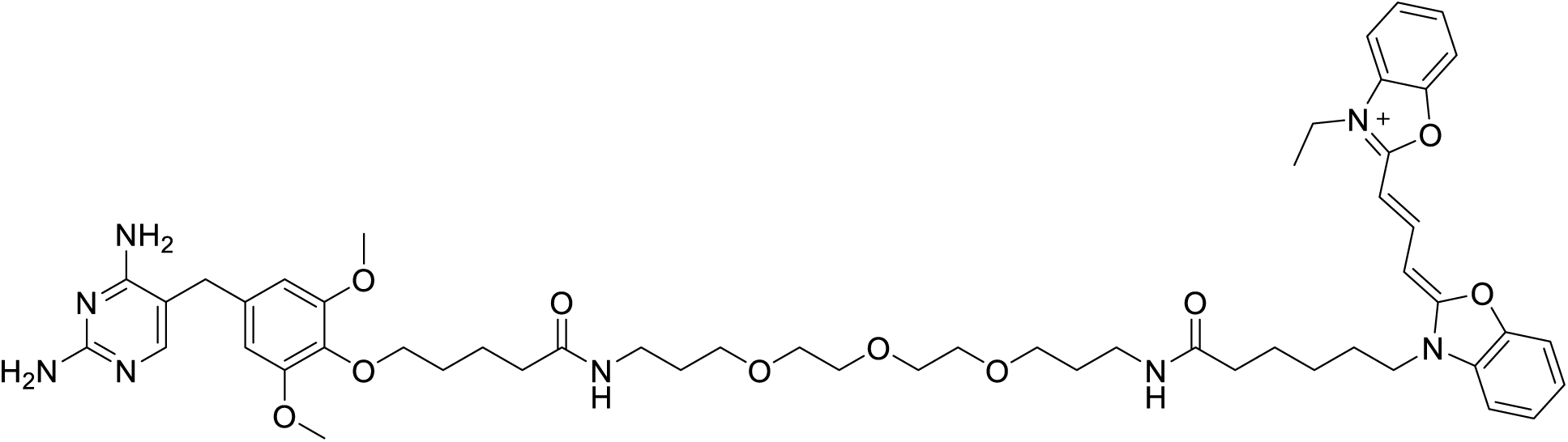

TMP-Cy2 was synthesized by strategy B, using Cy2 carboxylic acid. The product was obtained as a yellow solid.

^1^H NMR (500 MHz, Chloroform-*d*) δ 8.47 (t, J = 13.2 Hz, 1H), 7.50 (dd, J = 8.8, 7.5 Hz, 2H), 7.45 – 7.30 (m, 8H), 6.82 (d, J = 5.5 Hz, 1H), 6.49 (s, 2H), 6.36 (dd, J = 29.3, 13.2 Hz, 2H), 4.28 (q, J = 7.2 Hz, 2H), 4.19 (t, J = 7.5 Hz, 2H), 3.98 (t, J = 6.2 Hz, 2H), 3.81 (s, 6H), 3.37 (q, J = 6.0 Hz, 2H), 3.28 (q, J = 6.6 Hz, 2H), 2.29 (t, J = 7.3 Hz, 4H), 1.89 (t, J = 7.3 Hz, 2H), 1.84 – 1.69 (m, 12H), 1.52 (t, J = 7.2 Hz, 7H).

^13^C NMR (126 MHz, Chloroform-*d*) δ 173.39, 173.20, 162.01, 161.68, 153.67, 147.95, 146.97, 146.82, 130.83, 126.23, 125.21, 110.91, 110.82, 110.74, 110.23, 106.06, 70.43, 70.37, 70.15, 70.09, 70.00, 56.23, 44.57, 39.74, 37.82, 36.96, 36.13, 35.88, 29.70, 29.39, 29.18, 28.95, 26.16, 25.02, 22.50, 13.22.

HRMS [M]^+^: Calcd 979.5288, found 979.5299, [M+H]^2+^: Calcd 490.2680, found 490.2686.

##### TMP-FluH4orescein

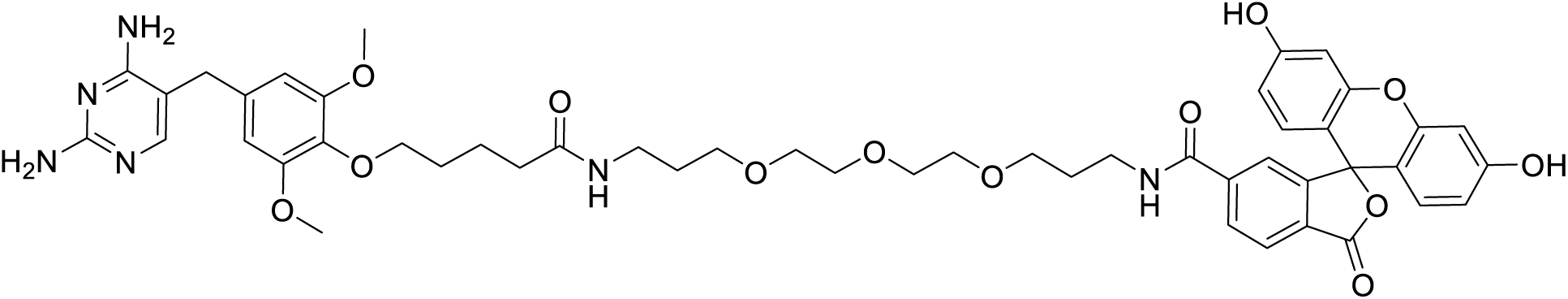

TMP-Fluorescein was synthesized by strategy A, using fluorescein NHS ester. The product was obtained as an orange solid.

^1^H NMR (500 MHz, DMSO-*d_6_*) δ 8.81 (t, *J* = 5.3 Hz, 1H), 8.45 (s, 1H), 8.24 (d, *J* = 8.9 Hz, 1H), 7.78 – 7.71 (m, 1H), 7.50 (s, 1H), 7.37 (d, *J* = 8.0 Hz, 1H), 6.69 (d, *J* = 2.0 Hz, 2H), 6.57 (s, 2H), 6.54 (s, 2H), 6.22 (s, 2H), 5.88 (s, 2H), 3.76 (t, *J* = 6.1 Hz, 2H), 3.70 (s, 6H), 3.47 (dd, *J* = 7.7, 5.0 Hz, 9H), 3.41 – 3.36 (m, 6H), 3.17 (s, 2H), 3.09 – 3.05 (m, 2H), 2.08 (t, *J* = 7.0 Hz, 2H), 1.84 – 1.76 (m, 2H), 1.64 – 1.59 (m, 4H), 1.58 – 1.53 (m, 2H).

^13^C NMR (126 MHz, DMSO-*d_6_*) δ 172.40, 168.67, 165.04, 162.79, 162.12, 160.22, 154.88, 153.32, 152.31, 136.79, 135.92, 135.32, 135.08, 129.60, 126.99, 124.71, 123.72, 109.54, 106.47, 106.32, 102.75, 72.49, 70.26, 70.23, 70.06, 70.00, 68.65, 68.56, 56.30, 49.06, 37.32, 36.18, 35.53, 29.71, 29.65, 22.32.

HRMS [M+H]^+^: Calcd 973.3978, found 973.3954.

##### TMP-AF488

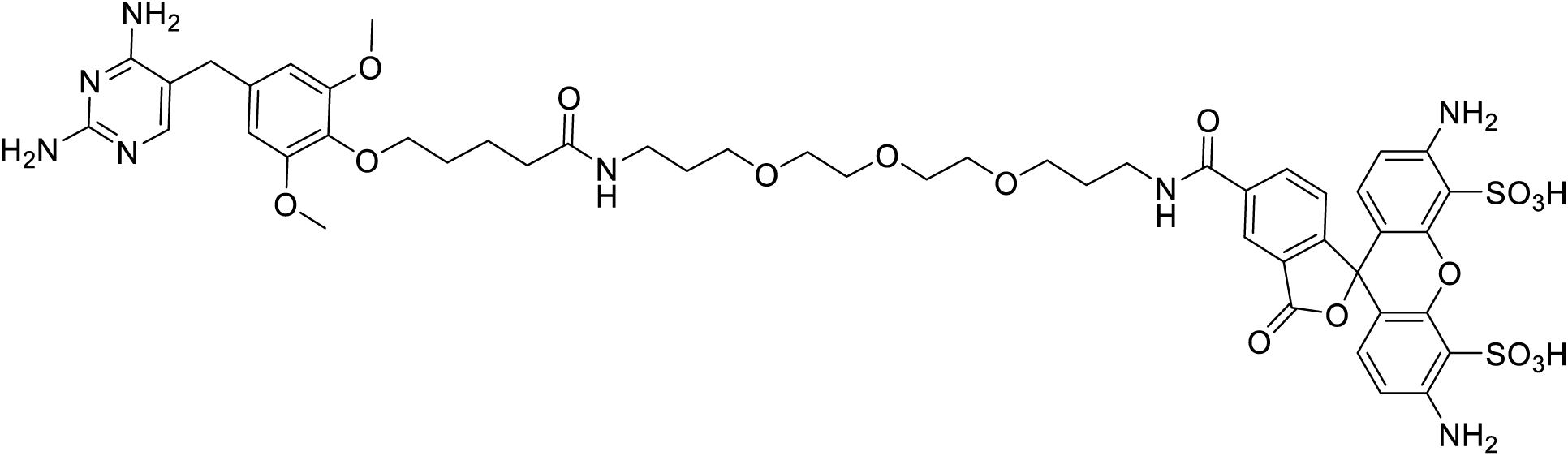

TMP-Fluorescein was synthesized by strategy A, using AF488 NHS ester. The product was obtained as an orange solid.

^1^H NMR (500 MHz, DMSO-*d_6_*) δ 8.71 (t, *J* = 5.3 Hz, 1H), 8.45 (s, 1H), 8.10 (d, *J* = 8.0 Hz, 1H), 7.68 (t, *J* = 5.5 Hz, 2H), 7.60 (s, 2H), 7.24 (d, *J* = 7.9 Hz, 1H), 7.19 (s, 1H), 6.59 (d, *J* = 8.6 Hz, 2H), 6.53 (s, 1H), 6.50 (s, 2H), 3.72 (t, *J* = 6.1 Hz, 2H), 3.65 (s, 6H), 3.46 (d, *J* = 5.8 Hz, 8H), 3.41 (d, *J* = 6.2 Hz, 6H), 3.01 (q, *J* = 6.5 Hz, 2H), 2.02 (t, *J* = 7.1 Hz, 2H), 1.73 (p, *J* = 6.4 Hz, 2H), 1.54 (ddq, *J* = 18.6, 14.1, 7.4, 7.0 Hz, 6H).

^13^C NMR (126 MHz, DMSO-*d_6_*) δ 172.46, 168.82, 165.35, 164.05, 163.55, 153.50, 147.51, 146.84, 144.87, 136.32, 135.63, 130.16, 127.62, 123.90, 114.41, 108.73, 106.72, 72.50, 70.29, 70.27, 70.09, 68.73, 68.57, 56.35, 40.44, 37.33, 36.21, 35.53, 32.83, 29.86, 29.75, 29.62, 22.33.

HRMS [M]^-^: Calcd 1093.3289, found 1093.3293. [M-2H+Na]^-^: Calcd 1115.3108, found 1115.3116.

##### TMP-Rh110

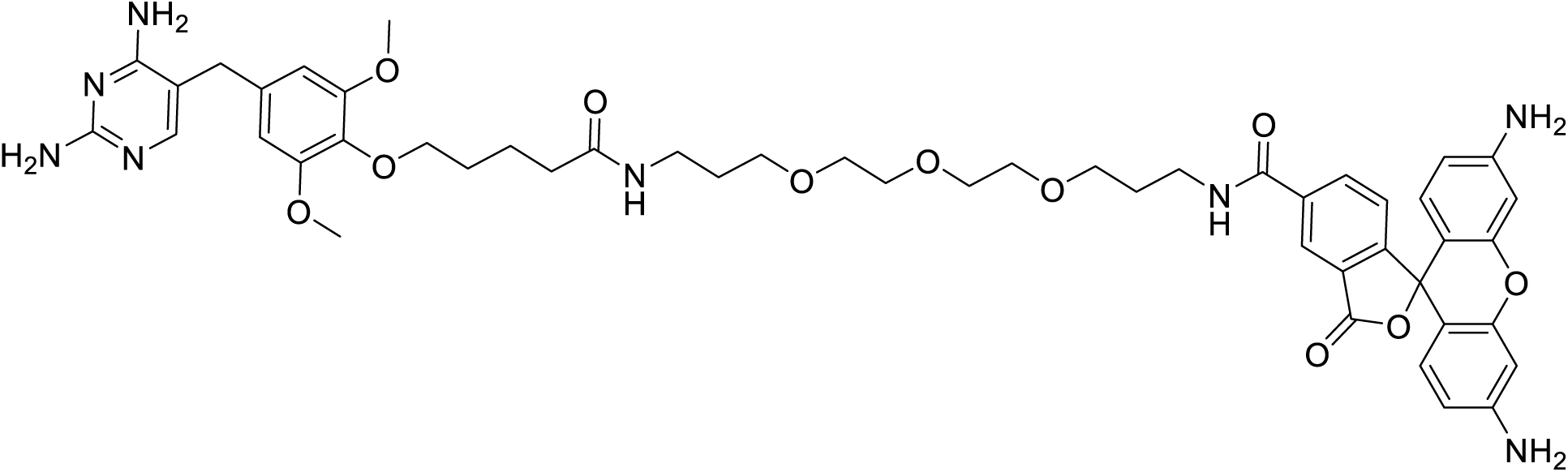

TMP-Rh110 was synthesized by strategy A, using rhodamine 110 NHS ester. The product was obtained as an orange solid.

^1^H NMR (500 MHz, DMSO-*d*_6_) δ 8.80 (s, 1H), 8.40 (s, 1H), 8.22 (d, *J* = 7.8 Hz, 1H), 7.75 (t, *J* = 4.7 Hz, 1H), 7.51 (s, 1H), 7.32 (d, *J* = 8.0 Hz, 1H), 6.54 (s, 2H), 6.40 (s, 2H), 6.36 – 6.27 (m, 4H), 6.07 (s, 2H), 5.67 (s, 2H), 5.62 (s, 4H), 3.76 (t, *J* = 5.7 Hz, 2H), 3.70 (s, 6H), 3.52 (m, 8H), 3.17 (s, 7H), 3.10 – 3.05 (m, 2H), 2.08 (t, *J* = 6.9 Hz, 2H), 1.80 (q, *J* = 6.3 Hz, 2H), 1.66 – 1.59 (m, 4H), 1.59 – 1.54 (m, 2H).

^13^C NMR (126 MHz, DMSO-*d*_6_) δ 165.17, 162.67, 156.15, 153.30, 152.69, 151.72, 136.14, 128.95, 111.32, 106.28, 105.88, 99.61, 72.49, 70.22, 70.05, 69.99, 68.67, 68.56, 56.30, 49.06, 37.32, 36.19, 33.43, 29.87, 29.71, 29.64, 22.32.

HRMS [M+H]^+^: Calcd 935.4298, found 935.4313. [M+2H]^2+^: Calcd 468.2185, found 468.2191.

##### TMP-BODIPY1

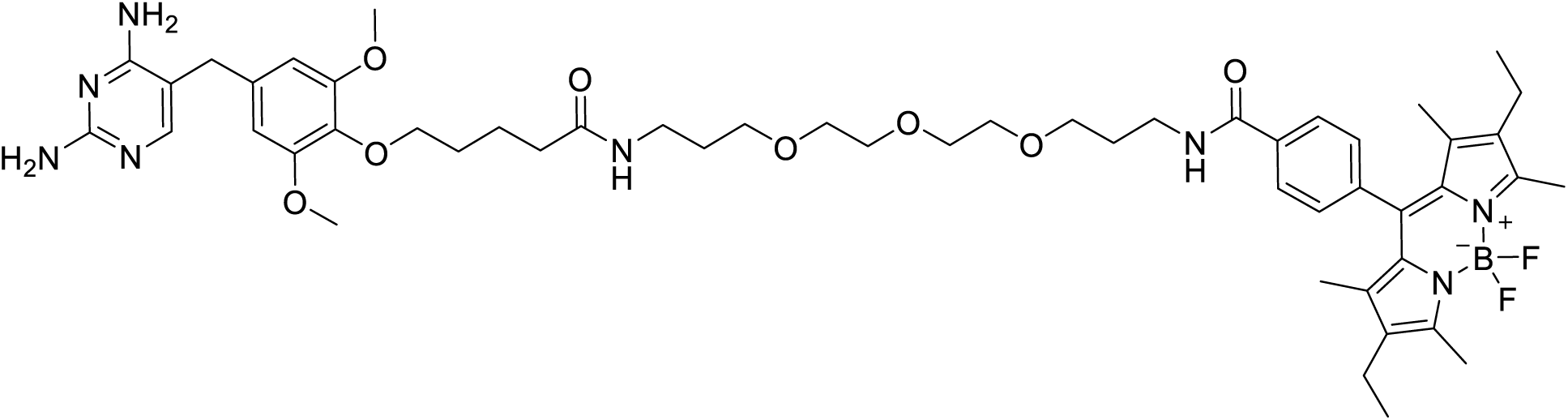

TMP-BODIPY1 was synthesized by strategy B, using BODIPY1 carboxylic acid. The product was obtained as a dark red solid.

^1^H NMR (500 MHz, Chloroform-*d*) δ 7.98 (d, *J* = 7.8 Hz, 2H), 7.77 (s, 1H), 7.41 (s, 1H), 7.36 (d, *J* = 7.8 Hz, 2H), 6.37 (s, 2H), 6.22 (s, 1H), 4.69 (s, 2H), 4.58 (s, 2H), 3.94 (t, *J* = 5.7 Hz, 2H), 3.77 (s, 6H), 3.69 (d, *J* = 5.4 Hz, 3H), 3.66 (s, 4H), 3.64 – 3.60 (m, 7H), 3.54 (s, 2H), 3.45 (d, *J* = 5.8 Hz, 2H), 3.28 (q, *J* = 6.2 Hz, 2H), 2.53 (s, 6H), 1.94 (p, *J* = 5.3 Hz, 2H), 1.89 – 1.83 (m, 2H), 1.82 – 1.76 (m, 4H), 1.75 (d, *J* = 8.1 Hz, 2H), 1.69 (p, *J* = 5.8 Hz, 2H), 1.25 (s, 6H), 0.97 (t, *J* = 7.5 Hz, 6H).

^13^C NMR (126 MHz, Chloroform-*d*) δ 173.05, 166.24, 162.67, 162.24, 156.92, 154.10, 153.73, 139.02, 138.86, 138.18, 135.90, 134.91, 133.83, 133.01, 130.41, 128.58, 127.80, 106.46, 105.03, 72.96, 70.83, 70.47, 70.45, 70.27, 70.05, 69.77, 56.11, 39.15, 37.50, 36.31, 34.71, 29.48, 29.26, 28.87, 22.45, 17.06, 14.61, 12.54, 11.88.

HRMS [M+H]^+^: Calcd 985.5529, found 985.5539. [M+2H]^2+^: Calcd 493.2801, found 493.2802.

##### TMP-BODIPY2

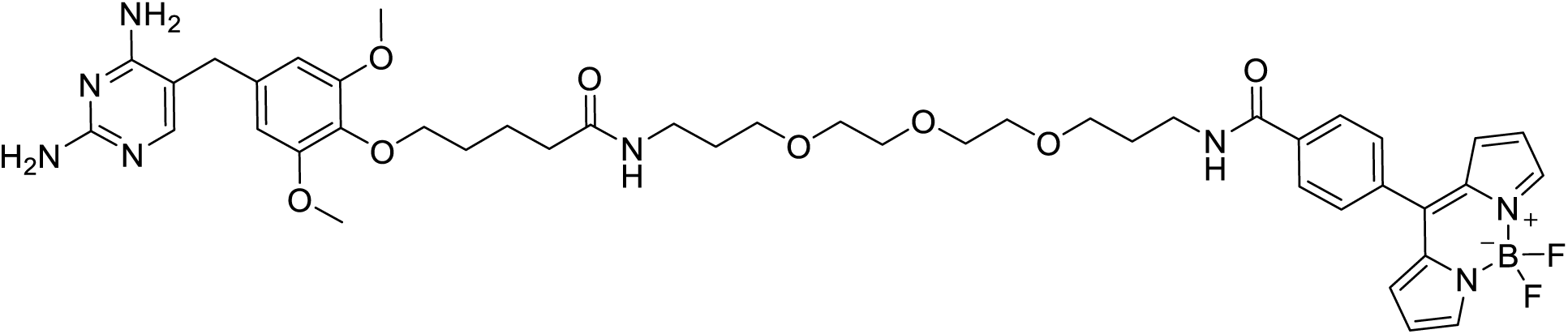

TMP-BODIPY2 was synthesized by strategy B, using BODIPY2 carboxylic acid. The product was obtained as a dark red solid.

^1^H NMR (500 MHz, Chloroform-*d*) δ 7.91 (s, 2H), 7.76 (s, 1H), 7.52 (d, *J* = 8.5 Hz, 2H), 7.03 (d, *J* = 8.5 Hz, 2H), 6.97 (d, *J* = 3.7 Hz, 2H), 6.57 – 6.52 (m, 2H), 6.44 (t, *J* = 4.8 Hz, 1H), 6.37 (s, 2H), 6.27 (t, *J* = 4.9 Hz, 1H), 4.74 (s, 2H), 4.62 (s, 2H), 4.10 (t, *J* = 6.1 Hz, 2H), 3.94 (t, *J* = 5.9 Hz, 2H), 3.77 (s, 6H), 3.63 (s, 6H), 3.60 – 3.57 (m, 4H), 3.54 (dt, *J* = 10.1, 5.9 Hz, 4H), 3.48 (s, 1H), 3.35 (dq, *J* = 12.6, 6.2 Hz, 4H), 2.39 (t, *J* = 7.2 Hz, 2H), 2.26 (t, *J* = 7.1 Hz, 2H), 2.16 (p, *J* = 6.6 Hz, 2H), 1.78 (dq, *J* = 13.0, 6.4 Hz, 8H).

^13^C NMR (126 MHz, Chloroform-*d*) δ 173.21, 172.26, 162.67, 162.24, 161.46, 153.68, 147.42, 143.35, 135.83, 134.78, 133.89, 131.35, 126.24, 114.55, 106.43, 105.05, 72.95, 70.35, 69.96, 69.94, 69.83, 69.66, 67.40, 56.11, 37.67, 37.40, 36.26, 34.67, 32.58, 29.42, 29.23, 29.04, 25.07, 22.47.

HRMS [M+H]^+^: Calcd 895.4096, found 895.4098.

##### TMP-JF525

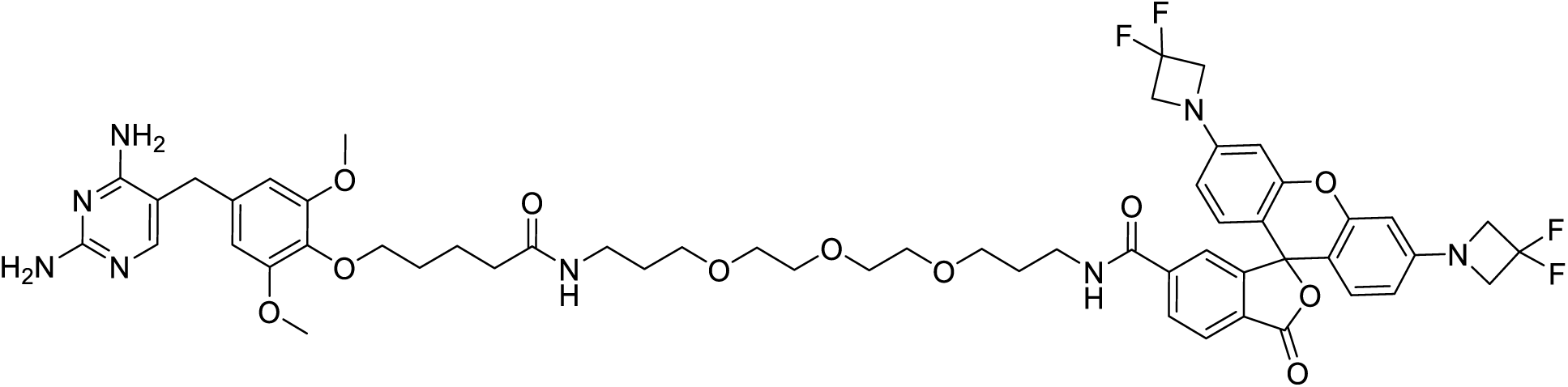

TMP-JF525 was synthesized by strategy A, using JF585 NHS ester. The product was obtained as a magenta solid.

^1^H NMR (400 MHz, Methanol-*d_4_*) δ 8.04 – 8.00 (m, 1H), 7.97 (dd, *J* = 8.0, 0.6 Hz, 1H), 7.53 (d, *J* = 2.0 Hz, 1H), 7.14 (s, 1H), 6.63 (d, *J* = 8.7 Hz, 2H), 6.44 (s, 2H), 6.39 (d, *J* = 2.3 Hz, 2H), 6.28 (dd, *J* = 8.7, 2.4 Hz, 2H), 4.23 (t, *J* = 11.9 Hz, 8H), 3.79 (t, *J* = 6.1 Hz, 2H), 3.68 (s, 6H), 3.54 (s, 2H), 3.41 (d, *J* = 4.3 Hz, 8H), 3.33 (ddt, *J* = 12.4, 6.2, 2.8 Hz, 6H), 3.10 (td, *J* = 7.1, 2.8 Hz, 4H), 2.12 (t, *J* = 7.3 Hz, 2H), 1.74 – 1.62 (m, 4H), 1.58 (p, *J* = 6.4 Hz, 4H).

^13^C NMR (126 MHz, Chloroform-*d*) δ 173.23, 165.56, 155.40, 154.04, 152.55, 151.51, 141.02, 136.40, 130.71, 129.17, 128.39, 123.57, 115.53, 109.05, 105.50, 99.19, 99.17, 72.84, 70.41, 70.18, 70.07, 69.91, 69.52, 63.30, 56.25, 45.85, 39.13, 37.35, 36.19, 34.05, 29.70, 29.37, 29.30, 28.69, 22.47.

HRMS [M+H]^+^: Calcd 1087.4547, found 1087.4531. [M+Na]^+^: Calcd 1109.4366, found 1109.4351.

[M+2H]^2+^: Calcd 544.2310, found 544.2311. [M+H+Na]^2+^: Calcd 555.2220, found 555.2218.

##### SLF’-TMR

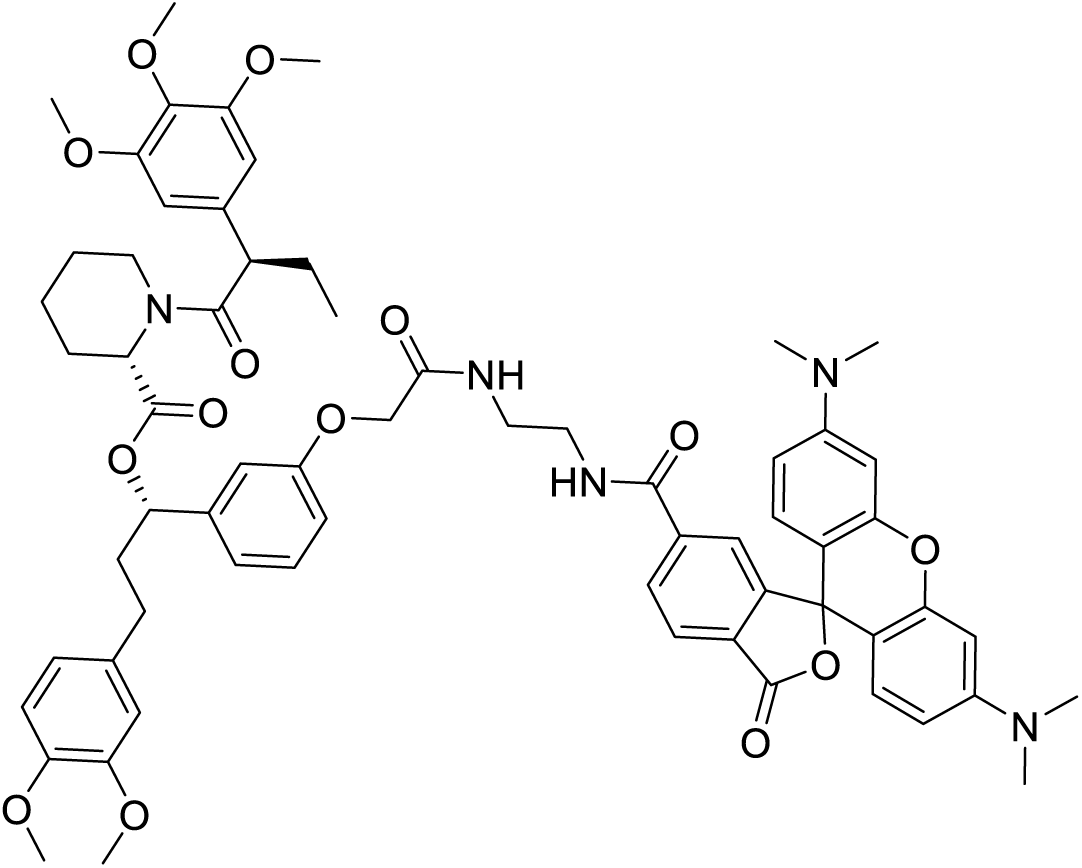

SLF’-TMR was synthesized by strategy B, using 6-TAMRA. The product was obtained as a magenta solid.

^1^H NMR (500 MHz, Chloroform-*d*) δ 8.44 (s, 1H), 8.08 (d, *J* = 8.0 Hz, 1H), 8.02 (s, 1H), 7.59 (s, 1H), 7.19 (d, *J* = 8.0 Hz, 1H), 7.13 (t, *J* = 7.9 Hz, 1H), 6.77 (t, *J* = 9.0 Hz, 2H), 6.72 (s, 1H), 6.64 (d, *J* = 7.0 Hz, 2H), 6.61 (d, *J* = 8.8 Hz, 2H), 6.57 (d, *J* = 8.8 Hz, 1H), 6.48 (s, 2H), 6.42 (s, 2H), 6.39 (d, *J* = 8.6 Hz, 2H), 5.65 – 5.59 (m, 1H), 5.43 (d, *J* = 3.4 Hz, 1H), 4.48 (s, 2H), 3.99 (s, 1H), 3.84 (s, 7H), 3.82 (s, 2H), 3.77 (s, 3H), 3.67 (s, 6H), 3.61 (d, *J* = 7.4 Hz, 4H), 2.98 (s, 13H), 2.83 (t, *J* = 12.5 Hz, 1H), 2.55 (ddd, *J* = 14.8, 7.1, 3.2 Hz, 1H), 2.51 – 2.42 (m, 1H), 2.30 (d, *J* = 12.9 Hz, 1H), 2.11 – 2.02 (m, 2H), 1.93 (dt, *J* = 14.1, 7.0 Hz, 1H), 1.71 (dt, *J* = 12.7, 6.8 Hz, 3H), 1.61 (d, *J* = 12.5 Hz, 1H), 1.44 (q, *J* = 13.0 Hz, 1H), 0.89 (t, *J* = 7.2 Hz, 3H).

^13^C NMR (126 MHz, Chloroform-*d*) δ 172.90, 170.55, 170.03, 169.03, 166.55, 157.46, 153.15, 152.83, 152.38, 152.15, 148.87, 147.35, 142.18, 136.64, 135.74, 135.59, 135.28, 133.63, 133.34, 129.75, 128.87, 128.64, 126.47, 124.60, 124.34, 123.88, 120.20, 119.91, 113.87, 112.55, 111.71, 111.28, 108.91, 108.70, 106.64, 105.96, 105.10, 98.55, 98.42, 75.64, 67.23, 60.81, 56.32, 56.01, 55.92, 55.85, 52.20, 50.70, 43.56, 41.22, 40.26, 39.70, 38.12, 31.35, 29.71, 28.29, 26.80, 25.30, 20.89, 12.57.

HRMS [M+H]^+^: Calcd 1148.5227, found 1148.5255.

##### SLF’-MaP555

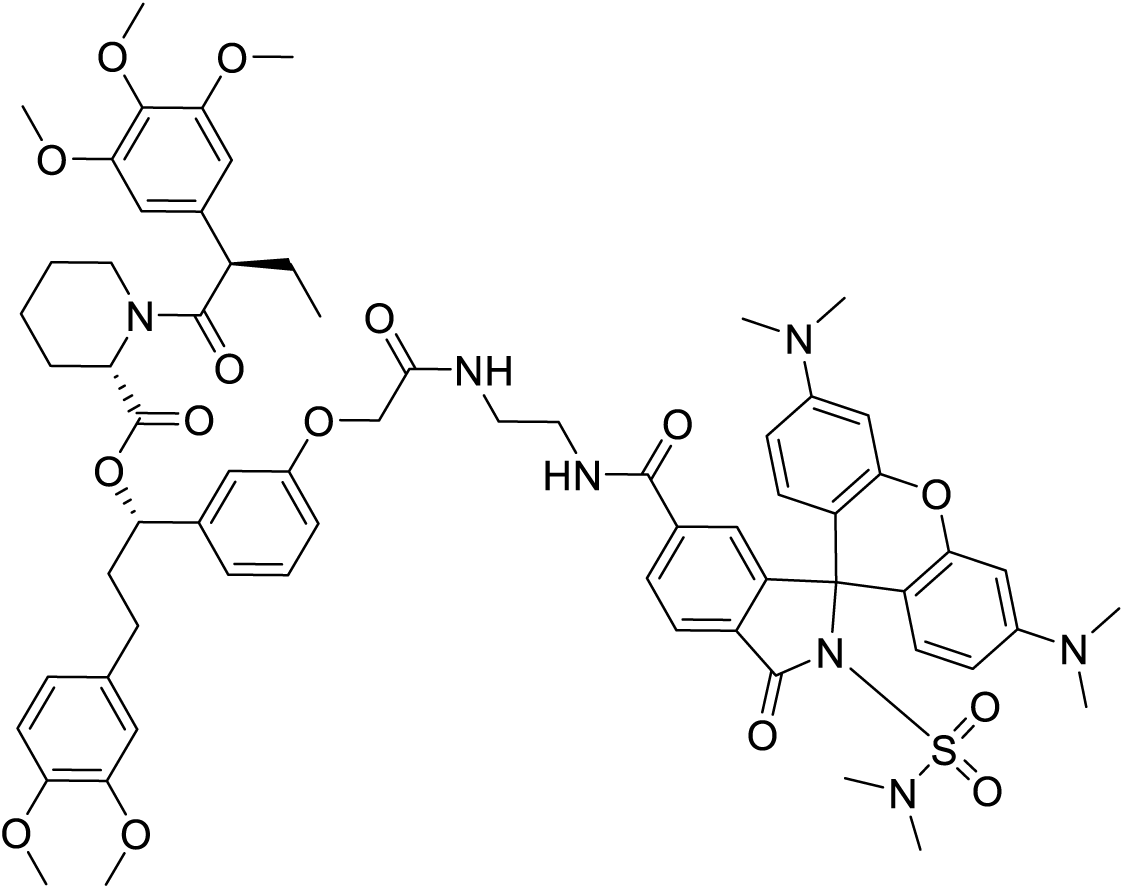

SLF’-MaP555 was synthesized by strategy A, using MaP555 NHS ester. The product was obtained as a pink solid.

^1^H NMR (500 MHz, Chloroform-*d*) δ 7.93 (d, *J* = 8.0 Hz, 1H), 7.82 (d, *J* = 8.1 Hz, 1H), 7.50 (s, 1H), 7.36 – 7.30 (m, 2H), 7.11 (t, *J* = 7.8 Hz, 1H), 6.77 (d, *J* = 8.0 Hz, 1H), 6.74 – 6.62 (m, 6H), 6.54 (dd, *J* = 8.4, 5.9 Hz, 3H), 6.46 (s, 2H), 6.39 (s, 2H), 6.31 (dd, *J* = 7.3, 4.0 Hz, 2H), 5.66 – 5.60 (m, 1H), 5.42 (d, *J* = 4.2 Hz, 1H), 4.44 (s, 2H), 3.84 (s, 7H), 3.83 (s, 2H), 3.76 (s, 3H), 3.65 (s, 6H), 3.60 – 3.56 (m, 2H), 3.52 (s, 3H), 3.49 (s, 2H), 2.94 (s, 12H), 2.72 (s, 6H), 2.56 (ddd, *J* = 14.7, 10.1, 5.8 Hz, 2H), 2.51 – 2.44 (m, 1H), 2.30 (d, *J* = 13.0 Hz, 1H), 2.05 (tq, *J* = 15.0, 7.9 Hz, 3H), 1.92 (dq, *J* = 14.9, 8.0, 6.5 Hz, 1H), 1.71 (dt, *J* = 13.2, 7.0 Hz, 4H), 1.49 – 1.38 (m, 2H), 0.89 (t, *J* = 7.0 Hz, 3H).

^13^C NMR (126 MHz, Chloroform-*d*) δ 172.80, 170.59, 170.07, 166.52, 166.47, 157.27, 154.19, 153.21, 153.15, 151.54, 148.88, 147.37, 142.32, 140.00, 136.62, 135.25, 133.28, 130.56, 129.73, 128.49, 127.35, 123.97, 123.85, 120.20, 120.03, 113.61, 112.67, 111.71, 111.29, 106.53, 106.50, 105.02, 98.85, 69.01, 67.24, 60.80, 55.96, 55.92, 55.85, 52.15, 50.77, 43.51, 41.26, 40.22, 39.26, 38.12, 37.90, 31.34, 29.70, 28.32, 26.75, 25.28, 20.85, 12.57.

HRMS [M+H]^+^: Calcd 1254.5428, found 1254.5448.

##### SLF’-JF635

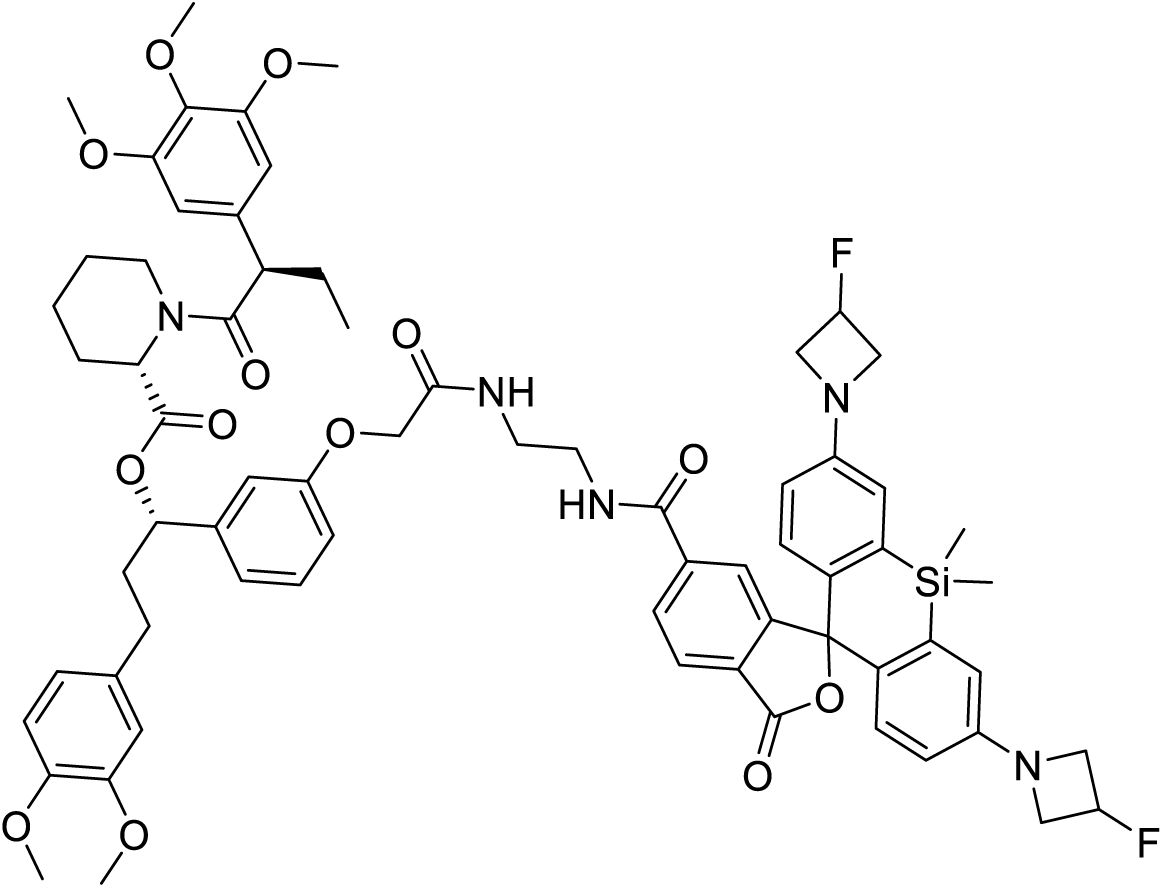

SLF’-JF635 was synthesized by strategy A, using JF635 NHS ester. The product was obtained as a white solid.

^1^H NMR (500 MHz, Chloroform-*d*) δ 7.93 (d, J = 7.9 Hz, 1H), 7.82 – 7.75 (m, 3H), 7.37 (d, J = 5.3 Hz, 1H), 7.06 – 7.00 (m, 1H), 6.80 (ddd, J = 20.1, 12.0, 8.3 Hz, 4H), 6.70 (dd, J = 6.8, 3.1 Hz, 4H), 6.67 – 6.64 (m, 2H), 6.40 (d, J = 21.9 Hz, 2H), 6.29 (ddd, J = 8.4, 5.1, 2.7 Hz, 2H), 5.85 – 5.58 (m, 1H), 5.46 (tt, J = 5.8, 3.6 Hz, 1H), 5.43 – 5.38 (m, 1H), 5.34 (tt, J = 5.9, 3.3 Hz, 1H), 4.49 (s, 2H), 4.22 – 4.14 (m, 4H), 4.02 – 3.97 (m, 2H), 3.94 (td, J = 9.4, 8.4, 5.0 Hz, 2H), 3.76 (s, 2H), 3.64 (s, 6H), 3.35 (s, 1H), 2.83 (td, J = 13.3, 3.1 Hz, 1H), 2.57 (ddd, J = 14.5, 9.5, 5.3 Hz, 1H), 2.48 (ddd, J = 14.2, 9.2, 6.7 Hz, 1H), 2.30 (d, J = 13.6 Hz, 1H), 2.12 – 1.99 (m, 2H), 1.97 – 1.88 (m, 1H), 1.71 (dt, J = 14.0, 7.0 Hz, 4H), 0.91 – 0.87 (m, 6H), 0.84 (d, J = 7.4 Hz, 3H), 0.67 (d, J = 4.6 Hz, 3H), 0.59 (d, J = 2.2 Hz, 3H).

^13^C NMR (126 MHz, Chloroform-*d*) δ 172.99, 170.63, 170.48, 169.85, 166.35, 157.27, 154.74, 153.16, 149.90, 148.90, 147.40, 142.40, 139.19, 136.71, 136.64, 135.19, 133.30, 133.27, 129.73, 128.51, 127.72, 126.98, 125.99, 124.01, 120.21, 120.07, 116.27, 113.59, 113.00, 112.97, 112.40, 111.71, 111.30, 105.03, 104.64, 91.57, 83.50, 81.88, 75.58, 67.28, 60.81, 59.55, 59.37, 56.34, 55.97, 55.93, 55.86, 52.22, 50.78, 43.54, 41.74, 39.27, 38.19, 31.94, 31.38, 29.71, 29.67, 29.37, 28.28, 26.74, 25.24, 22.70, 20.80, 14.13, 12.56, −1.21, −1.25.

HRMS [M+H]+: Calcd 1250.5328, found 1250.5336.

##### SLF’’-TMR

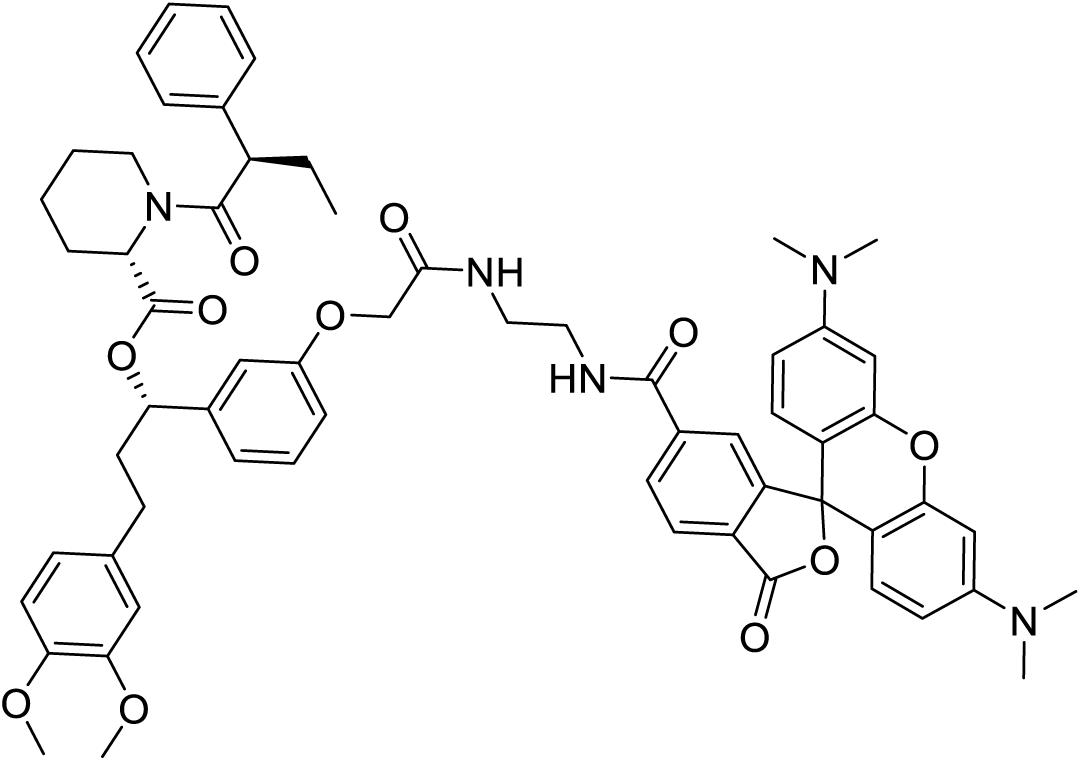

SLF’’-TMR was synthesized by strategy B, using 6-TAMRA. The product was obtained as a magenta solid. ^1^H NMR (500 MHz, Chloroform-*d*) δ 8.00 – 7.91 (m, 1H), 7.74 – 7.52 (m, 2H), 7.36 – 7.30 (m, 1H), 7.28 (s, 1H), 7.24 – 7.15 (m, 4H), 7.01 – 6.33 (m, 12H), 5.74 (ddd, J = 85.3, 8.0, 5.6 Hz, 1H), 5.44 – 4.59 (m, 1H), 4.56 – 4.32 (m, 2H), 3.87 – 3.83 (m, 6H), 3.69 – 3.44 (m, 5H), 2.97 (d, J = 5.2 Hz, 12H), 2.79 – 1.33 (m, 12H), 1.21 (d, J = 13.0 Hz, 1H), 0.86 (t, J = 7.3 Hz, 3H).

^13^C NMR (126 MHz, Chloroform-*d*) δ 172.90, 170.53, 169.81, 169.15, 157.22, 152.94, 152.91, 152.22, 152.19, 148.87, 147.35, 142.23, 139.50, 133.44, 129.71, 129.01, 128.75, 128.63, 128.05, 127.85, 127.53, 126.89, 125.12, 123.50, 120.15, 120.09, 114.58, 112.01, 111.71, 111.31, 106.41, 106.38, 98.59, 98.54, 67.22, 55.93, 55.85, 52.21, 50.60, 43.62, 40.28, 38.29, 31.37, 28.24, 26.70, 25.32, 20.95, 12.40.

##### SLF’-AF594

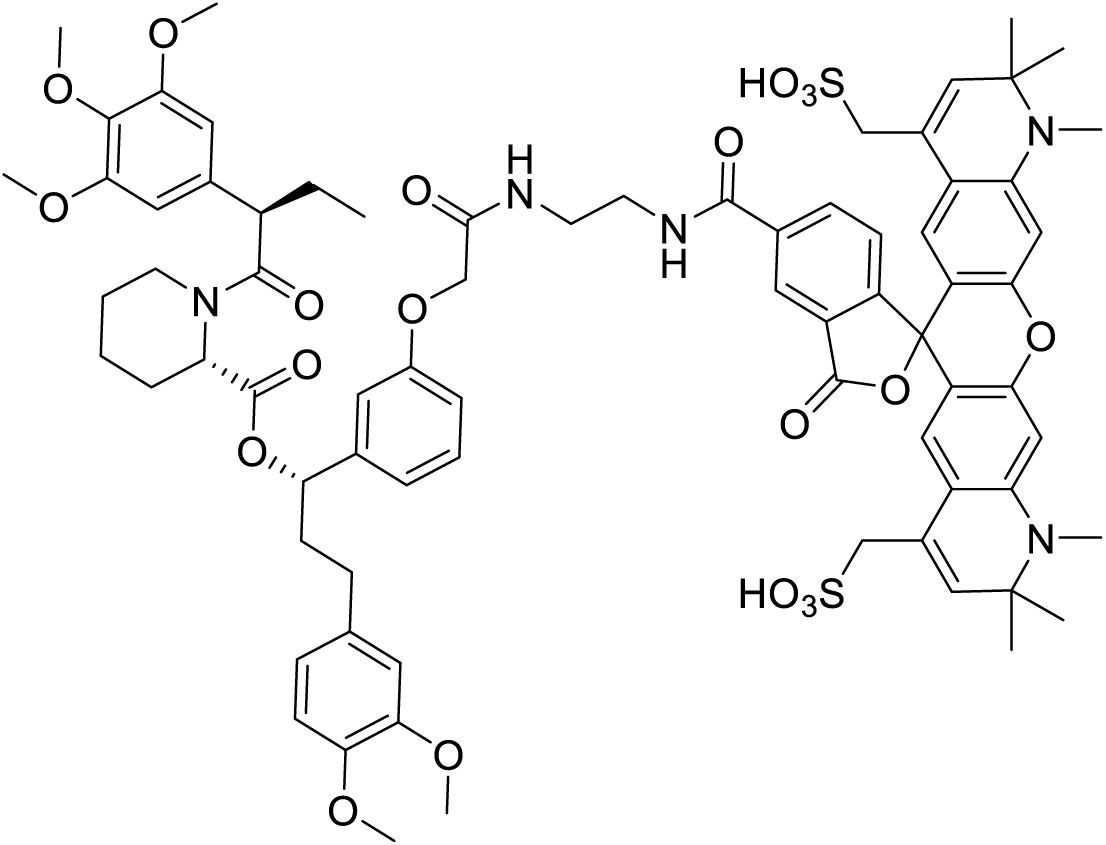

SLF’-AF594 was synthesized by strategy A, using AF594 NHS ester. The product was obtained as a purple solid.

^1^H NMR (500 MHz, DMSO-*d_6_*) δ 9.03 (t, *J* = 5.3 Hz, 1H), 8.52 (s, 1H), 8.43 (s, 2H), 8.20 (dd, *J* = 8.0, 1.5 Hz, 1H), 7.24 (d, *J* = 8.0 Hz, 1H), 7.18 (t, *J* = 7.9 Hz, 1H), 6.96 (d, *J* = 8.0 Hz, 1H), 6.91 – 6.82 (m, 4H), 6.73 (s, 2H), 6.64 (d, *J* = 8.4 Hz, 2H), 6.57 – 6.51 (m, 4H), 6.30 (s, 2H), 5.54 (s, 2H), 5.31 – 5.27 (m, 1H), 5.03 (s, 1H), 4.51 (s, 2H), 4.44 (d, *J* = 11.2 Hz, 2H), 4.02 (d, *J* = 8.9 Hz, 2H), 3.90 – 3.82 (m, 2H), 3.71 (d, *J* = 4.1 Hz, 6H), 3.64 (s, 1H), 3.60 (s, 6H), 3.57 (s, 3H), 3.17 (s, 6H), 2.68 – 2.61 (m, 2H), 2.40 – 2.32 (m, 2H), 2.17 (d, J = 11.0 Hz, 2H), 1.92 (ddd, J = 19.7, 11.4, 5.3 Hz, 4H), 1.57 (d, J = 6.9 Hz, 2H), 1.27 (d, J = 3.8 Hz, 12H), 0.88 – 0.83 (m, 3H).

^13^C NMR (126 MHz, DMSO-*d_6_*) δ 172.59, 170.75, 168.69, 166.72, 158.12, 153.27, 153.09, 152.21, 149.05, 147.46, 142.31, 136.38, 136.08, 135.99, 133.57, 129.95, 124.74, 123.26, 120.44, 119.19, 114.23, 113.40, 112.59, 112.30, 105.55, 97.11, 79.40, 79.14, 60.47, 60.30, 56.38, 56.00, 55.93, 55.78, 55.34, 53.31, 49.06, 31.41, 30.98, 29.46, 28.47, 28.20, 27.80, 22.56, 20.98, 12.73.

##### SLF’-HMSiR

SLF’-HMSiR was synthesized by strategy B, using HMSiR carboxylic acid. The product was obtained as a white solid.

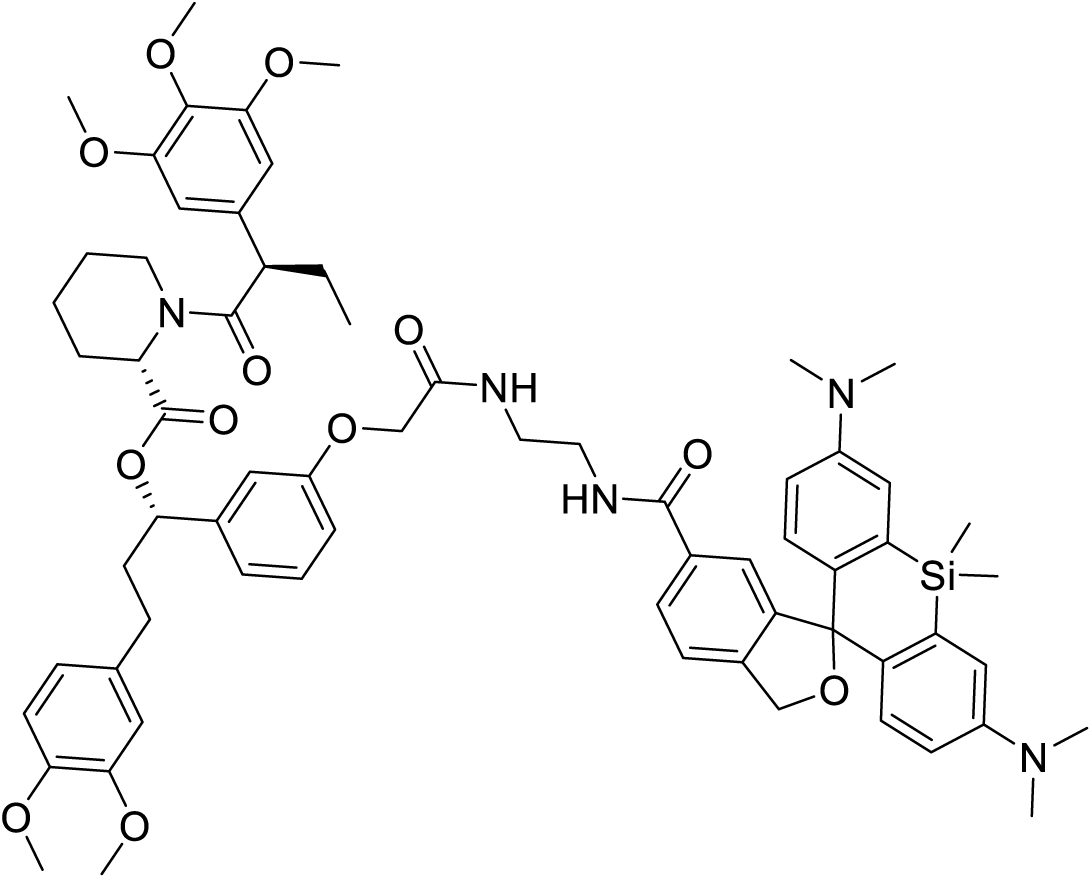

^1^H NMR (500 MHz, Chloroform-*d*) δ 7.64 (d, J = 7.9 Hz, 1H), 7.51 (s, 1H), 7.30 (d, J = 8.0 Hz, 2H), 7.19 (td, J = 7.8, 3.6 Hz, 1H), 7.11 (s, 1H), 7.05 (t, J = 7.8 Hz, 1H), 6.99 – 6.95 (m, 4H), 6.80 – 6.73 (m, 4H), 6.66 (d, J = 8.0 Hz, 4H), 6.62 – 6.59 (m, 2H), 6.41 (d, J = 6.5 Hz, 2H), 6.40 (s, 2H), 5.67 – 5.59 (m, 2H), 5.43 (d, J = 4.1 Hz, 2H), 5.29 (s, 2H), 5.12 (s, 2H), 4.49 – 4.44 (m, 4H), 3.86 – 3.84 (m, 15H), 3.78 (d, J = 8.0 Hz, 6H), 3.67 (s, 3H), 3.49 (d, J = 5.3 Hz, 2H), 3.35 (s, 2H), 2.93 (s, 12H), 2.80 (s, 1H), 2.55 (ddd, J = 14.4, 9.5, 5.3 Hz, 3H), 2.47 (ddt, J = 13.9, 7.3, 2.8 Hz, 2H), 2.30 (d, J = 11.1 Hz, 2H), 2.06 (tdt, J = 19.6, 10.1, 5.9 Hz, 8H), 1.97 – 1.91 (m, 2H), 0.89 (s, 3H), 0.67 (s, 3H), 0.54 (s, 3H).

^13^C NMR (126 MHz, Chloroform-*d*) δ 172.76, 170.58, 169.80, 167.77, 157.32, 153.17, 148.87, 148.79, 147.35, 136.64, 135.25, 133.56, 133.33, 129.74, 128.35, 126.00, 123.49, 121.26, 120.20, 120.10, 119.96, 116.78, 113.89, 112.56, 111.28, 105.01, 104.97, 104.60, 67.26, 60.80, 56.31, 55.97, 55.95, 55.92, 55.85, 52.13, 50.78, 40.80, 40.50, 39.50, 38.15, 31.93, 31.32, 29.70, 29.37, 29.32, 28.35, 28.33, 26.76, 22.70, 20.87, 14.13, 12.55.

HRMS [M+H]^+^: Calcd 1176.5724, found 1176.5693.

##### SLF’-JF525

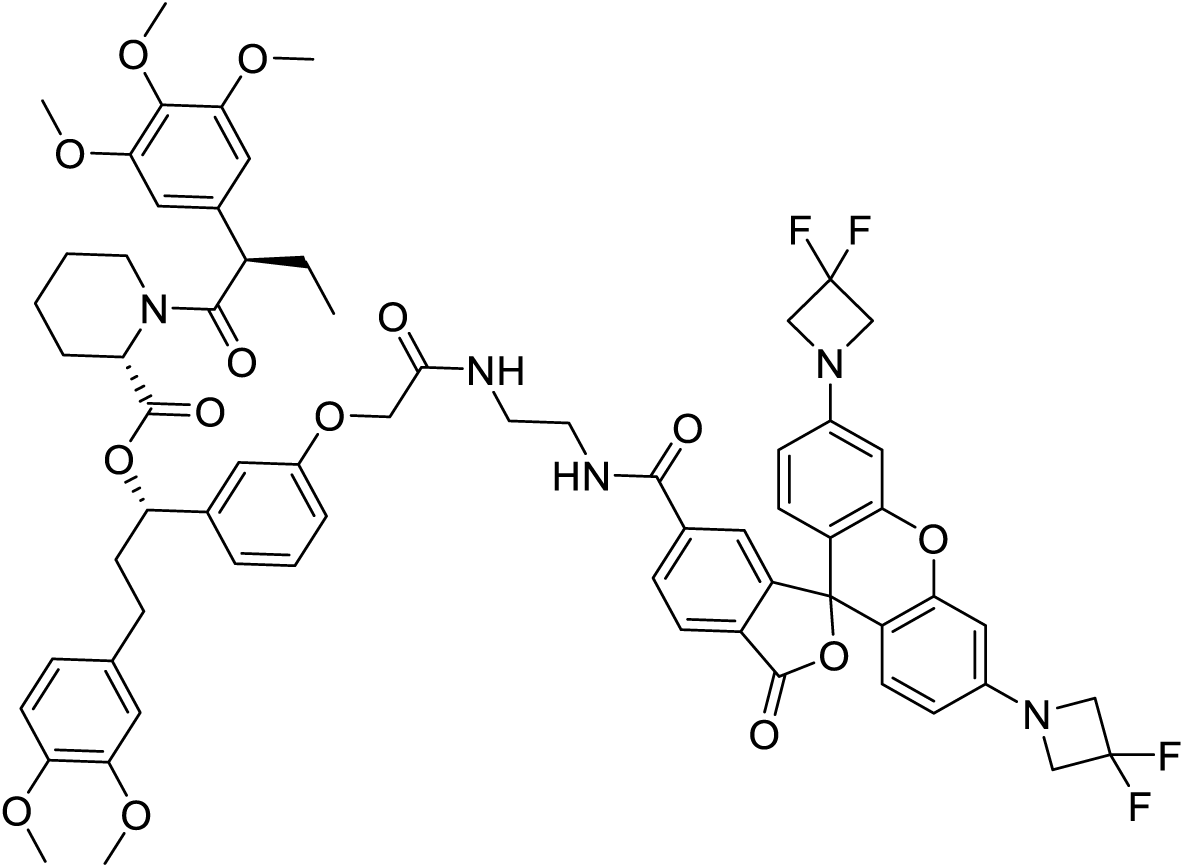

SLF’-JF525 was synthesized by strategy A, using JF525 NHS ester. The product was obtained as an orange solid.

^1^H NMR (500 MHz, Chloroform-*d*) δ 8.24 – 8.18 (m, 1H), 8.03 – 7.98 (m, 1H), 7.74 (d, J = 7.1 Hz, 1H), 7.16 – 7.06 (m, 2H), 6.71 (d, J = 7.7 Hz, 2H), 6.56 (dd, J = 24.3, 8.4 Hz, 5H), 6.34 (s, 2H), 6.24 (d, J = 2.2 Hz, 2H), 6.14 – 6.07 (m, 2H), 5.60 – 5.52 (m, 1H), 5.39 (d, J = 5.0 Hz, 1H), 4.42 – 4.37 (m, 2H), 4.19 (t, J = 11.6 Hz, 8H), 3.81 (s, 1H), 3.78 (d, J = 3.1 Hz, 6H), 3.77 (s, 3H), 3.69 – 3.65 (m, 6H), 3.52 – 3.49 (m, 2H), 2.72 (t, J = 11.1 Hz, 1H), 2.55 – 2.38 (m, 2H), 2.28 – 2.17 (m, 2H), 2.00 (dt, J = 14.6, 7.7 Hz, 4H), 1.86 (dd, J = 11.8, 5.8 Hz, 1H), 1.64 (dd, J = 13.5, 6.8 Hz, 4H), 0.83 (s, 3H).

^13^C NMR (126 MHz, Chloroform-*d*) δ 168.51, 153.31, 153.19, 152.42, 151.34, 148.89, 136.64, 135.33, 133.36, 130.88, 129.81, 129.49, 129.15, 125.28, 125.07, 120.20, 111.71, 111.28, 108.82, 108.76, 105.00, 104.60, 99.33, 81.90, 75.61, 71.34, 71.16, 70.70, 70.55, 68.89, 66.88, 64.51, 63.49, 63.28, 63.08, 60.80, 56.32, 55.98, 55.93, 55.85, 52.69, 52.07, 50.80, 43.49, 42.68, 39.51, 38.23, 31.94, 31.30, 29.71, 29.67,29.37, 28.37, 26.80, 25.87, 25.33, 22.70, 20.92, 14.13, 12.57, 9.19.

HRMS [M+H]^+^: Calcd 1244.4850, found 1244.4837.

## Notes

### Summary of Updates

Author list updated; Main text updated; Figures 1-7 revised; Supplemental files updated.

